# A BMP-to-GREM1 cancer-associated fibroblast switch at the colorectal cancer invasive front, revealed by histology-anchored spatial transcriptomics

**DOI:** 10.64898/2026.11.25.769142

**Authors:** Shamim Niwase, Jorge Pardo Romero, Akihisa Fujiyama, Gumpei Yoshimatsu

**Affiliations:** Division of Advanced Science, AI and Technology Innovation, OmiCore Inc., Fukuoka, Japan; Graduate School of Engineering, Kyushu University, Fukuoka, Japan; Tohoku University Graduate School of Medicine, Sendai, Japan

## Abstract

Which fibroblast state signals to tumour cells at the colorectal cancer invasive front, and from what distance, has not been resolved in human tissue, and spatial transcriptomics obscures the answer whenever tumour regions are defined from the expression being analysed and RNA is assigned to the wrong, neighbouring cell. We introduce a histology-anchored framework that segments cells from H&E (OmiCoreSeg), maps the tumour from H&E alone (OmiCoreTumorDetector; area under the ROC curve 0.91 against pathologist annotation) and returns RNA to the cell that made it (OmiCoreCloud). In three untreated stage II–IV colorectal adenocarcinomas profiled by Visium HD, Xenium and single-cell RNA-seq, with pre-registered tests carried to held-out patients, platforms and four bulk cohorts (1,319 patients), RNA spill reached only adjacent cells (8–12 µm), stromal programmes varied over *∼*400 µm, and expression-defined tumour regions overstated immune exclusion three- to fivefold. Across the H&E border, cancer-associated fibroblasts switch from a BMP4/5-producing, telocyte-like state in the tumour bed to a GREM1-producing palisade with TGF-*β*/SMAD3 and epithelial–mesenchymal transition (EMT) activity at the front (6/6 validation sections); adjacent tumour cells lose the BMP target ID1 (held-out odds ratio 0.47) and SMAD5 activity. No ligand–receptor pair was enriched at cell contact (0 of 1,233), whereas myeloid IL-1 and lymphoid LT*β* coupled to fibroblast programmes at niche scale. A GREM1-fibroblast score was associated with shorter survival (hazard ratio 1.23), and unlike the mouse and organoid GREM1–ISLR model the human axis is one-sided. These maps nominate GREM1 neutralisation and IL-1 blockade as front-targeted therapeutic hypotheses and provide a spatial reference of the human colorectal cancer front.

**Graphical summary:** **1, Measure.** RNA is anchored to histology with OmiCoreSeg (cells segmented from H&E), OmiCoreTumorDetector (a tumour map that never sees expression) and OmiCoreCloud (each cell’s own versus neighbouring-cell RNA). RNA spill reaches only adjacent cells (8–12 µm), whereas stromal programmes vary over *∼*400 µm, and RNA-defined tumour regions inflate immune exclusion three- to fivefold. **2, Discover.** Across the H&E tumour border, BMP4/5-producing fibroblasts beside ID1^+^ glands in the tumour bed give way to a GREM1-producing palisade (TGF-*β*/SMAD3, EMT) at the front, where tumour cells lose ID1 and SMAD5 activity; CD8 T cells collect at the margin, and myeloid IL-1 and lymphoid LT*β* couple to fibroblasts at niche scale rather than at cell contact. **3, Translate.** A GREM1-state score is associated with shorter survival (hazard ratio 1.23 per s.d.; 1,319 patients; not independent of molecular subtype); GREM1 neutralisation and IL-1/IL1R1 blockade are hypotheses for testing. Schematic; cell positions are illustrative.

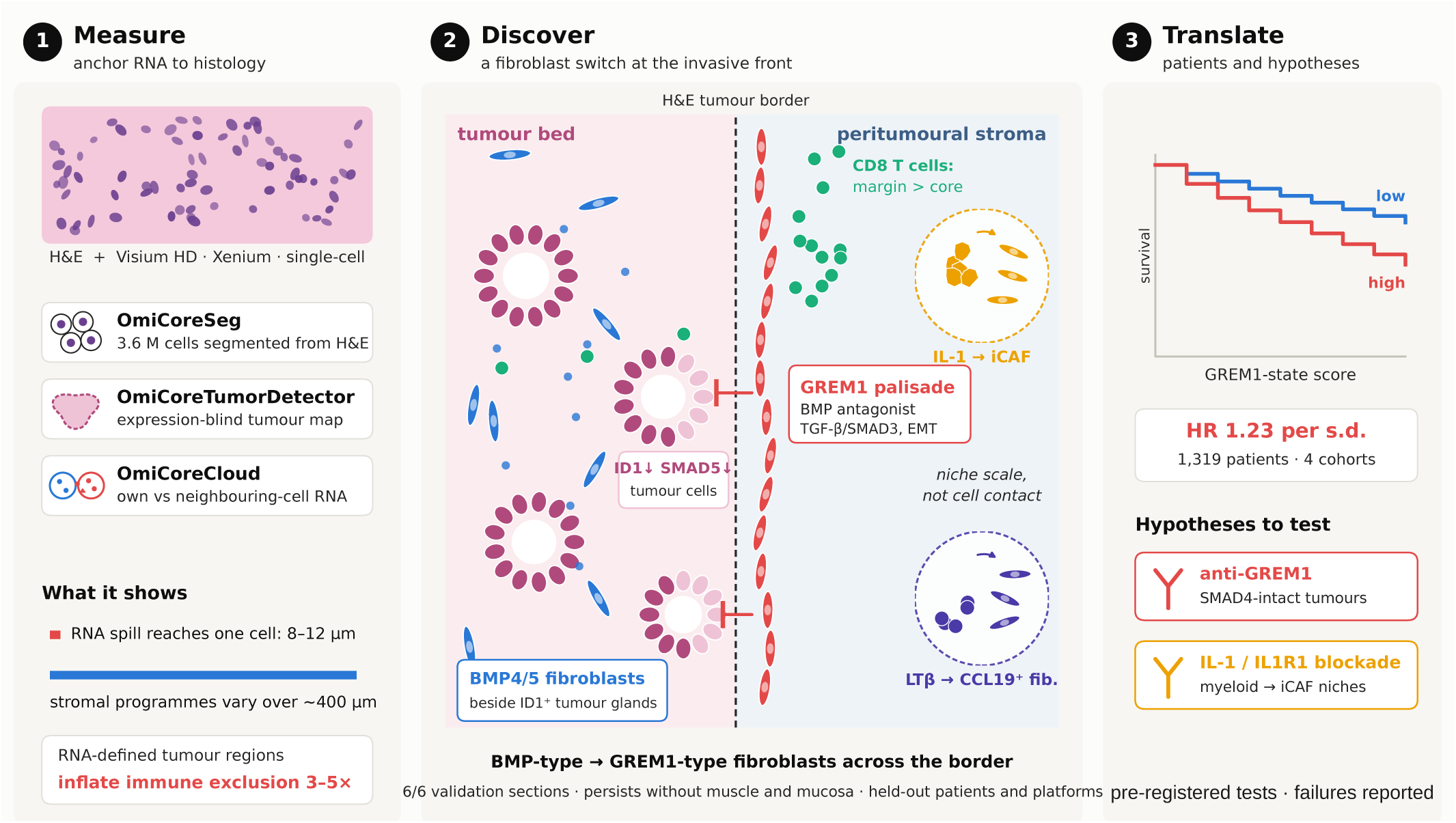

## Introduction

The invasive front of colorectal cancer (CRC) is where the tumour is graded, staged and, increasingly, measured. The type, density and position of immune cells at the front and in the core predict outcome ^1,2^. Stromal programmes define the poor-prognosis mesenchymal consensus molecular subtype CMS4^3,4^, and because stroma distorts bulk classification, tumour-intrinsic subtyping had to be developed separately ^5^. Single-cell and spatial atlases have mapped FAP^+^ fibroblast and SPP1^+^ macrophage niches, immune hubs and an organised tumour–stroma boundary ^6–10^. What these studies rarely resolve is *which* fibroblast state, at *which* position relative to the tumour, sends *which* signal to the tumour cells. A biomarker or a drug that acts on that signal needs exactly this information.

Bone morphogenetic protein (BMP) signalling is a strong candidate for such a signal. In the normal mouse colon, PDGFRA-high telocytes near the crypt top make BMP ligands, and GREM1-expressing trophocytes at the crypt base antagonise them, so that stemness is restricted to the crypt base ^11^. Ectopic epithelial GREM1 initiates colonic neoplasia ^12^. Kobayashi et al. ^13^ proposed that the stromal balance between GREM1 and the BMP potentiator ISLR drives colorectal carcinogenesis: GREM1^+^ and ISLR^+^ cancer-associated fibroblasts are distinct populations, TGF-*β* acting through FOXL1 drives *Grem1* at the expense of *Islr*, GREM1-high and ISLR-low tumours fare worse, and restoring BMP signalling—with a GREM1-neutralising antibody in Smad4-intact but not Smad4-mutant tumoroids, or with AAV8-delivered *Islr*—restrains tumour growth. The mechanism in that model was established in a mouse colonic fibroblast line, mouse organoids and a mouse liver-metastasis model; its human evidence is bulk, sorted-cell and single-cell expression profiles and in situ hybridisation. The direction of BMP effects is context-dependent, however: BMP4 and BMP–Notch crosstalk mark poor-prognosis mesenchymal CRC ^4^. The association of bulk GREM1 with outcome is established; its spatial and cellular organisation in human tumours is not. Mouse models differ from human colorectal cancer in ways that matter for the stroma: engineered tumours carry a uniform set of driver mutations and arise over weeks rather than decades, carcinogen and colitis models add inflammation, and the composition and function of the immune system differ between the two species ^14^. Whether the stromal states that organise the mouse colon and mouse tumours are the ones present at the human invasive front therefore has to be measured in human tissue.

Mapping that organisation from spatial transcriptomics raises three measurement problems. First, if the tumour is defined from the expression being analysed, compartment comparisons become partly circular. Second, at near-cellular resolution, RNA from neighbouring cells is assigned to the wrong cell, most strongly where cell types interdigitate, which is exactly at the tumour border ^15,16^. Third, zone contrasts are usually tested against gene- or label-permutation nulls that destroy spatial autocorrelation, so almost any partition of tissue looks significant. Cell–cell communication tools add a fourth problem: they infer signalling from the co-expression of a ligand and its receptor ^17–20^, which is not signalling and is the readout most exposed to spill.

Here we address these problems by construction, with three tools developed by OmiCore Inc. and a set of calibrated mathematical operators. OmiCoreSeg segments nuclei and cells from the H&E image. OmiCoreTumorDetector ^21^, which is open source, maps the tumour and eight other tissue classes from H&E and never sees expression. OmiCoreCloud attributes each cell’s RNA to the cell or to its neighbourhood and returns decontaminated and denoised profiles. On this footing we fitted an empirical spill kernel, computed signed distances, change surfaces, multiscale contrasts, curvature and persistent homology of the front, and read every zone claim against block-permutation nulls. Every finding from three discovery tumours was then re-tested, with rules written down before each run, on the same patients on a second platform, on new patients and in four bulk cohorts. We ask what each tool changes, what the front looks like when it is measured this way, and which candidate markers and therapeutic hypotheses survive. We report the pre-registered tests that failed alongside those that passed. The result is a cell-resolved, histology-anchored spatial atlas of the human colorectal cancer invasive front, with the level of evidence reached by each finding, and a measured test of which parts of the mouse-derived model of stromal BMP signalling hold in patients.

## Results

### Three histology-anchored measurements and a pre-registered validation design

Discovery used Visium HD sections of three untreated primary colorectal adenocarcinomas (P1, rectum, stage IIA; P2, sigmoid, IIIB; P5, IVA) and two normal-adjacent mucosae ^22^ (Fig. 1A; Supplementary Fig. 1). OmiCoreSeg segmented 3,565,452 cells on the five whole H&E sections; 792,159 lie in the capture areas and 674,823 were confidently typed by RCTD ^23^ against the patients’ Chromium Flex reference (260,506 cells, 38 level-2 states). On 23,550 expert-annotated PanNuke colon nuclei ^24^, OmiCoreSeg had the highest panoptic quality of nine configurations (PQ 0.538, 95% CI 0.526–0.550; InstanSeg 0.502, Cellpose-SAM 0.484, tuned StarDist 0.468; all comparators run on unmodified H&E), and it was higher than every comparator on 15 or 16 of 16 image mosaics (Fig. 1B; Supplementary Fig. 2; Supplementary Table 1). The gain comes from recall; its precision (0.757) was below that of InstanSeg (0.814) and Cellpose-SAM (0.812).

**Figure 1:**
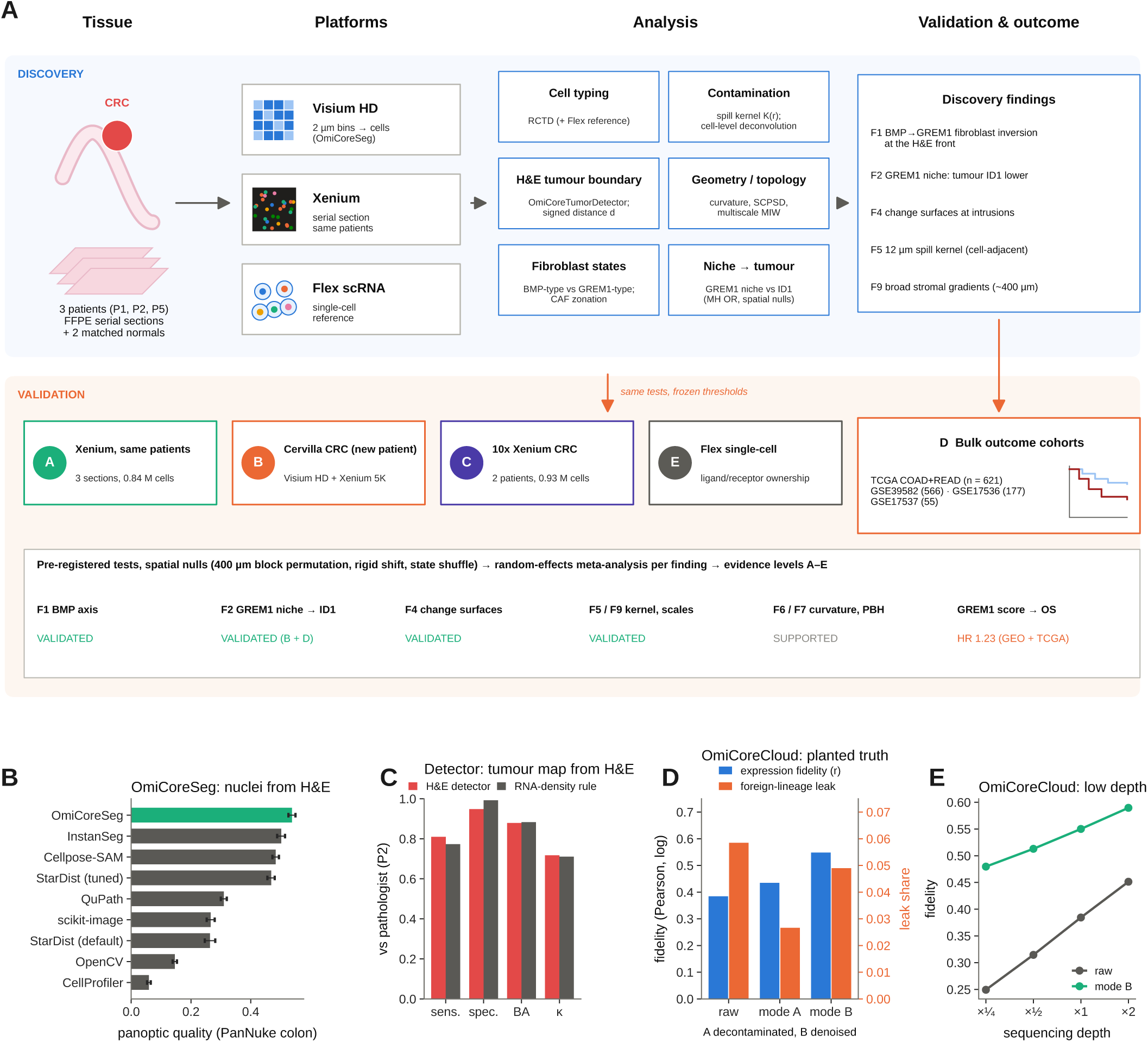
Study design and the three histology-anchored tools. **A**, Discovery on three colorectal cancers (Visium HD with OmiCoreSeg cells, Xenium serial sections, Flex single-cell reference); cell typing, spill kernel and cell-level deconvolution, the H&E tumour boundary (OmiCoreTumorDetector; signed distance *d*), geometry/topology and fibroblast-state analyses; validation of each finding with frozen tests at levels A (same patients, Xenium), B (new patient, Cervilla Visium HD and Xenium 5K), C (new patients, 10x Xenium), D (bulk TCGA and GEO) and E (Flex plausibility). Bottom strip, verdict per finding. **B**–**E**, What each tool measures. **B**, OmiCoreSeg panoptic quality on 23,550 PanNuke colon nuclei against open-source segmenters run on unmodified H&E (benchmark comparators; 95% bootstrap CI); OmiCoreSeg 0.538 [0.526–0.550]. **C**, OmiCoreTumorDetector versus an RNA-density rule against the pathologist annotation of section P2 (94,158 bins with typed cells): sensitivity 0.81 versus 0.77, specificity 0.95 versus 0.99, balanced accuracy 0.88 versus 0.88, *κ* 0.72 versus 0.71. **D**, OmiCoreCloud on planted truth (held-out patient on real P1 geometry, depth and contamination): expression fidelity and foreign-lineage leak of raw counts, mode A (decontaminated) and mode B (denoised). **E**, Mode B fidelity across sequencing depth (0.480 versus 0.249 raw at one quarter depth). OmiCoreSeg and OmiCoreCloud are shown by their inputs and outputs (see Methods and Code availability).

OmiCoreTumorDetector, trained only on public NCT-CRC-HE tiles ^25^, ranked RCTD tumour cells with an area under the ROC curve (AUROC) of 0.955, 0.854 and 0.952; its threshold was chosen on P1 alone. Against an independent pathologist annotation of section P2 it reached AUROC 0.911 (sensitivity 0.773, specificity 0.950, *κ* 0.697), and agreement fell monotonically as the map was shifted by 100–400 µm (Supplementary Fig. 3). In a head-to-head on the same bins, the detector and a conventional RNA-density rule had equal balanced accuracy (0.879 and 0.883); the detector was more sensitive and the RNA rule more specific (Fig. 1C). The detector is a compartment map, not a cell classifier: RNA identifies aneuploid tumour cells better (AUROC 0.81/0.97/1.00 against 0.75/0.94/1.00; aneuploid shares per class in Supplementary Fig. 4C). Its limit is the stain domain. On faded post-Xenium H&E from 10x Genomics it failed (AUROC 0.60 and 0.71), so every external section used an RNA tumour mask defined in advance. The detector scaled to 599 TCGA diagnostic slides (Supplementary Fig. 5).

OmiCoreCloud takes cell counts, positions, the reference atlas and an ambient prior as inputs and returns five outputs: a decontaminated profile (mode A), a denoised profile (mode B), a state cloud over reference states, its entropy and an atlas-misfit score (enthalpy). On planted truth, a held-out patient placed on real P1 geometry at real depth and contamination, mode B raised expression fidelity from 0.384 to 0.548 and mode A roughly halved foreign-lineage leak relative to raw counts; at one-quarter of the sequencing depth, mode B fidelity was 0.480 against 0.249 (Fig. 1D,E; Supplementary Fig. 6). It is not a better cell-state caller than simple correlation to the reference (balanced level-2 accuracy 0.46 versus 0.51), so cells were typed with RCTD.

Five validation levels were fixed before the validation runs: A, Xenium serial sections of the same patients (835,881 cells, 422 genes); B, one new patient ^26^ on Visium HD (8 µm bins) with the same patient’s Xenium 5K section (307,235 cells, 5,001 genes) as a platform replicate, not an extra level; C, two new patients on 10x Xenium; D, bulk TCGA COAD+READ (621 tumours) and three GEO cohorts ^27^; and E, single-cell plausibility in the Flex reference. A finding was VALIDATED when the validation-only random-effects pool excluded the null and at least two levels agreed (Supplementary Table 2).

### What histology anchoring changes

We next measured what anchoring to histology changes (Fig. 2).

**Figure 2:**
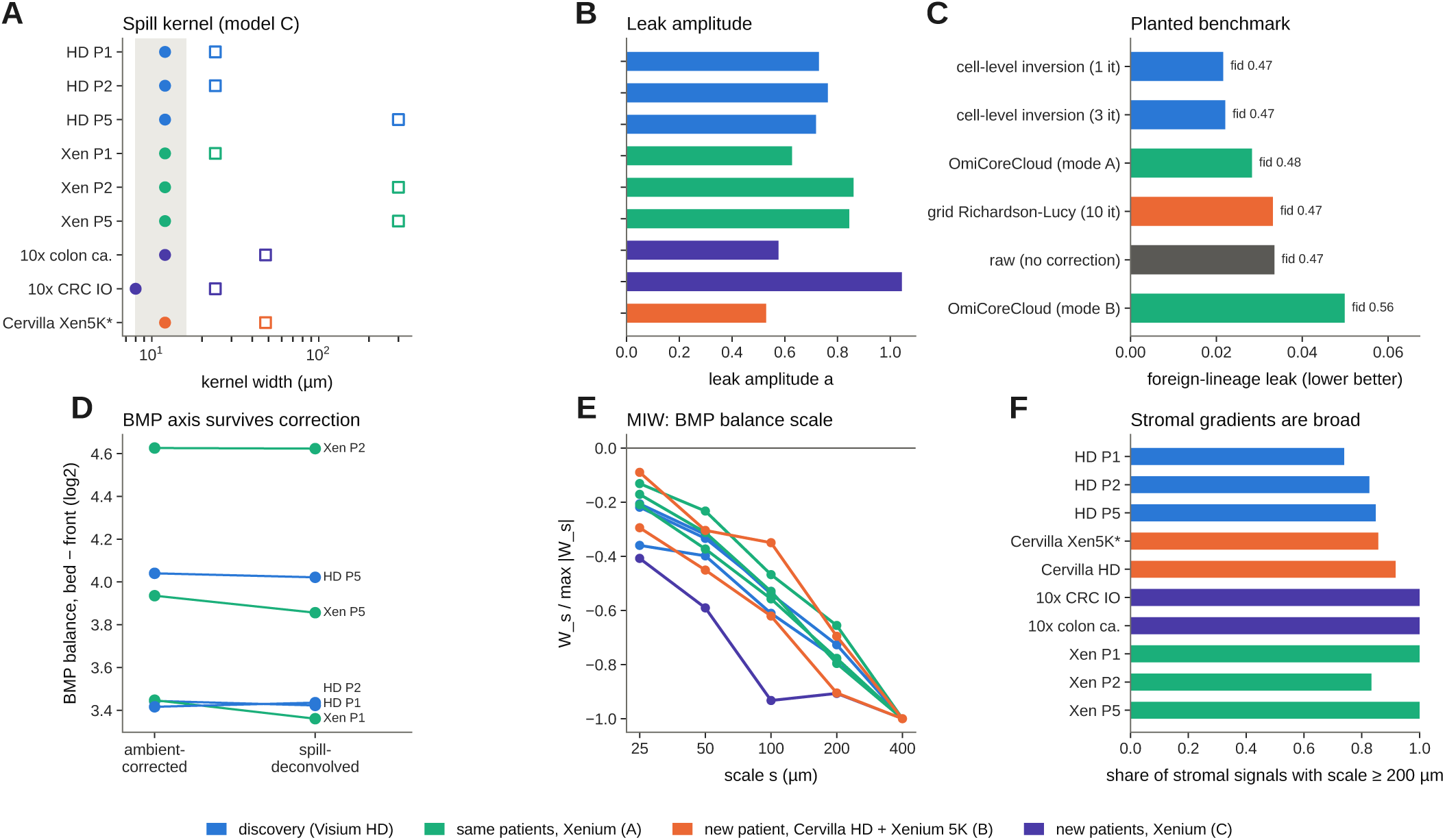
A cell-adjacent spill kernel and the scale of stromal programmes. **A**, Joint two-Gaussian kernel per section; *σ*_1_ (circles) is 8–12 µm on every platform (grey band, the pre-registered 8–16 µm range); *σ*_2_ (squares) is 24–48 µm or at the 300 µm grid edge. **B**, Relative leak amplitude (not a fraction). **C**, Planted-truth benchmark of corrections (foreign-lineage leak, fidelity annotated); OmiCoreCloud outputs labelled mode A/B. **D**, Fibroblast BMP balance (bed *−* front) before and after deconvolution (after deconvolution, P1/P2/P5: Visium HD +3.42/+3.44/+4.02, Xenium +3.36/+4.62/+3.86; raw values in the text; block nulls within *±*1.22). **E**, Multiscale interface wavelet (MIW) profile of the BMP balance across 25–400 µm. **F**, Share of stromal signals with characteristic scale *≥*200 µm. Colours, evidence level.

Two contributions were measured directly. The first concerns the tumour region. Against the H&E map, immune cells were 9.0-, 3.6- and 5.1-fold less frequent inside the tumour than beyond 50 µm outside it; 20 block-permuted maps gave 0.7–1.7-fold, and a whole-section map (Supplementary Fig. 7) gave 9.0, 3.5 and 4.8. With an expression-defined tumour (*≥*60% of the 20 nearest typed cells called tumour), the same cells gave 25.6-, 18.4- and 20.8-fold. The density rule carves real immune pockets and stromal septa out of the tumour: it kept 91–98% of intratumoural immune cells inside when they were placed at random tumour positions, but only 16–30% at their real positions (Supplementary Fig. 4). Expression-defined regions thus inflate exclusion three- to fivefold, and the detector region retained three to six times more intratumoural immune cells (three to eight times more purity-filtered macrophages). The region also changed biology. A Th17 programme that was not detected under RNA-defined regions was detected under the H&E map (IL17A +3.68, IL17F +2.84 log_2_, 3/3 tumours; owned-gene *q* 0.008 and 0.037), and an apparent loss of CCR7 in intratumoural T cells (*−*5.28) shrank to *−*0.13 under the H&E map, so we attribute it to the region definition.

The second concerns spill. For each lineage we fitted a two-Gaussian kernel to genes that only that lineage expresses, observed in cells of other types (Methods). The short component *σ*_1_ was 12 µm in all six discovery sections on both platforms and 8–12 µm in every external section, and shifting the source by 1 mm removed 82–100% of the likelihood gain (Fig. 2A,B). The pre-registered expectation that Xenium would leak much less than Visium HD failed: leak amplitudes were similar, because spill is neighbour mis-assignment shared by segmented platforms, not a diffuse 100 µm background. A cell-level inversion with this kernel removed 1.7–6.9% of counts and cut planted foreign-lineage leak by 36% at unchanged fidelity, whereas a pre-registered grid deconvolution failed its held-out check (Fig. 2C). Every headline survived the deconvolved matrix with pass rules fixed in advance (Fig. 2D). Multiscale interface wavelet (MIW) analysis agreed: the fibroblast BMP balance and 80–100% of CAF genes per section had a characteristic scale of 400 µm, replicated in new patients (Fig. 2E,F). Contamination acts at one cell diameter; stromal zonation acts over hundreds of micrometres. One claim, interferon/MHC-II induction at tumour-state fronts, was not supported, because the immune-cell markers CD3E and HLA-DRA rose with it, as expected from spill rather than induction.

### An H&E-anchored pathology atlas of the invasive front

The detector’s nine-class output maps the bowel wall (Fig. 3A–F). Against the P2 pathologist annotation, the classes agreed with *κ* = 0.67; neoplasm (recall 0.80, precision 0.95) and non-neoplastic epithelium (0.93) were recovered well, smooth muscle conservatively (0.38) and connective tissue poorly (0.27). We use these classes as an anatomical covariate, not as a diagnosis.

**Figure 3:**
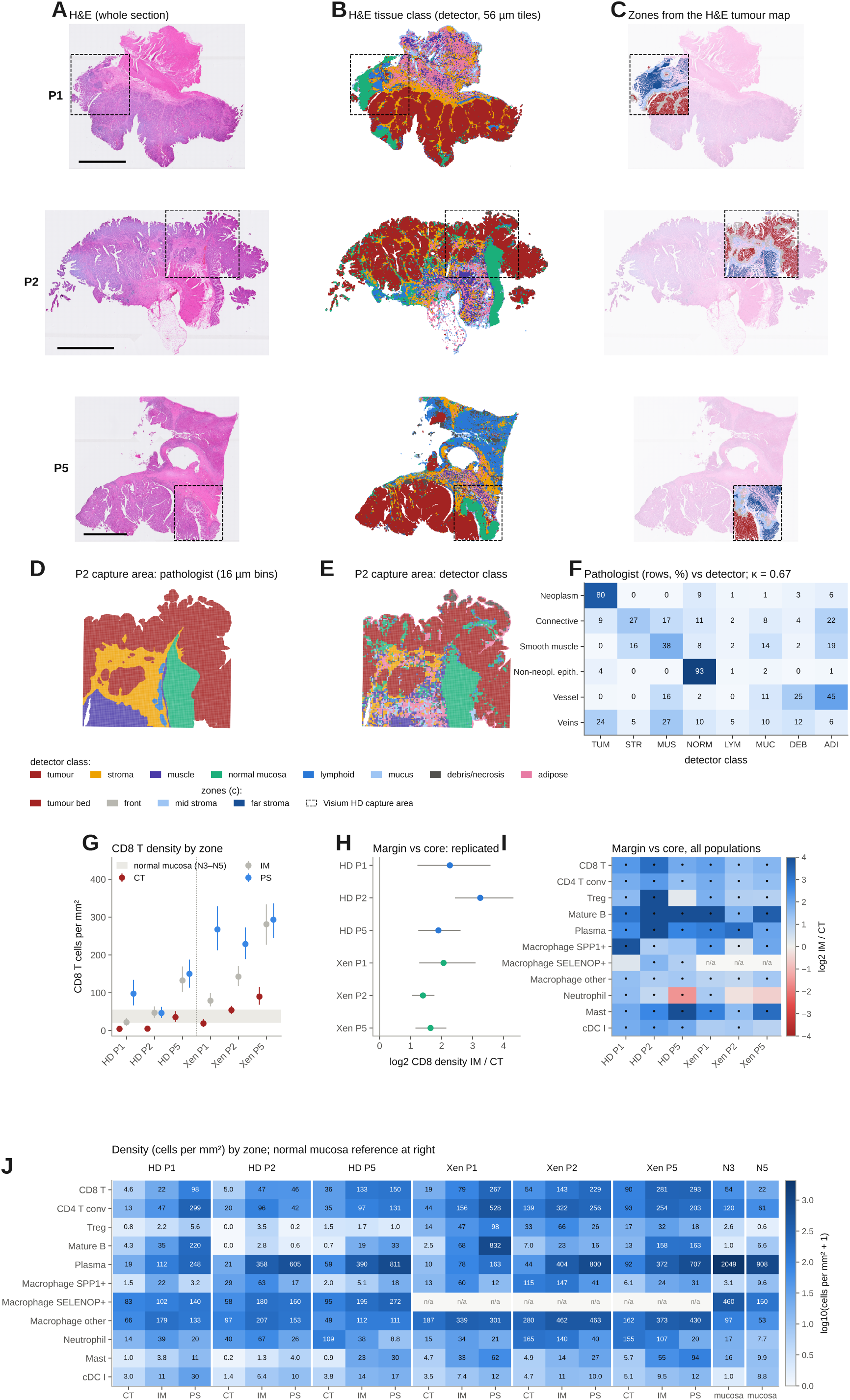
An H&E-anchored pathology atlas of the front. **A**–**F**, Wall anatomy from H&E alone. **A**, Whole-section H&E of P1, P2 and P5 (dashed box, capture area). **B**, Detector tissue class (arg-max of nine NCT-CRC-HE classes; 56 µm stride). **C**, Zones from the signed distance (bed *< −*50 µm; front *−*50 to 150 µm; mid 150–500 µm; far *>* 500 µm). **D**–**F**, Check against the pathologist annotation of P2 (16 µm bins): labels (**D**), detector class (**E**) and row-normalised confusion matrix (**F**); *κ* 0.67 over four shared classes (111,314 bins); neoplasm recall 0.80 (precision 0.95), non-neoplastic epithelium 0.93, smooth muscle 0.38, connective tissue 0.27. Scale bars, 5 mm. **G**–**J**, Immunoscore-type readout. Zones: CT, *d < −*250 µm; IM, *−*250 to +250 µm; PS, 250–1,000 µm; cells per mm^2^ of occupied tissue on a 25 µm grid. **G**, CD8 density by zone on Visium HD and Xenium with 400 µm block-bootstrap 95% intervals; grey band, normal mucosa. **H**, log_2_ CD8 IM/CT; pre-registered test met in 6/6 (HD 2.27/3.25/1.90; Xenium 2.06/1.40/1.64). **I**, log_2_ IM/CT for all populations; dot, interval excludes 0. **J**, Density per population, section and zone, with normal mucosa at right.

#### The tumours

Tumour cells kept MLH1, MSH2, MSH6 and PMS2 at 0.68–1.96 times the level of normal epithelium, far above a pre-set silencing threshold of 0.25, and CD8A transcripts were low. All three tumours are therefore most consistent with mismatch-repair-proficient (pMMR) tumours, an inference rather than a diagnosis. Each has a chromosomal-instability karyotype with 20q gain, and none has 18q (SMAD4) loss (Supplementary Fig. 1). All three were stroma-low by the Mesker criterion ^28^ (tumour–stroma ratio 4.9–8.4% by area; 14.7–19.0% with a TCGA-style definition, the 31st–40th percentile of 589 TCGA patients), with minor necrosis (1.7–2.9%) and mucin (0.4–1.4%) (Supplementary Fig. 8).

#### Immune organisation

We rebuilt the Immunoscore-type comparison of the tumour core (CT) and invasive margin (IM) ^2^ against the H&E border: CT more than 250 µm inside, IM a 500 µm band across the border (Fig. 3G–J). The pre-registered test, CD8 density IM *>* CT on both platforms, replicated in 6/6 sections with every interval excluding zero: log_2_ IM/CT 2.27, 3.25 and 1.90 on Visium HD and 2.06, 1.40 and 1.64 on Xenium, a 2.6- to 9.5-fold margin enrichment. The P1 and P2 cores held about 5 CD8 cells per mm^2^, fewer than normal mucosa (22–54). CD4, B, plasma and mast cells, dendritic cells and SPP1^+^ macrophages were also margin-enriched; neutrophils were the exception, dense in the P5 core (109–155 per mm^2^). Strictly defined NK cells were excluded from the tumour bed on both platforms (log_2_ bed/front *−*3.1, *−*1.1, *−*2.5 on Visium HD), and SPP1^+^ macrophages were enriched in the bed (+2.1 and +2.8 versus far stroma) while resident LYVE1^+^ macrophages were depleted. C1Q^+^ macrophage zonation was inconsistent between sections, so we make no claim about it, and an apparent loss of macrophage SPI1 activity near the tumour failed the block null and was withdrawn. A Crohn-like lymphoid reaction surrounded every tumour: 32, 44 and 48 aggregates lay within 2 mm of the border and few inside; 9–15% were secondary follicle-like, and tumour-associated aggregates carried CCL19 and LAMP3 (Supplementary Fig. 8).

#### Tumour-cell states

A pre-registered test that a fetal/revival programme is more frequent at the front (*−*100 to 0 µm) than in the core (*< −*300 µm) met its rule in 6/6 sections, but the effect was small (+0.1 to +3.5 percentage points) and its interval excluded zero in only three (Supplementary Fig. 9). Partial-EMT programmes were higher at the front and proliferating tumour cells less frequent (descriptive; an earlier pre-registered test of proliferation at tumour-state fronts failed on an independent patient, odds ratio 1.34 in the opposite direction).

These tissue phenotypes are robust measurements, but none stratified outcome when reduced to an H&E-only index. The pre-registered TCGA immune-exclusion index was not prognostic (progression-free interval hazard ratio 0.93, 0.78–1.11; overall survival 0.97, 0.79–1.19), and no H&E map feature passed correction (Supplementary Fig. 5). H&E-measured exclusion is therefore a measurement-validity result, not a biomarker.

### Geometry and topology locate abrupt tumour-state changes at stromal intrusions

Is the tumour interior homogeneous, and does the shape of the front matter? Averages along the signed distance treat the border as a straight line and the tumour as uniform; they cannot see where the border folds, where stroma intrudes or where the tumour state changes abruptly. We therefore used four operators, each computed from the H&E map or from the tumour-state field without reference to the gene being tested (Fig. 4A; Methods): the signed distance *d* (equation 3), which gives every cell a coordinate relative to the H&E border; total-variation change surfaces (spatial change-point surface detection, SCPSD; equation 11), which locate abrupt changes of tumour state; the curvature *κ* of the front (equation 13), which separates convex from concave stretches; and persistent homology of the tumour map (equation 14), which counts tumour islands and enclosed stromal pockets. For SCPSD, we rasterised tumour-cell state (ten principal components of tumour-owned genes, 16 µm pixels), denoised it by total variation, which preserves edges, and defined change surfaces as the top decile of the denoised gradient (Methods). Surfaces were stable over a fourfold range of the regularisation (Jaccard 0.84–0.97) and concentrated where stroma intrudes into the tumour (Fig. 4B): odds ratios 9.61, 9.92 and 7.23 on Visium HD and 4.43, 2.04 and 5.88 on Xenium, against at most 1.89 for 1 mm rigid shifts of the map and 1.2–2.1 for a state-shuffle control. The surfaces replicated in every new section (10x colon cancer 4.88; 10x CRC immuno-oncology 3.19; Cervilla Visium HD 32.7, inflated by 8 µm bin mixing; Cervilla Xenium 5K 3.19). By the pre-registered sign test the finding is VALIDATED: 7/7 validation sections (*P* = 0.0078) and 10/10 overall (*P* = 0.00098). The Xenium odds ratios (2.0–5.9) give the more conservative estimate of effect size.

**Figure 4:**
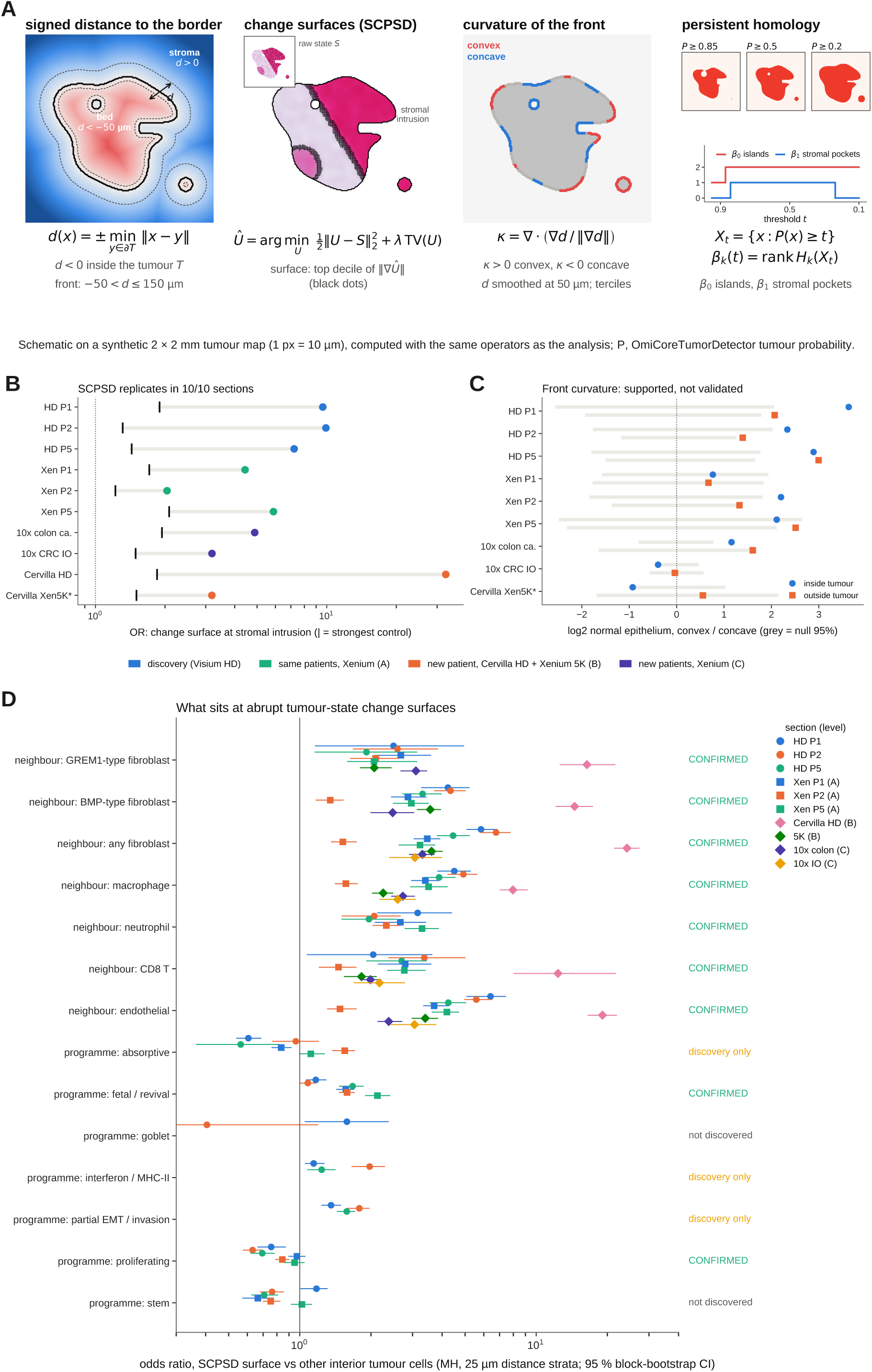
Geometry and topology of the front. **A**, The four operators, drawn on a synthetic 2 mm tumour map with the same code as the analysis. Signed distance (equation 3): *d* to the H&E border, negative inside the tumour; the front is *−*50 *< d ≤* 150 µm. Change surfaces (SCPSD; equations 11, 12): the tumour-state field *S* is denoised by total variation and the surface is the top decile of ||*∇Û*||. Curvature (equation 13): *κ* of the smoothed border is positive where the tumour bulges into stroma (convex) and negative where stroma bulges in (concave); terciles per section. Persistent homology (equation 14): superlevel sets *X_t_*= *{P ≥ t}* of the tumour probability *P*; the Betti numbers *β*_0_ and *β*_1_ count tumour islands and enclosed stromal pockets across thresholds. **B**, Odds ratio of total-variation change surfaces (SCPSD) at stromal intrusions in ten sections (discovery and levels A, B, C); tick, strongest control (maximum of 1 mm rigid-shift and state-shuffle controls). Validation 7/7 (sign test *P* = 0.0078); 10/10 overall. Cervilla Visium HD is inflated by 8 µm bin mixing. **C**, Normal epithelium at convex versus concave stretches of the front (log_2_), inside and outside the tumour; grey, 95% block-permuted curvature null; SUPPORTED, not validated. **D**, What sits at SCPSD surfaces (pre-registered). Odds ratios (surface versus non-surface interior tumour cells, stratified by 25 µm distance bins) for neighbour types within 30 µm (BMP-type, GREM1-type and all fibroblasts, CD8 T cells, macrophages, neutrophils, endothelium) and for tumour programmes, per section and platform, with block-bootstrap 95% CI; pre-stated hypothesis OR_GREM1_ *>* OR_BMP_. Circles, Visium HD discovery; squares, Oliveira Xenium (A); diamonds, external sections (B, C; RNA-mask distance, level-1 neighbour types). Right: verdict (CONFIRMED = 3/3 HD with CI excluding 1 in *≥*2/3, plus Xenium 3/3 and all measurable external sections). The hypothesis OR_GREM1_ *>* OR_BMP_ fails (HD 2.5/2.6/1.9 versus 4.2/4.3/3.3). Xenium programmes use platform markers (stated deviation).

What sits at these surfaces was tested with rules fixed before the validation runs (Fig. 4D). Surface and non-surface interior tumour cells were compared, stratified by distance, for neighbouring cell types within 30 µm and for tumour programmes, with the stated hypothesis that GREM1-type fibroblasts are enriched at surfaces relative to BMP-type fibroblasts. Every neighbour type was enriched at the surfaces and the enrichment was confirmed on Xenium 3/3 and on all measurable external sections: endothelium (HD odds ratios 6.4, 5.6 and 4.3), fibroblasts (5.8, 6.8, 4.5), macrophages (4.5, 4.9, 3.9), neutrophils (3.2, 2.1, 2.0) and CD8 T cells (2.0, 3.4, 2.7). Because surfaces were already known to sit at stromal intrusions, these enrichments describe what the intrusions are—vascular, fibroblast and macrophage septa that carry CD8 T cells—rather than a separate discovery. The pre-stated hypothesis failed: BMP-type fibroblasts were more enriched than GREM1-type fibroblasts in 3/3 HD sections (4.2, 4.3, 3.3 versus 2.5, 2.6, 1.9) and on Xenium 5K (3.6 versus 2.1), with mixed results elsewhere. Abrupt state changes inside the tumour therefore mark bed septa, not the GREM1 palisade of the outer front. The tumour cells on the surfaces were more often fetal/revival-like (HD 1.2, 1.1, 1.7; Xenium markers 1.6, 1.6, 2.1) and less often proliferating (HD 0.76, 0.63, 0.69; Xenium 0.97, 0.84, 0.95), both confirmed across platforms; absorptive, partial-EMT and interferon programmes were discovery-only, and stem and goblet programmes were not enriched.

The shape of the front carried weaker information. The H&E front was tortuous (boundary length 4.2, 5.8 and 4.1 times the hull perimeter), and normal epithelium was enriched at convex stretches in all discovery sections (log_2_ +2.3 to +3.6 inside the tumour), an expansile pattern towards mucosa (Fig. 4C). In validation the direction held in 4/7 and 5/7 sections, but the pooled effects were not significant (inside +0.34 [*−*0.65, 1.32]; outside +0.83 [*−*0.09, 1.75]): SUPPORTED, not validated. Pre-registered links between front shape and CAF state, GREM1 or tumour budding failed. Exact cubical persistent homology of the tumour map counted 157, 278 and 229 tumour islands and 169, 260 and 145 enclosed stromal pockets, in which CD4 T cells were enriched (log_2_ +0.82 and +1.64; Supplementary Fig. 10). Barrier and access metrics that worked in simulation, including geodesic and anisotropic transport, collapsed to cell density on tissue (Supplementary Fig. 11).

### Fibroblasts switch from a BMP-producing to a GREM1-producing state across the H&E border

Fibroblast genes that changed between zones pointed to the BMP pathway: BMP4, BMP5, WNT4, F3 and CXCL14 fell from the tumour bed outwards, while the antagonist GREM1, with SFRP2, rose towards the border. We therefore pre-specified a fibroblast BMP balance, log_2_[(BMP4 +BMP5 + 0.5)/(GREM1 + 0.5)] in fibroblast pseudobulks, and contrasted the tumour bed (*d < −*50 µm) with the front (*−*50 *< d ≤* 150 µm). In discovery the bed-minus-front contrast of the balance was +3.44, +3.42 and +4.04 log_2_ units (block nulls within *±*1.22), and +3.45, +4.63 and +3.94 on the Xenium serial sections; the spill-deconvolved values were almost identical (Fig. 5A). The balance reflects discrete states. BMP-type fibroblasts (*≥*1 BMP4/BMP5 and no GREM1 transcript) made up 15–18% of bed fibroblasts but 3–8% at the front; GREM1-type fibroblasts made up 1–6% of the bed and 16–34% of the front (Xenium 50–79%; panel b; profiles along the signed distance in Supplementary Fig. 12). Contamination cannot produce this pattern: BMP5, which tumour cells do not express, was 1.0–1.6 transcripts per 10,000 in bed fibroblasts but 0.14–0.45 at the front.

**Figure 5:**
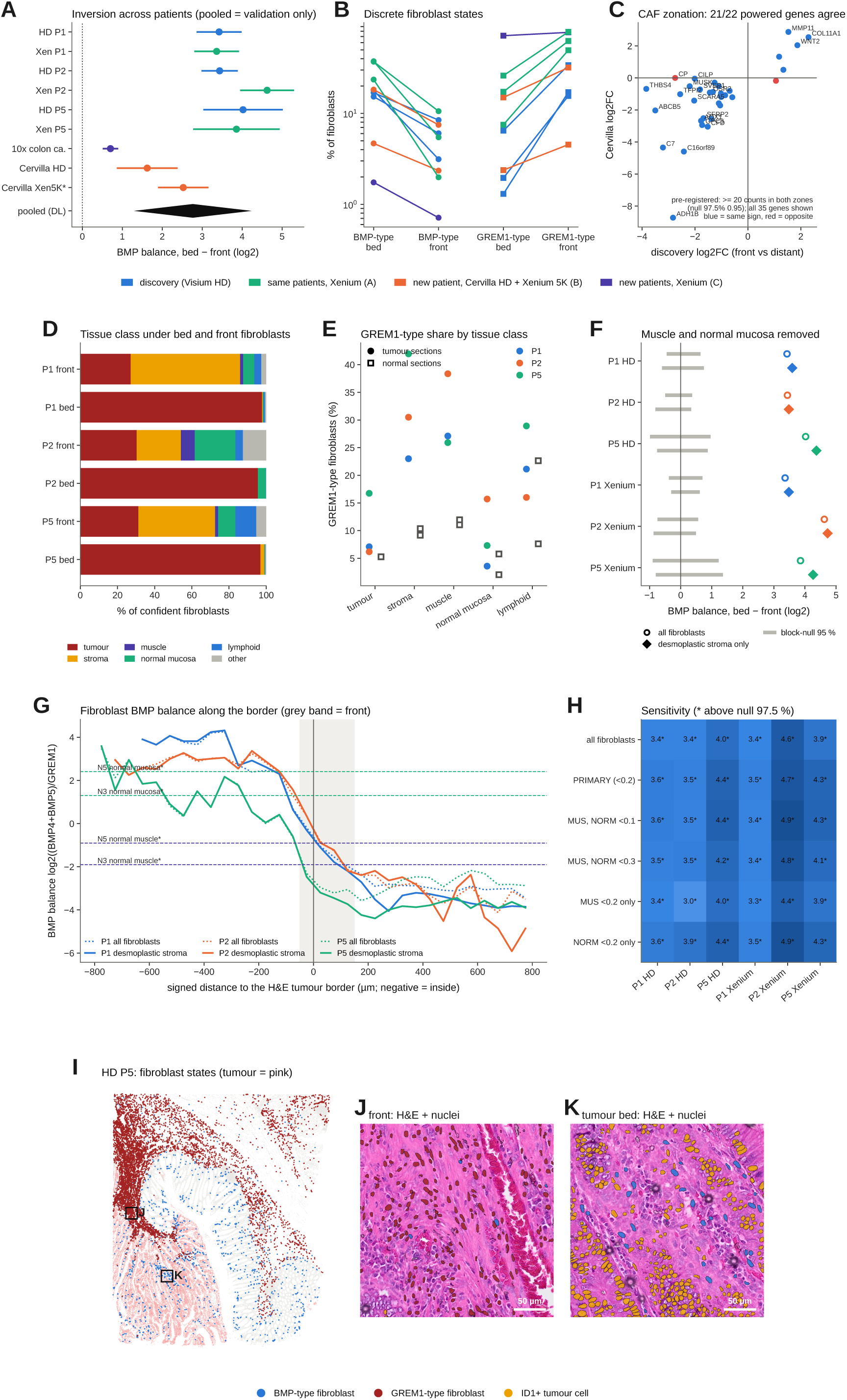
The fibroblast BMP*→*GREM1 switch. **A**, BMP balance contrast (bed *−* front, log_2_) per section with 95% interval; diamond, DerSimonian–Laird pool of validation sections only (+2.76 [1.30, 4.23], 6 sections); * Cervilla Xenium 5K, a platform replicate of the level-B patient. **B**, BMP-type (BMP4/5^+^ GREM1*^−^*) and GREM1-type (GREM1^+^ BMP4/5*^−^*) fibroblasts, % of fibroblasts in bed versus front; every section inverts. **C**, CAF zonation genes, discovery log_2_FC (near versus distant stroma) against the held-out Cervilla patient; the pre-registered test counts the 22 genes with *≥*20 counts in both zones (21/22 agree; null 97.5% 0.947); all 35 discovery genes are plotted. **D**–**H**, Pre-registered wall-anatomy control. **D**, Detector class under bed and front fibroblasts. **E**, GREM1-type share of fibroblasts by tissue class in tumour (circles) and normal-adjacent (squares) sections. **F**, Bed *−* front contrast for all fibroblasts (open) and after removing fibroblasts in muscle or normal-mucosa tiles (filled); grey, 95% range of 20 block-permuted maps. **G**, Balance along the signed distance (dotted, all fibroblasts; solid, desmoplastic stroma only); dashed, normal mucosa and muscle. **H**, Sensitivity to thresholds and exclusions; * above the null 97.5%. **I**–**K**, Fibroblast states in situ. **I**, Whole Visium HD section P5, BMP-type (blue) and GREM1-type (dark red) fibroblasts, tumour cells pink; boxes J and K mark the two zooms. **J**, H&E at the front with OmiCoreSeg nuclei coloured by state, showing a GREM1-type palisade. **K**, Tumour bed with BMP-type fibroblasts between glands and ID1^+^ tumour nuclei (yellow).

The pre-registered external tests replicated the switch: Cervilla Visium HD +1.62 (null 97.5% quantile 0.43), Cervilla Xenium 5K +2.53 (null 0.62; BMP-type 18.3% versus 7.5%, GREM1-type 15.0% versus 31.9%) and the 10x colon-cancer section +0.70 (null 0.16), about fivefold weaker than in discovery, with GREM1-type fibroblasts dominant at every depth. The validation-only pool was +2.76 [1.30, 4.23], positive in 6/6 sections (Supplementary Fig. 13). The finding is VALIDATED (levels A, B and C), universal in direction and variable in size (*I*^2^ = 0.98; prediction interval *−*2.64 to 8.17). The peritumoural CAF gene set that accompanies it (COL11A1, MMP11, INHBA, WNT2 and WNT5A up; THBS4, C7, ADH1B, SCARA5 and others down at the edge) also replicated: 21 of 22 testable genes kept their sign in the Cervilla patient (0.955; null 97.5% quantile 0.947; Supplementary Table 3), and 11 of 12 on Xenium 5K (0.917; null 0.877; panel c).

#### The switch is not wall anatomy

In the normal colon, GREM1 is made by smooth muscle and, as characterised in the mouse, by crypt-base trophocytes ^11^, so a front that cuts through muscle could mimic the switch. In a pre-registered control, each fibroblast took the detector tissue class of its position, and fibroblasts in tiles with muscle or normal-mucosa probability *≥*0.2 were removed (Fig. 5D–H). Front fibroblasts lay in muscle tiles at only 1.6–7.4%. The restricted contrast was +3.59, +3.48 and +4.37 on Visium HD (null 97.5% quantiles 0.75, 0.34 and 0.87) and +3.48, +4.73 and +4.26 on Xenium, and every sensitivity variant replicated (+3.0 to +4.9). GREM1-type fibroblasts made up 23–42% of tumour-section stroma tiles against 9–10% of normal stroma and 11–12% of normal muscle. The front balance of P1 and P2 (*−*1.20, *−*1.03) lies within the range of normal muscle; only mid and far peritumoural stroma (*−*2.7 to *−*3.3) goes beyond every normal compartment. Within the same tissue class, the GREM1 state is a tumour-associated state, but the detector’s muscle class is conservative, so a residual muscle contribution cannot be excluded entirely.

On H&E, GREM1-type fibroblasts lined the tumour front, and BMP-type fibroblasts threaded between tumour glands in the bed (Fig. 5I–K). Within 30 µm of front tumour cells, GREM1-type fibroblasts were more depleted than BMP-type fibroblasts (log_2_ observed/expected *−*3.7 versus *−*1.4): a stromal palisade set back from the tumour, which fits a secreted antagonist acting over tens of micrometres. The two states were segregated (outward BMP-type*→*GREM1-type neighbour transitions depleted 6/6, observed/expected 0.24–0.74) with no double-positive intermediate; a pre-registered directional test failed, so we make no claim that one state converts into the other.

### Two fibroblast programmes and a tumour BMP-response deficit

#### Fibroblast-state transcriptomes

To ask what else distinguishes the two states, we compared their whole transcriptomes on the spill-deconvolved matrix (pseudobulk per section and state, limma-voom ^29,30^ with section blocking; defining genes excluded; genes owned by other lineages removed). Of 11,959 genes, 309 met the discovery rule (*q <* 0.05, *|* log_2_ FC*| ≥* 1, same sign in 3/3 sections; Supplementary Table 4): 176 higher in GREM1-type and 133 higher in BMP-type fibroblasts (Fig. 6D). BMP-type fibroblasts were marked by F3, TRPA1, SPRY4, WNT4 (*−*5.0 log_2_), SALL1, WNT5A, EDNRB and CXCL14, a telocyte-like, PDGFRA-associated programme that OmiCoreCloud had flagged earlier as poorly explained by the reference atlas (supportive only; novelty AUROC about 0.59). GREM1-type fibroblasts were marked by SFRP2, SFRP1, OGN, MGP, MEDAG, TNXB, FGF7, PTGIS and CTHRC1.

**Figure 6:**
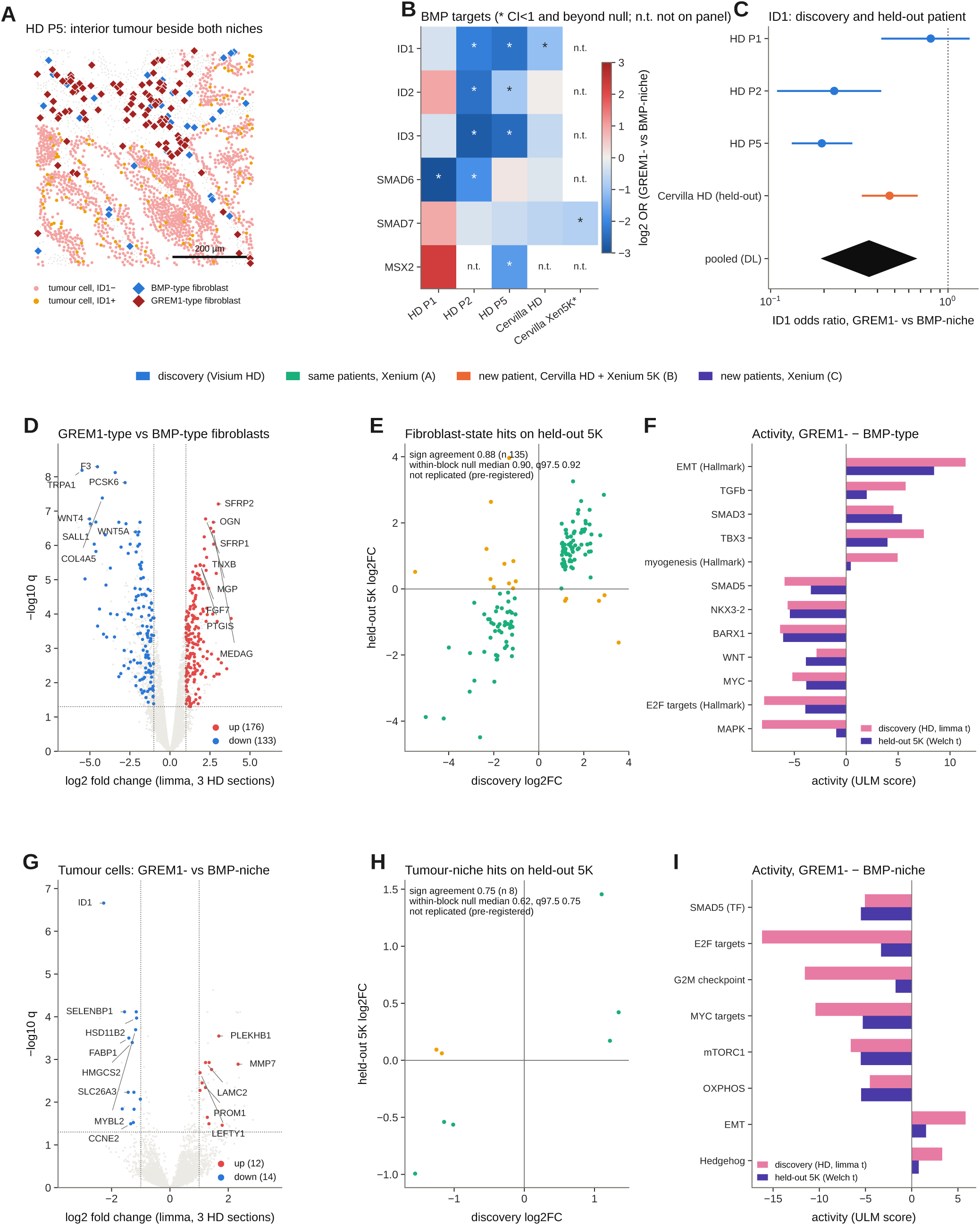
Fibroblast-state programmes and the tumour response. **A**, Visium HD P5 window of interior tumour cells (*d < −*50 µm) next to both fibroblast niches; ID1^+^ tumour nuclei in yellow; scale bar 200 µm. **B**, Mantel–Haenszel odds ratio (GREM1-niche versus BMP-niche tumour cells, stratified by distance and fibroblast count) for BMP target genes; * CI *<* 1 and beyond the spatial-null 2.5% quantile; n.t., not on panel. **C**, ID1 odds ratio per section; the held-out Cervilla patient (pre-registered confirmation) gives 0.47 [0.33–0.67]. The diamond pools discovery and held-out sections and is descriptive only; the confirmation rests on the Cervilla test. ID1 is not on the Xenium 5K panel; there SMAD7 is lower (0.60 [0.49–0.77]). **D**–**I**, Whole-transcriptome comparisons. **D**, GREM1-type versus BMP-type fibroblasts (limma-voom on per-section pseudobulks, 3 HD sections; defining genes and non-fibroblast-owned genes excluded); coloured, 309 discovery hits (176 up, 133 down). **E**, Discovery versus held-out Xenium 5K log_2_FC for the 135 testable hits; sign agreement 0.88 against a within-block null median 0.90 and 97.5% quantile 0.92: not replicated (pre-registered). **F**, Pathway and TF activity (decoupler ULM; PROGENy, Hallmark, CollecTRI), GREM1-type minus BMP-type, discovery (limma *t*) and held-out 5K (Welch *t*). **G**, Tumour cells in GREM1 versus BMP niches (tumour-owned genes); 26 hits (12 up, 14 down), ID1 ranked first. **H**, Held-out 5K for the 8 testable hits; agreement 0.75, null median 0.62 and 97.5% quantile 0.75: not replicated (tie). **I**, Activity, GREM1-niche minus BMP-niche; SMAD5 discovery *−*5.1, 5K *−*5.5. Pathway activities were tested by a same-sign rule without a spatial null.

The pre-registered held-out test did not replicate this list. On Xenium 5K, 135 hits were testable, and their sign agreement was 0.881; but permuting state labels among fibroblasts within 400 µm blocks gave a median of 0.896 and a 97.5% quantile of 0.919 (panel b). BMP-type and GREM1-type fibroblasts occupy different 400 µm blocks, so a within-block permutation keeps most of the contrast: at this scale, the state transcriptome cannot be separated from position. Agreement with the same patients’ Xenium sections was 0.94, 0.71 and 0.94 (17 panel genes each). Because no gene passed the held-out test, the pre-registered locked marker panel built from replicated genes is empty, and its survival test was not run.

Pathway and transcription-factor activity, which has no spatial null and is read by a same-sign rule, separated the states clearly (panel c; Supplementary Table 5). GREM1-type fibroblasts had higher TGF-*β* (PROGENy +5.7), SMAD3 (CollecTRI +4.6; Xenium 5K +5.4) and EMT activity (Hallmark +11.5; 5K +8.5), and lower E2F, G2M, MYC and mTORC1 activity: a TGF-*β*/SMAD3-driven, matrix-producing, non-proliferating CAF state. BMP-type fibroblasts had higher BMP-responsive SMAD5 (*−*5.9 in the GREM1-minus-BMP contrast; 5K *−*3.4), NKX3-2 and BARX1 activity, higher WNT and MAPK activity, and expressed WNT4, WNT5A and F3: a BMP-active, telocyte-like state. Replication by sign was 8/8 for PROGENy, 8/9 for Hallmark and 24/28 for CollecTRI. Hallmark myogenesis was also higher in GREM1-type fibroblasts, which keeps the smooth-muscle question open for the bulk score.

#### Tumour cells in the GREM1 niche

If GREM1-type fibroblasts antagonise BMP locally, adjacent tumour cells should show less BMP pathway output. We compared interior tumour cells (*d < −*50 µm) whose fibroblast neighbours within 30 µm included a GREM1-type and no BMP-type cell (GREM1 niche) with the reverse (BMP niche), by Mantel–Haenszel odds ratios stratified by distance and fibroblast count, with a 400 µm block bootstrap and a block-permuted state null (Fig. 6A–C).

The pre-registered primary readout, stemness (LGR5, ASCL2, AXIN2 or RNF43), failed: odds ratios 0.99, 1.28 and 1.19 in discovery, and 0.72 (0.60–0.89) on Xenium 5K, in the *opposite* direction. The hypothesis that GREM1 releases tumour stemness is refuted in these data. The secondary readout, ID1, was positive: it was detected in 23% versus 38%, 12% versus 43% and 17% versus 63% of GREM1-niche versus BMP-niche tumour cells (odds ratios 0.80, 0.23 and 0.19; P2 and P5 below the null), and in P2 and P5 ID2 and ID3 fell too. ID2 is informative because fibroblast spill would push it up. A confirmation rule recorded before the held-out run was met in the only external section carrying ID1, BMP4 and GREM1: Cervilla Visium HD, 0.47 (0.33–0.67), below the null 2.5% quantile of 0.65. ID1 is on no Xenium panel; on Xenium 5K, the BMP target SMAD7 was lower in GREM1-niche tumour cells (0.60, 0.49–0.77), which is consistent but not the registered test. Under our evidence hierarchy the finding is VALIDATED by levels B and D (the held-out patient plus a pre-registered bulk coupling, below); the automated meta-analysis rule could not be applied, because only one external section measures ID1.

A genome-wide comparison of the same tumour cells, with tumour-owned genes only, ranked ID1 first among 11,403 genes (log_2_FC *−*2.27, *q* = 2 *×* 10^−7^; Fig. 6G). Of 26 hits, 14 were lower in the GREM1 niche (ID1, SELENBP1, HSD11B2, MYBL2, FABP1, HMGCS2, SLC26A3, CCNE2) and 12 higher (MMP7, LAMC2, PROM1, LEFTY1, PLEKHB1). The held-out gene-level test failed by a tie (8 testable genes; agreement 0.75 against a null 97.5% quantile of 0.75; panel e), while same-patient Xenium agreed in 6/7, 7/7 and 7/7. At pathway level, SMAD5 activity was lower in GREM1-niche tumour cells in discovery (*−*5.1) and on held-out Xenium 5K (*−*5.5), as were E2F, G2M, MYC, mTORC1 and oxidative-phosphorylation programmes, while EMT and Hedgehog programmes were higher (Hallmark 16/19 and CollecTRI 11/14 replicated by sign; panel f). PROGENy pathway scores did not replicate (1/6). Tumour cells beside GREM1-type fibroblasts therefore show a pathway-level loss of BMP response, centred on ID1 and SMAD5, together with a less proliferative, more invasive-looking state.

Single-cell ownership supports a paracrine route: in the Flex reference, GREM1 is made by CAFs (18.7 counts per 10,000, CP10k) and a smooth-muscle-like state (20.8), BMP5 by fibroblasts and myofibroblasts, and tumour cells express BMPR2, BMPR1A, ACVR1 and ID1 (Supplementary Fig. 14). At population level, tumours with a higher fibroblast BMP balance had higher residualised tumour ID1–3 expression (TCGA *ρ* = +0.31, above a random-set 95% quantile of 0.15; GEO pooled +0.22 [0.12–0.32]).

### Measured against the mouse model, the stromal BMP balance is one-sided in human tumours

The prior model of this axis is a balance between GREM1 and the BMP potentiator ISLR, built chiefly in a mouse colonic fibroblast line, mouse organoids and a mouse metastasis model ^13^. ISLR and FOXL1 are measured on Visium HD but are on none of the Xenium panels we use, including the 5K, so we tested that model on the four Visium HD sections (three discovery, one held-out), with the same 400 µm block bootstrap and block-permutation nulls used throughout, and added a third fibroblast state to the two already defined: ISLR-type (ISLR detected, GREM1 not), corresponding to the ISLR^+^ CAF that the prior work described as distinct from GREM1^+^ CAFs (Supplementary Figs 15–18).

#### What holds

ISLR is fibroblast-owned in the Flex reference, as Kobayashi et al. reported (CAF 7.8, proliferating fibroblast 9.4 CP10k, against GREM1 18.7 in CAFs). Their distance ordering holds and sharpens: GREM1-type fibroblasts stood further from tumour cells than either other state in 8/8 sections (median 66–341 µm against 15–62 µm for BMP-type; within 250 µm of the border, 49–135 against 14–49 µm, Supplementary Fig. 15D), and only 5–28% of GREM1-type fibroblasts lay within 30 µm of a tumour cell, against 36–76% of BMP-type and 31–51% of ISLR-type. Measured cell by cell against a border that no expression value defined, the ordering reproduces the in situ hybridisation result they reported in desmoplastic rectal cancer.

#### What does not

First, ISLR marks no state opposite GREM1. GREM1 and ISLR were co-detected in fibroblasts as often as chance or slightly more (odds ratio 1.00, 1.10, 1.30, 1.38; 4/4 sections), not less; ISLR detection did not separate GREM1-type from BMP-type fibroblasts (log_2_ rate ratio *−*0.02 to *−*0.58, beyond the block-permutation null in 1/4 sections; fibroblast pseudobulk log_2_FC +0.16, *q* = 0.59); and ISLR-type fibroblasts were enriched in the tumour bed (6.9–16.6%) rather than at the front (3.7–7.8% outside), on the same side as the BMP-producing state (Supplementary Fig. 15B,C). Second, ISLR does not act as a potentiator where it sits: tumour cells whose only fibroblast neighbours were ISLR-type had *lower* ID1 than those beside BMP-type fibroblasts (odds ratios 0.65, 0.50, 0.32, 0.61; 4/4 beyond the null; ID1–3 4/4), and were indistinguishable from GREM1-niche tumour cells (ID1 0.42–0.91, 1/4 beyond the null). What predicts tumour BMP-target output next to a fibroblast is whether that fibroblast makes a BMP ligand, not whether it carries ISLR. Third, the FOXL1 arm runs the other way. FOXL1 was lower in GREM1-type than in BMP-type fibroblasts in 4/4 sections (log_2_ *−*0.95 to *−*4.46; 3/4 beyond the null), as was PDGFRA in 8/8 (all beyond the null), so the FOXL1^+^PDGFRA^+^ telocyte identity that the mouse work assigns to Grem1^+^ fibroblasts belongs here to the BMP4/5-producing bed state (Supplementary Fig. 16). TGFB1 itself was also lower in the GREM1-type state (4/5 beyond the null) although TGF-*β*/SMAD3 pathway activity is higher in it (above): the ligand and the response sit on opposite sides of the border. Fourth, ACTA2 and TAGLN did not separate the states consistently (log_2_ *−*0.55 to +0.47, signs mixed across sections; Supplementary Fig. 15E), so at single-gene level we neither confirm nor refute the myofibroblastic identity of GREM1^+^ CAFs, while Hallmark myogenesis was higher in them (above). Fifth, their downstream readouts do not follow: the stem set was flat (odds ratios 0.72–1.28 over eight sections, no consistent direction, as reported above) and KRT20 moved in both directions (0.33–1.67; 2/5 beyond the null), so the Lgr5-release and Krt20-block mechanism of the tumoroid experiments is not visible in these tumours (Supplementary Fig. 17).

#### In bulk

Across TCGA and the three GEO cohorts GREM1 and ISLR rose together rather than in opposition (Spearman +0.64 to +0.74). Tumours that were GREM1-high and ISLR-low were correspondingly rare (13/621, 11/566, 2/177 and 0/55 against 69, 63, 20 and 6 expected under independence), which reproduces their observation that the two adverse groups barely overlap—but because both genes track stromal content, not because they are opposed; both were highest in the stroma-rich CIT subtype C4 (83% GREM1-high, 0% ISLR-low). The GREM1 arm replicated: for GREM1 measured as in the prior work (single gene, not residualised), the pooled overall-survival hazard ratio was 1.45 [1.06–1.97] for the top tertile and 1.22 [1.00–1.47] per s.d. The ISLR arm did not. ISLR-high was adverse rather than favourable (1.36 [1.05–1.76]), ISLR-low was null (1.00 [0.80–1.25]), and their ISLR *−* GREM1 balance carried no information (0.99 [0.88–1.10]), whereas our state-based fibroblast BMP balance gave 0.84 [0.72–0.98] with its discovery cohort TCGA included (GEO only 0.88 [0.74–1.05], the failed pre-registered primary). Adjusting for CIT subtype in GSE39582 removed the GREM1 association (1.00 [0.82–1.23]) and left ISLR-low adverse (1.61 [1.08–2.41]), the direction they reported: in bulk, both genes are readouts of CMS4-like stromal biology, and the ISLR direction only appears once stromal content is accounted for. Finally, ISLR did not behave as a potentiator at population level either, being negative against residualised tumour ID1–3 in all four cohorts (*ρ −*0.06 to *−*0.10), where the fibroblast BMP balance was positive in the three larger ones (+0.24 to +0.31; 0.00 in GSE17537, *n* = 55; Supplementary Fig. 18).

### Signalling between immune cells and fibroblasts acts at niche scale

#### Contact-scale co-expression is negative

For 4,248 reference-directional LIANA pairs, we compared ligand–receptor co-expression in touching cell pairs (*≤*15 µm) with near but non-touching pairs (30–60 µm), against 500 random gene pairs. None of 1,233 testable Visium HD pairs was contact-specific; on Xenium, 1 of 158 pairs met the two-of-three rule (CXCL11–CXCR3), and 4.6% of 415 slide-tests exceeded the null 95th percentile, against 5% expected. A planted 9-point juxtacrine induction was detected in 68% of slides, so this negative is powered (Fig. 7D; Supplementary Fig. 11, row N28).

**Figure 7:**
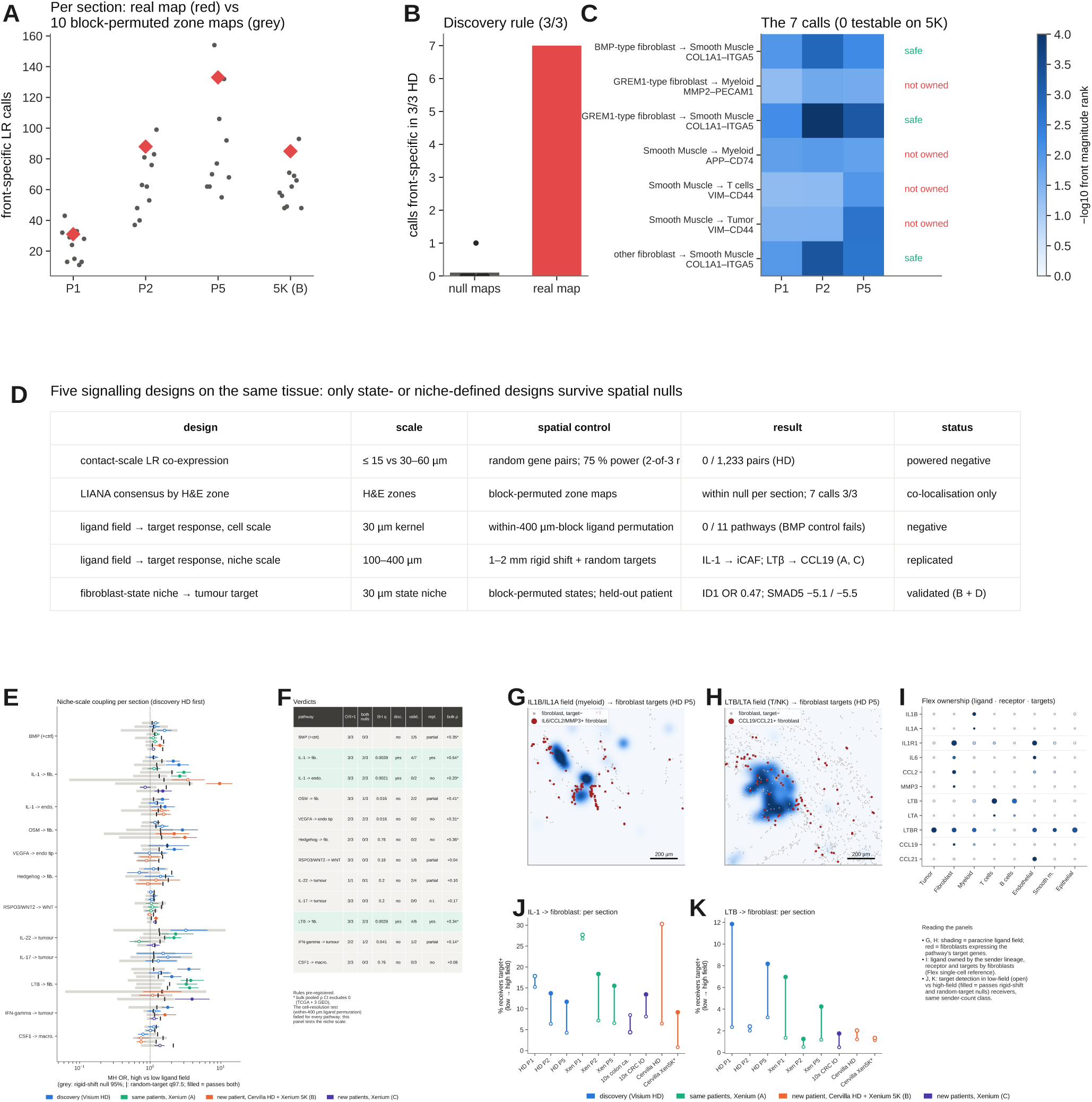
Niche-scale signalling. **A**–**D**, Zone-resolved cell–cell communication with LIANA consensus scoring: front-specific interactions (magnitude rank *≤*0.05 at the front in 3/3 sections and not in the bed), with fibroblasts split into BMP-type, GREM1-type and other; marked, beyond a 10-map block null and contamination-safe by ownership; held-out Xenium 5K status. **A**, Front-specific calls per section on the real zone map (red) and on ten block-permuted zone maps (grey). **B**, Calls front-specific in 3/3 discovery sections (7) versus the null maps (mean 0.1). **C**, The seven calls, *−* log_10_ front magnitude rank per section, with the ownership flag; none is testable on Xenium 5K. **D**, The five signalling designs applied to this tissue and their outcome. Co-expression, not receptor engagement. **E**, **F**, Ligand-field*→*target-response at niche scale. **E**, Odds ratio per pathway and section against a null in which the ligand field is read at receiver positions shifted 1–2 mm (grey) and against 20 detection-matched random target sets (*|*); filled, passes both. **F**, Verdicts (test registered after the cell-resolution result): IL-1*→*fibroblast (*q* 0.003; 4/7 validation sections at A, B, C with the IL6/CCL2/MMP3 readout; 2/6 with CCL2 + MMP3 only; bulk *ρ* +0.54) and LT*β →*fibroblast (*q* 0.003; 4/6 at A, C, also with CCL19 only; not in the held-out patient; bulk *ρ* +0.34); IL-1*→*endothelium coupled but not replicated; the BMP control is not coupled by ligand amount. **G**–**K**, The two axes in situ. **G**, **H**, HD P5 windows; shading, ligand field (myeloid IL1B/IL1A; T/NK LTB/LTA); red, fibroblasts detecting the targets (IL6/CCL2/MMP3; CCL19/CCL21); scale bars 200 µm. **I**, Flex ownership by lineage; CCL21 is mainly endothelial, so CCL19 is the cleaner fibroblast readout. **J**, **K**, Target-positive receivers in low-field (open) versus high-field receivers per section; filled, passes both nulls; * Cervilla Xenium 5K, where IL6 was the only IL-1 target on the panel.

#### Zone-resolved communication with a standard tool

We then ran LIANA consensus scoring ^20^ (CellPhoneDB, Connectome, log2FC, NATMI, SingleCellSignalR and CellChat-like methods) separately in the tumour bed, the front and distant stroma, with fibroblasts split into BMP-type, GREM1-type and other, a 10-map block null, an ownership filter and held-out replication on Xenium 5K (Fig. 7A–D; Supplementary Fig. 19). In each section, the number of front-specific interactions on the real zone map (31, 88 and 133 in P1, P2 and P5; 85 on Xenium 5K) lay inside the range obtained on ten block-permuted zone maps (11–43, 37–99, 55–154 and 48–93): single-section zone calls could not be distinguished from arbitrary zones. Requiring the call in all three tumours left seven interactions, against a null mean of 0.1 (95th percentile 0.55). Three of them were contamination-safe by ownership, and all three were the same generic collagen–integrin pair (COL1A1*→*ITGA5 from fibroblasts to smooth muscle); the other four (for example GREM1-type fibroblast MMP2*→*PECAM1, VIM*→*CD44) used genes that are not owned by the sender or the receiver. None could be tested on Xenium 5K, whose panel lacks COL1A1, MMP2, CD74 and VIM. The most highly ranked interactions were the same collagen–CD44 pairs in the bed, at the front and in distant stroma (Supplementary Fig. 19). A standard CCC tool, run with a spatial null and an ownership filter, therefore recovered co-localisation, not front-specific signalling (Fig. 7D compares the five signalling designs applied to the same tissue).

#### Receiver-response tests

We therefore read signalling through receiver-owned target genes. In a ligand-field*→* target-response (LFTR) test, each receiver was assigned a Gaussian-weighted ligand field from senders within 100 µm, and receivers in the high and low tertiles were compared within classes of equal sender count, so that the contrast is ligand amount, not co-location. The screen was calibrated (2.0% false exceedances against 2.5% expected). At cell resolution no pathway passed, including the BMP positive control (Supplementary Fig. 20): permuting ligand among senders within 400 µm blocks reproduced every response. A follow-up, registered after that failure and before it was run, replaced the null with a rigid 1–2 mm shift of the ligand field. Two immune-to-fibroblast axes were then coupled in discovery and replicated (Fig. 7E–K):

- myeloid IL-1 (IL1B/IL1A)*→*fibroblast IL6, CCL2 and MMP3, an inflammatory-CAF (iCAF) programme: discovery odds ratios 1.13, 2.12 and 2.55 (*q* = 0.003); 4/7 validation sections at levels A, B and C; bulk *ρ* +0.54 [0.50, 0.58]. With fibroblast-specific targets (CCL2 and MMP3; IL6 is also endothelial) discovery held in 3/3 but replication was partial (2/6);
- lymphoid LT*β* (LTB/LTA)*→*fibroblast CCL19 and CCL21, a lymphoid-organiser programme: 3.53, 1.36 and 2.02 (*q* = 0.003); 4/6 validation sections at levels A and C, robust to a CCL19-only readout; bulk *ρ* +0.34 [0.29, 0.38]. It failed in the held-out patient (Visium HD and Xenium 5K).

In tissue, IL6/CCL2/MMP3-positive fibroblasts clustered within dense myeloid IL-1 fields, and CCL19-positive fibroblasts within T/NK LTB fields; Flex ownership supported both directions (IL1B myeloid 4.67 versus 0.13 CP10k in fibroblasts; IL1R1 fibrob-last 3.16; LTB T and B cells; CCL19 fibroblast-high). The BMP control was not coupled by ligand amount either; BMP signalling was detected only by the fibroblast-*state* design, consistent with availability set by an antagonist. Neither target score predicted survival (IL-1 1.02 [0.84–1.25]; LT*β* 1.01 [0.88–1.15]). The same data say “no signalling” or “replicated signalling” depending on the scale the null preserves, and that scale, about 100–400 µm, matches the stromal scale from MIW.

### Clinical translation: a GREM1-state score in four cohorts

To carry the spatial axis to outcome, we scored bulk tumours with a BMP-type score (BMP4, BMP5, WNT4, F3, CXCL14) and a GREM1-type score (GREM1, SFRP2), each residualised on fibroblast abundance so that it measures fibroblast *state* rather than stromal amount (Fig. 8A–C). In TCGA (578 patients, 115 deaths), the balance was associated with longer overall survival (hazard ratio 0.75 per s.d., 0.62–0.92) and the GREM1-type score with shorter survival (1.36, 1.11–1.66; Supplementary Table 6). The pre-registered GEO replication failed for its primary endpoint, the balance: 0.88 (0.74–1.05), probably a winner’s curse in the discovery cohort, TCGA. The pre-stated secondary, the GREM1-type score, replicated without heterogeneity: GEO-only 1.18 (1.05–1.34; *I*^2^ = 0), and 1.23 (1.11–1.36) over all four cohorts (1,319 patients, 380 deaths); leaving out any cohort kept the pooled interval above 1 (Supplementary Table 7). The BMP-type score alone carried no signal (1.00), and no score was associated with relapse in the pooled adjusted analysis (the BMP-type score was univariably adverse for relapse in GSE39582 only).

**Figure 8:**
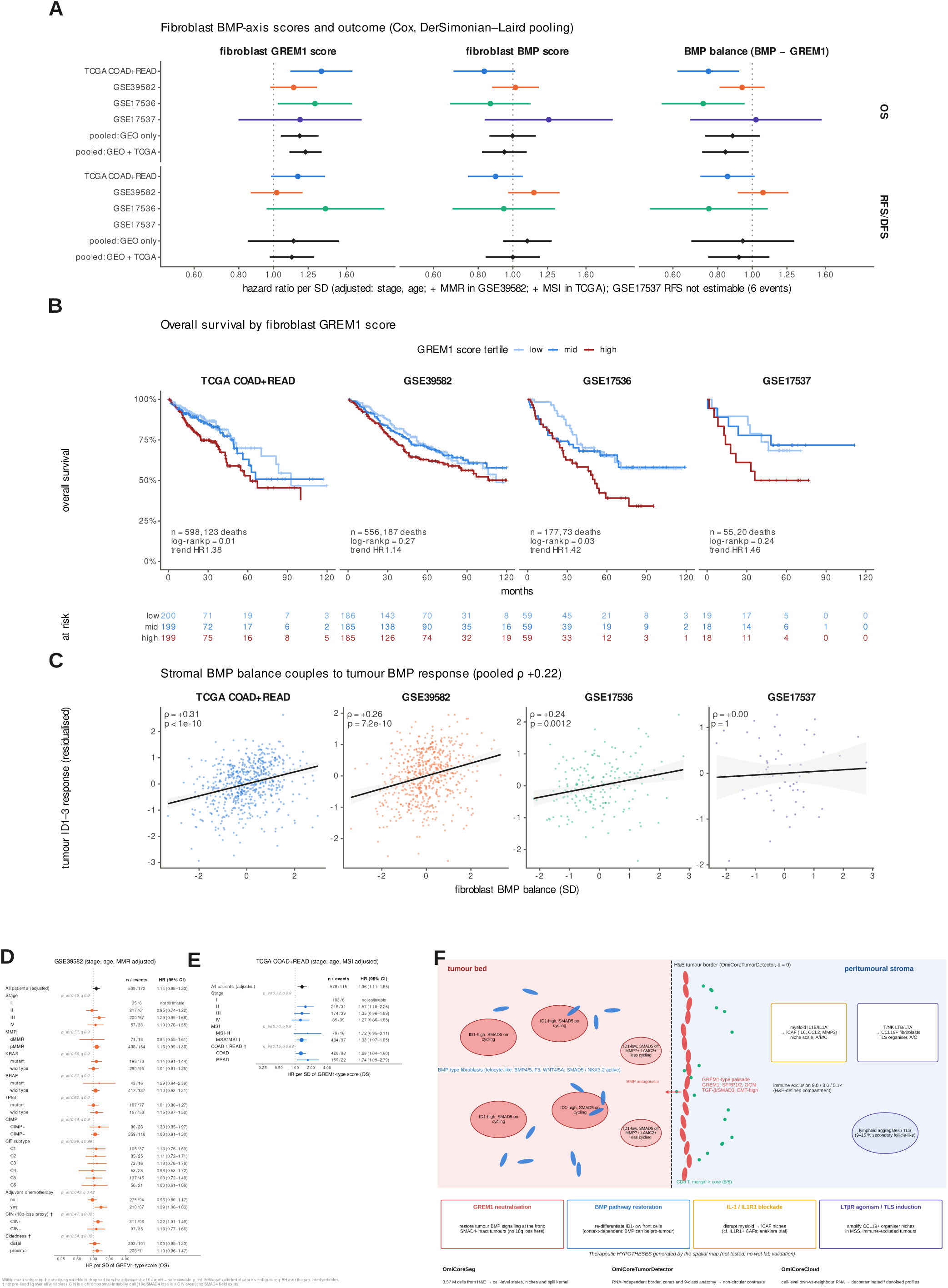
Clinical translation and model. **A**, Cox hazard ratio per s.d. of the fibroblast GREM1-type, BMP-type and balance scores (residualised on fibroblast abundance; adjusted for stage and age, + MMR in GSE39582, + MSI and site in TCGA), per cohort and DerSimonian–Laird pooled, for overall survival and relapse/disease-free survival. GREM1-type: GEO-only 1.18 [1.05–1.34], with TCGA 1.23 [1.11–1.36], *I*^2^ 0. The pre-registered primary (balance, OS, GEO only) failed: 0.88 [0.74–1.05]. **B**, OS by GREM1-score tertile with log-rank *P*, trend hazard ratio and numbers at risk. **C**, Fibroblast BMP balance versus residualised tumour ID1–3; TCGA *ρ* +0.31; GEO pooled +0.22 [0.12–0.32]. **D**, **E**, GREM1-type score within clinical and molecular subgroups of GSE39582 (**D**) and TCGA (**E**); diamond, whole cohort; *p*_int_, likelihood-ratio interaction; *q*, Benjamini–Hochberg over pre-listed variables; *†*, not pre-listed. No interaction reaches *q <* 0.05. **F**, Working model relative to the H&E border: BMP4/5-type, telocyte-like fibroblasts (SMAD5/NKX3-2/BARX1 activity) thread between ID1-high glands in the bed; at the front, a palisade of GREM1/SFRP2-type fibroblasts (TGF-*β*/SMAD3, EMT-high, non-proliferating) accompanies tumour cells with low ID1 and SMAD5 activity; CD8 T cells and lymphoid aggregates collect at the margin; myeloid IL-1*→*iCAF and lymphoid LT*β →*CCL19 couplings act at niche scale. Call-outs mark therapeutic hypotheses (GREM1 neutralisation in SMAD4-intact tumours, BMP restoration, IL-1/IL1R1 blockade, LT*β*R/TLS induction); none was tested here. The model is associative; niche positions of the immune axes are schematic.

Three findings limit this score. First, it is a replication: GSE39582 very probably overlaps the cohort in which Kobayashi et al. ^13^ reported poorer survival for GREM1-high tumours. Second, in GSE39582, adding CIT subtype, KRAS, BRAF, TP53, CIMP and adjuvant chemotherapy removed the association (0.97, 0.78–1.21; *n* = 280), so it is not independent of molecular subtype. Third, its gain in discrimination was small (pooled Δ*C* +0.009, *−*0.001 to 0.019). It was consistent across stage, MMR/MSI, mutation, CIMP and chemotherapy subgroups (no interaction *q <* 0.05; MSS/MSI-L 1.33, 1.07–1.65; Fig. 8D,E). An exploratory interaction with adjuvant chemotherapy in stage II–III GSE39582 (1.79, 1.20–2.67) is confounded by indication. A SMAD4 interaction could not be tested.

The marker panel we had pre-registered to improve on this score could not be built, because the gene-level state transcriptome did not replicate (above). The two-gene GREM1-type score therefore remains the bulk form of the axis. Fig. 8F assembles the validated results relative to the H&E border, and Table 1 lists every claim with its evidence level and verdict.

**Table 1:**
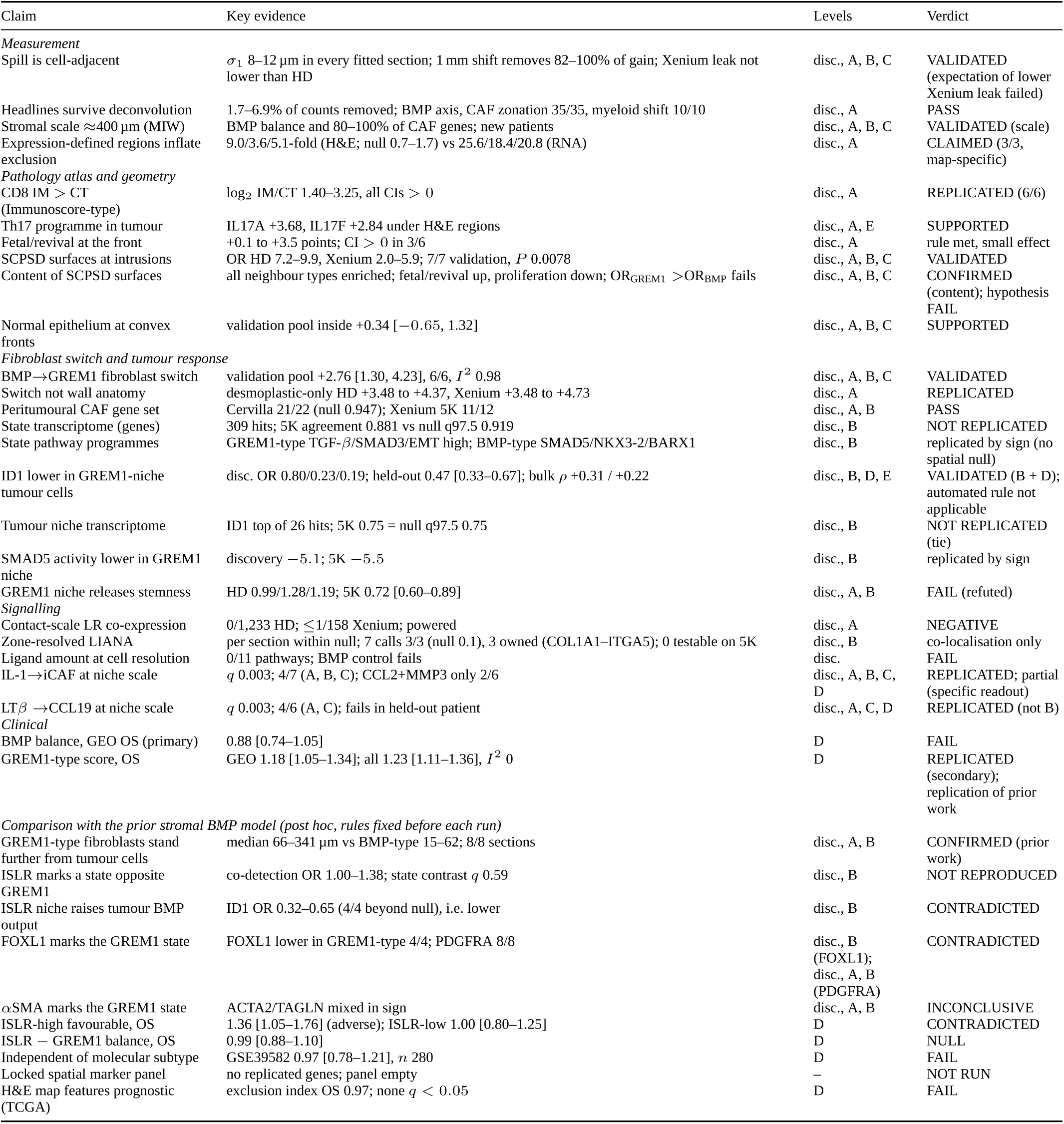
Claims, evidence level and verdict. Evidence levels: disc., discovery Visium HD (P1, P2, P5); A, same patients on Xenium; B, held-out patient (Cervilla Visium HD; Xenium 5K is its platform replicate); C, new patients on 10x Xenium; D, bulk TCGA and GEO; E, Flex single-cell plausibility. Verdicts follow rules fixed before the runs (Methods). OS, overall survival; HR per s.d.

| Claim | Key evidence | Levels | Verdict |
|---|---|---|---|
| *Measurement* |  |  |  |
| Spill is cell-adjacent | σ<sub>1</sub> 8–12 µm in every fitted section; 1 mm shift removes 82–100% of gain; Xenium leak not lower than HD | disc., A, B, C | VALIDATED (expectation of lower Xenium leak failed) |
| Headlines survive deconvolution | 1.7–6.9% of counts removed; BMP axis, CAF zonation 35/35, myeloid shift 10/10 | disc., A | PASS |
| Stromal scale ≈400 µm (MIW) | BMP balance and 80–100% of CAF genes; new patients | disc., A, B, C | VALIDATED (scale) |
| Expression-defined regions inflate exclusion | 9.0/3.6/5.1-fold (H&E; null 0.7–1.7) vs 25.6/18.4/20.8 (RNA) | disc., A | CLAIMED (3/3, map-specific) |
| *Pathology atlas and geometry* |  |  |  |
| CD8 IM > CT (Immunoscore-type) | log<sub>2</sub> IM/CT 1.40–3.25, all CIs > 0 | disc., A | REPLICATED (6/6) |
| Th17 programme in tumour | IL17A +3.68, IL17F +2.84 under H&E regions | disc., A, E | SUPPORTED |
| Fetal/revival at the front | +0.1 to +3.5 points; CI > 0 in 3/6 | disc., A | rule met, small effect |
| SCPSD surfaces at intrusions | OR HD 7.2–9.9, Xenium 2.0–5.9; 7/7 validation, *P* 0.0078 | disc., A, B, C | VALIDATED |
| Content of SCPSD surfaces | all neighbour types enriched; fetal/revival up, proliferation down; OR<sub>GREM1</sub> >OR<sub>BMP</sub> fails | disc., A, B, C | CONFIRMED (content); hypothesis FAIL |
| Normal epithelium at convex fronts | validation pool inside +0.34 [−0.65, 1.32] | disc., A, B, C | SUPPORTED |
| *Fibroblast switch and tumour response* |  |  |  |
| BMP→GREM1 fibroblast switch | validation pool +2.76 [1.30, 4.23], 6/6, *I*<sup>2</sup> 0.98 | disc., A, B, C | VALIDATED |
| Switch not wall anatomy | desmoplastic-only HD +3.48 to +4.37, Xenium +3.48 to +4.73 | disc., A | REPLICATED |
| Peritumoural CAF gene set | Cervilla 21/22 (null 0.947); Xenium 5K 11/12 | disc., A, B | PASS |
| State transcriptome (genes) | 309 hits; 5K agreement 0.881 vs null q97.5 0.919 | disc., B | NOT REPLICATED |
| State pathway programmes | GREM1-type TGF-β/SMAD3/EMT high; BMP-type SMAD5/NKX3-2/BARX1 | disc., B | replicated by sign (no spatial null) |
| ID1 lower in GREM1-niche tumour cells | disc. OR 0.80/0.23/0.19; held-out 0.47 [0.33–0.67]; bulk ρ +0.31 / +0.22 | disc., B, D, E | VALIDATED (B + D); automated rule not applicable |
| Tumour niche transcriptome | ID1 top of 26 hits; 5K 0.75 = null q97.5 0.75 | disc., B | NOT REPLICATED (tie) |
| SMAD5 activity lower in GREM1 niche | discovery −5.1; 5K −5.5 | disc., B | replicated by sign |
| GREM1 niche releases stemness | HD 0.99/1.28/1.19; 5K 0.72 [0.60–0.89] | disc., A, B | FAIL (refuted) |
| *Signalling* |  |  |  |
| Contact-scale LR co-expression | 0/1,233 HD; ≤1/158 Xenium; powered | disc., A | NEGATIVE |
| Zone-resolved LIANA | per section within null; 7 calls 3/3 (null 0.1), 3 owned (COL1A1–ITGA5); 0 testable on 5K | disc., B | co-localisation only |
| Ligand amount at cell resolution | 0/11 pathways; BMP control fails | disc. | FAIL |
| IL-1→iCAF at niche scale | *q* 0.003; 4/7 (A, B, C); CCL2+MMP3 only 2/6 | disc., A, B, C, D | REPLICATED; partial (specific readout) |
| LTβ →CCL19 at niche scale | *q* 0.003; 4/6 (A, C); fails in held-out patient | disc., A, C, D | REPLICATED (not B) |
| *Clinical* |  |  |  |
| BMP balance, GEO OS (primary) | 0.88 [0.74–1.05] | D | FAIL |
| GREM1-type score, OS | GEO 1.18 [1.05–1.34]; all 1.23 [1.11–1.36], *I*<sup>2</sup> 0 | D | REPLICATED (secondary); replication of prior work |
| *Comparison with the prior stromal BMP model (post hoc, rules fixed before each run)* |  |  |  |
| GREM1-type fibroblasts stand further from tumour cells | median 66–341 µm vs BMP-type 15–62; 8/8 sections | disc., A, B | CONFIRMED (prior work) |
| ISLR marks a state opposite GREM1 | co-detection OR 1.00–1.38; state contrast *q* 0.59 | disc., B | NOT REPRODUCED |
| ISLR niche raises tumour BMP output | ID1 OR 0.32–0.65 (4/4 beyond null), i.e. lower | disc., B | CONTRADICTED |
| FOXL1 marks the GREM1 state | FOXL1 lower in GREM1-type 4/4; PDGFRA 8/8 | disc., B (FOXL1); disc., A, B (PDGFRA) | CONTRADICTED |
| αSMA marks the GREM1 state | ACTA2/TAGLN mixed in sign | disc., A, B | INCONCLUSIVE |
| ISLR-high favourable, OS | 1.36 [1.05–1.76] (adverse); ISLR-low 1.00 [0.80–1.25] | D | CONTRADICTED |
| ISLR − GREM1 balance, OS | 0.99 [0.88–1.10] | D | NULL |
| Independent of molecular subtype | GSE39582 0.97 [0.78–1.21], *n* 280 | D | FAIL |
| Locked spatial marker panel | no replicated genes; panel empty | – | NOT RUN |
| H&E map features prognostic (TCGA) | exclusion index OS 0.97; none *q* < 0.05 | D | FAIL |

## Discussion

Three measurements that standard pipelines approximate—cells, an expression-blind tumour compartment and the separation of a cell’s own RNA from its neighbours’—changed what could be said about the CRC front. The H&E region removed a three- to fivefold inflation of immune exclusion, reinstated a Th17 programme and withdrew an artefactual CCR7 loss. The spill kernel showed that contamination is a one-cell phenomenon on both segmented platforms, and the multiscale contrast that stromal zonation acts at about 400 µm; this scale separation, a property of the tissue, is consistent with stromal results surviving every correction. The block null removed two-thirds of zone programmes that a gene-permutation test had declared significant. The external replications of the BMP switch, SCPSD and CAF zonation used an RNA mask, so those three findings do not require the detector.

With these measurements, the front resolves into a small number of organised programmes. Fibroblasts occupy two segregated states. In the tumour bed, a minority of fibroblasts keep a BMP-producing, telocyte-like programme (BMP4/5, WNT4, WNT5A, F3, CXCL14; SMAD5, NKX3-2 and BARX1 activity) that resembles the crypt-top lamina propria ^11^. At the front, GREM1/SFRP2-type fibroblasts form a palisade set back from tumour cells, with TGF-*β*/SMAD3 and EMT activity and little proliferation, two to five times more frequent than in normal stroma or muscle. The switch is anchored to an expression-independent border, replicates on a second platform, in a new patient and in a further public section, survives deconvolution and persists after removal of muscle and mucosa. Beside the GREM1 state, tumour cells lose BMP-target output, centred on ID1 and SMAD5 activity, and are less proliferative and more EMT-like, at the same front where fetal/revival programmes are modestly enriched. The arrangement resembles a tumour that keeps a crypt-top-like BMP niche inside and meets a crypt-base-like antagonist niche at its edge. The immune side of the same front is margin-restricted: CD8, CD4, B and plasma cells and a Crohn-like ring of lymphoid aggregates collect outside the border, NK cells are excluded from the bed, and SPP1^+^ macrophages enrich inside it. Signals from immune cells reach fibroblasts at niche scale (IL-1 to iCAF programmes; LT*β* to CCL19-producing fibroblasts), not at cell contact, where co-expression was no more specific than random gene pairs.

What is new, given that bulk GREM1 is already linked to outcome ^13^, is the spatial and cellular resolution: discrete BMP-type and GREM1-type fibroblast states inverting across the H&E border on independent patients and platforms; their pathway programmes; a niche-to-tumour BMP-response deficit confirmed on a held-out patient and coupled at population level; and a bulk score that measures fibroblast state rather than stromal amount. The pathway result may explain why stromal GREM1 and CMS4-like biology are hard to separate: the GREM1 state is itself a TGF-*β*/EMT-high CAF state, and in GSE39582 the score lost its association after adjustment for molecular subtype.

### Relation to the mouse stromal BMP–GREM1 model

The stromal BMP axis in colorectal cancer was defined by work done mostly in mice: a mouse colonic fibroblast line for the GREM1 and ISLR functions and for the TGF-*β*–FOXL1 regulation, mouse tumoroids for the stemness and differentiation mechanism, and a mouse liver-metastasis model for the therapeutics ^13^. Its human evidence is bulk, sorted-cell and single-cell expression profiles and in situ hybridisation, which cannot say which fibroblast state stands where, or what the tumour cells beside it do. Table 2 sets that model against the measurements made here. Five of its statements hold in human tissue: GREM1 and ISLR are fibroblast-owned; GREM1^+^ fibroblasts stand back from tumour cells while the other states sit close; tumour cells beside them lose BMP-target output; bulk GREM1 is adverse; and both genes are highest in stroma-rich tumours. Five do not, and one (*α*SMA identity) is inconclusive. Four of the five are set out here; the fifth, the stemness readout, is discussed in the next paragraph. ISLR is not the human counter-state—it is co-expressed with GREM1 at chance, sits on the tumour-bed side, and does not separate the two states—so the balance that names the prior model is not the axis that inverts across the human border; that axis is between fibroblasts that make BMP ligand and fibroblasts that make its antagonist. ISLR does not potentiate: tumour cells beside ISLR-type fibroblasts lose BMP-target output just as they do beside GREM1-type fibroblasts, and in bulk ISLR does not track tumour ID1–3. The FOXL1^+^PDGFRA^+^ telocyte identity belongs here to the BMP-producing state, not to the GREM1 state, which matches the BMP-producing telocytes of the normal mouse colon ^11^ but is the opposite of the assignment in the mouse tumour model; TGFB1 transcript is likewise on the bed side although TGF-*β*/SMAD3 activity is highest at the front. And the prognostic model is one-sided: GREM1-high is adverse, ISLR-high is adverse too rather than favourable, and the ISLR *−* GREM1 balance carries nothing, although adjusting for molecular subtype recovers their ISLR-low direction and removes the GREM1 one.

**Table 2:** The mouse and organoid model of stromal BMP signalling. ^13^ **against the human spatial measurements made here.** Their evidence is labelled by system. Ours is from four Visium HD sections (three discovery, one held-out patient) unless a count of eight sections is given, which includes the four Xenium sections for genes on those panels; ISLR and FOXL1 are on no Xenium panel. Every spatial contrast carries a 400 µm block bootstrap and a 400 µm block-permutation null of the fibroblast state labels; bulk estimates are DerSimonian–Laird pools over TCGA, GSE39582, GSE17536 and GSE17537 (1,319 patients), adjusted for stage and age (+ MMR where available).

| Prior claim | Their evidence (system) | This study, in human tissue | Verdict |
|---|---|---|---|
| GREM1 and ISLR are CAF-specific | sorted-cell microarray, ISH (human) | GREM1 18.7, ISLR 7.8–9.4 CP10k in CAF/fibroblast states | CONFIRMED |
| ISLR<sup>+</sup> CAFs lie closer to cancer cells than GREM1<sup>+</sup> | ISH distance, 7 patients (human) | GREM1-type median 66–341 µm vs BMP-type 15–62 and ISLR-type 29–97; 8/8 sections | CONFIRMED and refined |
| GREM1 antagonises BMP in the tumour | reporter, pSMAD1/5, tumoroids (mouse line) | tumour ID1 beside GREM1-type fibroblasts, held-out odds ratio 0.47 [0.33–0.67]; SMAD5 activity −5.1/ − 5.5 | CONFIRMED in situ |
| Bulk GREM1-high is adverse | 53 rectal ISH, 556 colon array (human) | raw GREM1: top tertile 1.45 [1.06–1.97]; per s.d. 1.22 [1.00–1.47] | REPLICATED |
| GREM1<sup>+</sup> and ISLR<sup>+</sup> CAFs are distinct, inversely related populations | dual smFISH, scRNA-seq (mouse + human) | co-detection odds ratio 1.00–1.38 (4/4, at or above chance); ISLR does not separate the states (*q* = 0.59) | NOT REPRODUCED |
| ISLR potentiates BMP in neighbouring cells | BMP7 + conditioned medium, AAV8-*Islr* (mouse) | tumour ID1 beside ISLR-type *lower* than beside BMP-type (0.32–0.65, 4/4); bulk *ρ* with ID1–3 −0.06 to −0.10 (4/4 negative) | CONTRADICTED |
| TGF-*β*–FOXL1 drives *Grem1* at the expense of *Islr* | overexpression, CRISPR, ChIP (mouse line) | FOXL1 lower in GREM1-type (4/4), PDGFRA lower (8/8), TGFB1 lower (4/5), while TGF-*β*/SMAD3 activity is higher | CONTRADICTED (gene assignment) |
| GREM1<sup>+</sup> CAFs are *α*SMA<sup>+</sup> myofibroblastic | smFISH + IF, gel contraction (mouse + human) | ACTA2/TAGLN mixed in sign across sections; Hallmark myogenesis higher | INCONCLUSIVE |
| GREM1 antagonism releases Lgr5<sup>+</sup> stemness and blocks *Krt20* | AP vs APS tumoroids, GREM1-nAb (mouse) | stem set 0.72–1.28 (8 sections, no direction); KRT20 0.33–1.67 (2/5 beyond null) | NOT REPRODUCED |
| ISLR-high is favourable; the ISLR − GREM1 balance drives outcome | 53 rectal ISH, 556 colon array (human) | ISLR-high adverse 1.36 [1.05–1.76]; ISLR-low null 1.00 [0.80–1.25]; balance 0.99 [0.88–1.10]; ISLR-low adverse 1.61 [1.08–2.41] only after subtype adjustment | CONTRADICTED (one-sided) |
| Both highest in stroma-rich/CMS4 | TCGA, microarray (human) | CIT C4: 83% GREM1-high, 0% ISLR-low; GREM1–ISLR *ρ* +0.64 to +0.74 | CONFIRMED; consistent with the one-sidedness |

What this changes for human prognosis is the interpretation rather than the direction of the GREM1 signal. In these tumours the GREM1 state is a TGF-*β*/SMAD3-driven, EMT-high, non-proliferating fibroblast state that forms a palisade set back from the invasive front, beside tumour cells that have lost BMP-target output and are themselves less proliferative and more EMT-like. This may explain why a GREM1 score is prognostic, why it is not independent of CMS4, and why both GREM1 and ISLR—two stromal genes—look adverse in bulk: on this interpretation, what is being measured is the extent of that front. It also says which readouts are informative. Lgr5 and Ki-67, the readouts of the tumoroid experiments, are not the human correlates of GREM1 antagonism: stemness is flat and proliferation falls. Tumour ID1 and SMAD5 activity in the cells adjacent to GREM1^+^ fibroblasts are, and they are measurable in tissue. For the therapeutic arms, the implication is asymmetric: GREM1 neutralisation keeps a human-anchored rationale and a pharmacodynamic readout, with SMAD4/18q status as the stratifier suggested by the mouse tumoroid work, while adding an ISLR-type potentiator is harder to justify from these data, since the human front is short of BMP ligand rather than short of a potentiator of it, and ISLR carries neither the spatial nor the prognostic behaviour the model assigns to it. These remain observational findings in four Visium HD sections and four bulk cohorts, with the limits set out below; they say what the human tumour looks like, not what an intervention would do.

### A spatial reference for the human colorectal cancer front

These maps are specific to human colorectal cancer, and that is where they add most. Much of what is known about the stromal BMP gradient comes from the normal mouse intestine ^11^, and much of the mechanism of the GREM1 axis from mouse cells, tumoroids and mouse in vivo models ^13^. The human tumours keep the organisation of the normal mouse colon—FOXL1^+^PDGFRA^+^ telocyte-like fibroblasts make BMP ligands and a GREM1-producing state antagonises them—but differ from the mouse tumour model in several respects (Table 2): FOXL1 marks the BMP-producing state rather than the GREM1 state; ISLR marks no opposing fibroblast population and does not track tumour BMP output; and tumour cells beside GREM1-type fibroblasts lose BMP-target output without gaining stemness markers. These differences became visible because cell states and positions were measured in patient tissue, several of them relative to a border that no expression value defined. Alongside existing single-cell and spatial atlases of colorectal cancer ^6–10^, this work therefore adds a cell-resolved map of the human invasive front that is anchored to the H&E pathologists already read, states the level of evidence for every finding, and makes the cell boundaries and corrected expression profiles available on request (Code availability), so that findings from mouse and organoid models can be checked against human tissue before they are carried into biomarker or drug studies.

### Geometry and topology: value and limits

Treating the front as a shape rather than a line added information that zone averages miss. The signed distance makes every comparison relative to a border that no expression value defined; change surfaces showed that the tumour interior is not homogeneous but changes state abruptly where stroma intrudes, a result validated in all seven validation sections; and the curvature and topology of the map placed normal epithelium at convex stretches of the front and CD4 T cells in enclosed stromal pockets. Each operator states its scale (50 µm smoothing, 16 µm state pixels, 400 µm null blocks) and was tested against rigid shifts or block nulls rather than label permutations. The approach also has clear limits. All four operators inherit the quality and the threshold of the tumour map (*P ≥* 0.4), and their outputs move with the smoothing scale and the regularisation *λ*, which is why stability ranges are reported. Sections are two-dimensional cuts of a three-dimensional tumour, so an enclosed pocket or an island may be the cross-section of an invagination or of a bud connected out of plane. Curvature is a second derivative and is noisy on an 8 µm raster, and the convex-front result was supported but not validated (4/7 and 5/7 sections). Odds ratios at surfaces are inflated where 8 µm bins mix neighbouring cells (Cervilla Visium HD, 32.7), so the Xenium values give the more conservative estimate. Pre-registered links between front shape and CAF state, GREM1 or tumour budding failed, and geodesic, anisotropic and barrier metrics worked only in simulation. Here, geometry and topology locate where to look; they are not stand-alone biomarkers.

### Therapeutic hypotheses

These data are observational, and every implication is a hypothesis for experimental testing. (i) *GREM1 neutralisation.* A neutralising antibody restored BMP targets in mouse Smad4-intact, but not Smad4-mutant, tumoroids ^13^. Our three tumours lack 18q loss, and tumour ID1 and SMAD5 activity beside GREM1-type fibroblasts are candidate pharmacodynamic readouts. Because the stemness arm was refuted, we would not expect GREM1 blockade to act by reducing stemness. (ii) *BMP restoration.* In mice, AAV8-delivered *Islr* raised BMP signalling and prolonged survival ^13^; in these human tumours ISLR neither marks the counter-state nor tracks tumour BMP output, so restoration would have to supply ligand rather than potentiate it. BMP agonists such as a BMP7 variant differentiated chemoresistant CRC stem-like cells ^31^, but BMP activity marks poor-prognosis mesenchymal CRC ^4^, the BMP-type score was not protective, and in GSE39582 it was univariably adverse for relapse; restoration would need defined contexts. Stromal Hedgehog activation lowers Grem1 and restrains tumours in mice ^32^, and the bed fibroblasts carry a Hedgehog-responsive programme, an indirect route. (iii) *IL-1/IL1R1 blockade.* IL1R1^+^ CAFs drive tumour growth and immunosuppression in CRC, and fibroblast-specific *Il1r1* deletion or anakinra reduced tumour growth in mouse models ^33^; anakinra with fluorouracil and bevacizumab was tolerated in a single-arm phase 2 study ^34^. Our niche-scale IL-1*→*iCAF coupling supports a spatial readout for such studies, with the caveat that its fibroblast-specific replication was partial. (iv) *LTβR and TLS induction.* The LT*β →*CCL19 axis is the lymphoid-organiser programme of TLS neogenesis, and TLS maturity is associated with outcome in CRC ^35–37^; in these pMMR-like tumours, CD8 T cells and TLS-like aggregates sit at the margin, not in the core. Whether LT*β*R agonism would move them inward is untested here, and the axis failed in the held-out patient. Front-CAF genes (COL11A1, MMP11, INHBA) lie on the FAP/CMS4 axis, where FAPI-PET imaging is more realistic in the near term than therapy ^38^.

### Pre-registered failures

Several tests failed, and we report them as results (Supplementary Fig. 11; Supplementary Table 2): the GEO primary for the BMP balance (0.88, 0.74–1.05); independence of the GREM1 score from molecular subtype; the stemness arm (opposite direction on Xenium 5K); proliferation at tumour-state fronts on an external patient; front shape, budding and clone–niche tests; the gene-level held-out replication of both state transcriptomes, which left the locked marker panel empty and its survival test unrun; cell-resolution attribution of every signalling pathway; the LT*β* axis in the held-out patient; the detector on faded post-Xenium H&E; geodesic, anisotropic and barrier metrics that worked only in simulation; and H&E map features as prognostic markers in TCGA. Two signals produced by our own exploratory methods, a CD8 immune-access index and a Schrödinger-bridge model of gradual state change between zones, were reproducible but were matched by control maps or depended on a tuning parameter, and were withdrawn (Supplementary Fig. 11, rows N12 and N23).

### Limitations

Although the spatial discovery cohort comprised three primary colorectal tumours, the principal findings were evaluated across complementary spatial transcriptomic platforms and three additional patients, with clinical associations further examined in 1,319 patients across four bulk transcriptomic cohorts. The discovery tumours represent stroma-low, pMMR-like, chromosomally unstable colorectal cancers, with MMR status inferred from transcriptomic profiles. Despite consistent replication of the BMP-to-GREM1 fibroblast switch across six evaluable validation sections, substantial inter-section heterogeneity (*I*^2^ = 0.98) warrants investigation in larger spatial cohorts encompassing diverse molecular and histopathological subtypes.

Differences in gene coverage, spatial resolution and histological image quality limited selected cross-platform comparisons. In particular, independent spatial evaluation of ID1, ISLR and FOXL1 was restricted to a single held-out Visium HD section, and external sections were analysed using RNA-based tumour masks. Pathway activity analyses lacked spatial null models, and niche-scale signalling analyses were specified following the unsuccessful cell-resolution tests. The bulk GREM1-associated survival findings should also be interpreted in the context of molecular subtype and tissue composition.

Analytical procedures were documented in a time-stamped internal record rather than a public registry. Although OmiCore-Seg and OmiCoreCloud are proprietary, their processed outputs and relevant downstream analytical scripts are available upon reasonable request to facilitate independent verification.

Finally, the identified spatial transcriptional associations provide a human tissue-based framework for investigating stromal BMP signalling rather than direct evidence of causal interactions. Complementary protein-level validation, in situ hybridisation and functional studies using SMAD4-stratified fibroblast–organoid co-cultures will be important for establishing the mechanistic and therapeutic relevance of these findings.

## Methods

### Data

Discovery: Visium HD sections (Cancer P1, P2, P5; Normal P3, P5), Xenium serial sections (P1, P2, P5; 422 genes after filtering) and the Chromium Flex reference (8 samples, 260,506 cells, 38 level-2 states) from Oliveira et al. ^22^ (GEO GSE280318, GSE280314). Validation: Cervilla CRC Visium HD (Zenodo 17999961; 8 µm bins with *≥*100 UMI) and Xenium 5K of the same patient (Zenodo 18000256; 307,235 cells, 5,001 genes) ^26^; 10x Genomics Xenium colon-cancer (560,594 cells, 425 genes) and CRC immuno-oncology (366,425 cells, 480 genes) sections. Bulk: TCGA COAD and READ STAR-TPM, clinical data and TCGA-CDR endpoints ^39^ (621 patients, 578 with complete covariates; 599 diagnostic slides); GEO GSE39582^27^ (566 tumours), GSE17536 (177) and GSE17537 (55) (GPL570 gene-level log_2_ expression).

### Comparison with the prior stromal BMP model

Fibroblast states, matrices, maps, signed distances and nulls are those of the main analysis. A third state was added for this comparison only: ISLR-type, a confident fibroblast with ISLR detected and GREM1 not, matching the mutually exclusive ISLR^+^ CAF of the prior model ^13^. Ownership was taken from the Flex reference as mean CP10k per level-2 state (up to 4,000 cells per state). For every gene and state pair, the detection rate in each state was compared as a log_2_ rate ratio with a 400 µm block bootstrap (500 draws) and a null from 400 µm block permutation of the fibroblast state labels (200 draws), one permutation serving all contrasts of a section. Distances were the Euclidean distance from each fibroblast to the nearest confident tumour cell, reported for all stroma and within 250 µm of the border, with block-bootstrapped medians. Tumour readouts used the design of the niche test (see Niche test, below) unchanged (interior tumour cells, *d < −*50 µm; fibroblast neighbours within 30 µm of one state only; Mantel–Haenszel odds ratios over 25 µm distance bins *×* neighbour count; block bootstrap and block-permuted state null) extended to the three state pairs and to ID1–3, SMAD7, the stem set, KRT20 and MKI67. In bulk, GREM1 and ISLR were scored as gene z of log expression, both raw—as the prior work measured them—and residualised on fibroblast abundance; tertile groups reproduced their GREM1-high and ISLR-low definitions; Cox models and DerSimonian–Laird pooling followed the bulk survival analysis (see Bulk scores and survival, below), with CIT molecular subtype added where available (GSE39582); CMS labels were not used for TCGA, so subtype adjustment was limited to GSE39582. Code is available as described under Code availability.

### Segmentation (OmiCoreSeg)

Nuclei and cells were segmented with OmiCoreSeg (version 0.2.0), in-house segmentation software developed by OmiCore Inc. (Fukuoka, Japan). OmiCoreSeg and its underlying technologies are the property of OmiCore Inc.; its implementation is not described here. It takes the full-resolution H&E image only (0.274 µm/px) and never sees gene expression. It returns nucleus and cell boundaries; every in-tissue 2 µm Visium HD bin was assigned to the cell containing its centre, so that the count of gene *g* in cell *i* is *x_ig_*= Σ*_b_*_∈_*_i_ y_bg_* over the bins *b* whose centres fall in the cell. Benchmark: PanNuke colon test folds ^24^ (962 patches, 16 mosaics, 23,550 nuclei). Predicted and annotated nuclei were matched one-to-one at intersection over union (IoU) *>* 0.5, and panoptic quality was

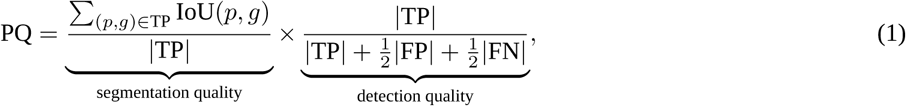

which rewards both finding nuclei (TP, true positives; FP, false positives; FN, missed nuclei) and outlining them accurately; 95% CIs came from 2,000 patch bootstraps. Open-source comparators were run on the unmodified H&E with their published models: InstanSeg ^40^, Cellpose-SAM ^41^, StarDist ^42^ (default and tuned), QuPath ^43^, scikit-image ^44^ and OpenCV ^45^ watershed pipelines, and CellProfiler ^46^. Settings were fixed on mosaics 00–07 and PQ is reported on both halves (0.539 and 0.537; Supplementary Table 1). Cells were typed with RCTD ^23^ (doublet mode) against the Flex reference; a cell was confident when RCTD called it a singlet or a certain doublet.

### Tumour map, signed distance and zones (OmiCoreTumorDetector)

The released model ^21^ is an ensemble of *M* = 3 convolutional classifiers trained on NCT-CRC-HE-100K ^25^, with Macenko normalisation ^47^ for one member. Each member *m* scores 224-pixel H&E tiles at 0.5 µm/px (112-pixel stride) and returns nine class probabilities *p_m,c_*(*x*). The tumour probability at position *x* is the ensemble mean, and the tumour is its superlevel set:

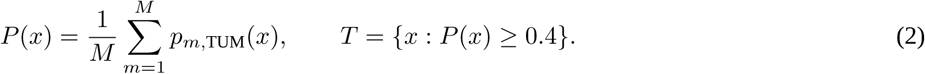

The map is computed from the image alone, so no expression value can influence where the tumour is. Every cell then receives its signed distance to the tumour border *∂T* (computed on an 8 µm raster; glass is not treated as a border):

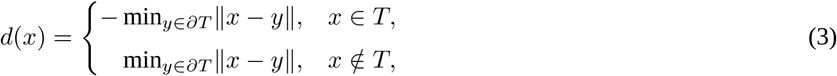

so that *d <* 0 inside the tumour, *d* = 0 on the H&E border and *d >* 0 in stroma. Compartments are intervals of *d*: tumour bed (*d < −*50 µm), front (*−*50 *< d ≤* 150 µm), mid stroma (150 *< d ≤* 500 µm) and far stroma (*d >* 500 µm); near (50 *< d ≤* 150 µm) versus distant (*d ≥* 250 µm) stroma for CAF zonation. Validation truths were RCTD and Xenium tumour cells, the pathologist annotation of section P2 (Zenodo 11402686; 16 µm bins) and ABSOLUTE purity ^48^ in TCGA. Off-domain recalibration replaced the fixed 0.4 by the crossing point of a per-slide two-component Gaussian mixture fitted to logit *P*.

For comparison we used two expression-defined tumours. The RNA mask (used for all external sections) is the smoothed share of tumour cells among confident cells,

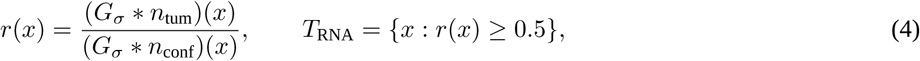

where *n*_tum_ and *n*_conf_ count tumour and confident cells on a 20 µm grid and *G_σ_* is a Gaussian kernel with *σ* = 100 µm (50 µm as a sensitivity analysis). The nearest-neighbour region assigns cell *i* to the tumour when at least 60% of its 20 nearest typed cells are tumour cells.

### OmiCoreCloud

OmiCoreCloud (OmiCore Inc.; access terms under Code availability) is described here by its principle, inputs and outputs, and was evaluated against planted ground truth. Principle: each cell is represented as a probability cloud over reference states, the cell’s own RNA is separated from RNA received from its neighbourhood, and expression is denoised in a depth-aware way; the implementation is proprietary. Inputs: OmiCoreSeg cell *×* gene counts, cell coordinates, the 38 Flex reference states and an ambient prior. Outputs per cell: a decontaminated profile (mode A), a denoised profile (mode B), state probabilities *π_ik_* over reference states *k*, the state entropy *H_i_*= *−* Σ*_k_ π_ik_* log *π_ik_* (low when the state is unambiguous) and an atlas-misfit score (enthalpy; high when no reference state explains the cell). Benchmark: Flex cells of a held-out patient were placed on real P1 geometry at real depth and contamination, so the true profile 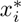 of every cell is known. On the test half, fidelity was the Pearson correlation between log(1 + *x̂_ig_*) and 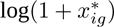, and foreign-lineage leak was the share of counts on genes owned by another lineage; depth and ambient sweeps and leave-one-sample-out runs are in Supplementary Fig. 6. Apart from the OmiCoreCloud arm of the tool ablation and the supportive atlas-misfit readout, no result in this paper was computed on its output; the headline analyses use the open cell-level deconvolution below.

### Spill kernel

RNA that a cell releases can be captured in a neighbouring cell. We measured this directly with genes that only one lineage expresses. For lineage *L*, a gene *g* is owned by *L* when its Flex expression is *≥*20 counts per million and *≥*3-fold above every other lineage; any count of *g* in a cell of another type must then have come from outside that cell. On an 8 µm raster with pixels *p*, let *y_g_*(*p*) be the counts of *g* in confident non-*L* cells, *Q_g_*(*p*) the counts of *g* in confident *L* cells (the source) and *e*(*p*) the exposure, the number of 2 µm bins covered by non-*L* cells (16 for a fully occupied pixel). The counts were modelled as Poisson with mean

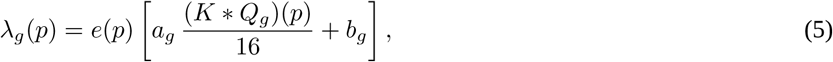

where *∗* is two-dimensional convolution, *a_g_ ≥* 0 is the leak amplitude of gene *g* (a relative amplitude, not a fraction), *b_g_ ≥* 0 a spatially flat background, and the kernel is a mixture of a short and a long isotropic Gaussian,

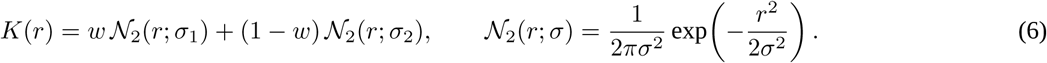

For each kernel shape *θ* = (*w, σ*_1_*, σ*_2_) on a grid (*σ*_1_ 4–24 µm, *σ*_2_ 16–300 µm), the non-negative amplitudes were fitted by projected Newton steps on the Poisson log-likelihood

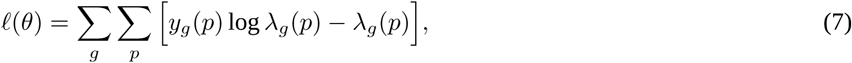

and *θ*^^^ = arg max*_θ_ ℓ*(*θ*). The short width *σ*_1_ is the distance over which RNA is mis-assigned between neighbours; a diffuse background would instead appear as a large weight on a wide *σ*_2_ or as a large *b_g_*. Controls were a planted dry test with known *θ*, a 1 mm shift of the source *Q_g_*(which should remove the likelihood gain if the signal is spatially specific) and a per-gene consistency test.

### Cell-level deconvolution

The fitted kernel defines how much RNA each cell receives from neighbours of a different type. Writing *o_i_*for the cell’s own counts (per gene), the observed counts are

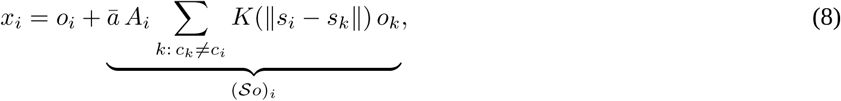

with *s_i_*the cell position, *A_i_*its area, *c_i_*its type and *a̅* the median leak amplitude of the section. Only cells of a different type contribute, because RNA exchanged between cells of the same type does not change a cell-type profile. The own counts were recovered by the non-negative fixed-point iteration

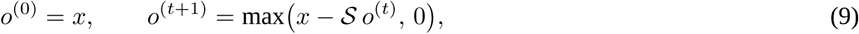

which subtracts the estimated received RNA while never producing negative counts. The number of iterations was chosen on held-out owned genes: *t*^∗^ = arg min*_t_*_∈{0,2,5,10,20,30,50}_ *F* (*t*) subject to *R*(*t*) *≥* 0.90, where *F* (*t*) is the remaining foreign signal (owned-gene counts in non-owner cells, relative to raw) and *R*(*t*) the retained counts in owner cells. A pre-registered grid Richardson–Lucy iteration failed its rule and was replaced (a recorded deviation) before any headline was re-run.

### Spatial nulls

Neighbouring tissue is correlated, so permuting labels cell by cell makes almost any zone contrast look significant. We therefore used a block-permutation null. The section is cut into 400 *×* 400 µm blocks *B*_1_*, …, B_m_*; a null map is made by drawing a random permutation *π* of the blocks and giving each position *x ∈ B_b_* the zone, distance or state label of the corresponding position in block *B_π_*_(*b*)_. This keeps the spatial structure inside each block but breaks its relation to the true tumour border. A statistic Δ passes when it exceeds the 97.5% quantile of its values on the null maps (20 maps for zone contrasts, 200 for niche tests).

### Multiscale interface wavelet (MIW)

For a per-cell signal *v* (for example, the BMP balance of fibroblasts or the expression of a CAF gene) and a scale *s*, the MIW coefficient contrasts the band just outside the border with the band just inside it:

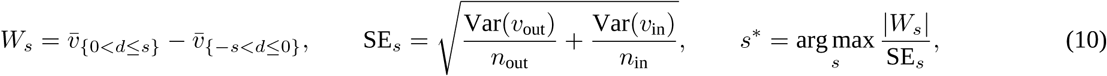

for *s* ∈ {25, 50, 100, 200, 400} µm. The characteristic scale *s*^∗^ is the band width at which the signal changes most reliably across the border: a contamination artefact peaks at one cell diameter, whereas a tissue programme peaks at the scale over which it is organised. *|W_s∗_ |* was compared with the block null.

### Change surfaces (SCPSD)

Tumour-cell state, the first ten principal components of tumour-owned genes, was rasterised at 16 µm into a vector field *S* : Ω *→* R^10^ over the tumour pixels Ω and denoised by total variation,

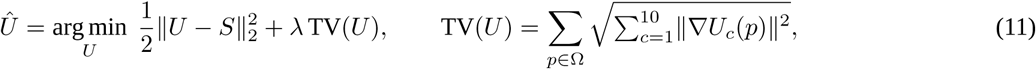

with *λ* = 0.1 (0.05 and 0.2 for stability). The TV penalty favours piecewise-constant solutions: it removes pixel noise but keeps sharp boundaries between tumour states. The change surface is the set of pixels with the steepest denoised gradient, 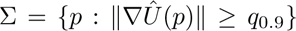, where *q*_0.9_ is the 90th percentile over Ω. With *I* the pixels containing stromal intrusions (non-tumour cells inside the tumour mask), enrichment of surfaces at intrusions was the odds ratio

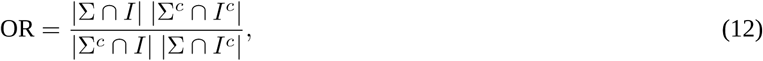

compared with the same odds ratio after 1 mm rigid shifts of the state map and after shuffling states among tumour cells; a section passed when its OR exceeded the larger control. Across *n* sections with *k* passes the verdict used the exact one-sided sign test 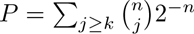. What sits at the surfaces was tested by Mantel–Haenszel odds ratios (equation 19), surface versus non-surface interior tumour cells, stratified by 25 µm distance bins.

### Curvature and topology of the front

The curvature of the H&E front is the divergence of the unit normal of the smoothed signed distance,

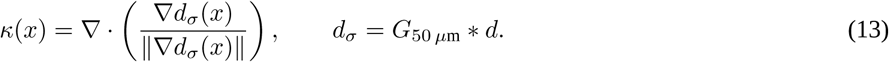

Because *∇d* points out of the tumour, *κ >* 0 where the tumour bulges into stroma (convex) and *κ <* 0 where stroma bulges into the tumour (concave); the top and bottom terciles of *κ* along the border of each section defined convex and concave stretches. Topology used the superlevel sets of the tumour probability,

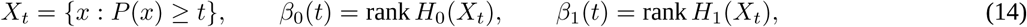

as *t* decreases from 1 to 0. *β*_0_ counts connected tumour islands and *β*_1_ counts enclosed holes, which in the plane are the bounded components of the stroma (Alexander duality), that is, stromal pockets surrounded by tumour. Cubical persistent homology records the threshold at which each island or pocket appears and disappears; classes that persist over a wide range of *t* are structures of the tissue rather than noise in *P*. Barrier and access metrics were exploratory: they were scored against 509 simulated tissues with known ground truth and did not transfer to real tissue (Supplementary Fig. 11, rows N9–N11).

### Fibroblast states, BMP balance, anatomy control and CAF zonation

BMP-type fibroblasts carry *≥*1 BMP4 or BMP5 transcript and no GREM1; GREM1-type fibroblasts carry *≥*1 GREM1 transcript and no BMP4 or BMP5. For a fibroblast pseudobulk with expression *c_g_* (counts per 10,000), the BMP balance and its bed-to-front contrast are

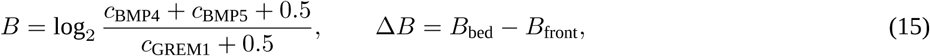

so that Δ*B >* 0 means that ligand production gives way to the antagonist towards the front; the pseudocount 0.5 keeps *B* finite. Δ*B* passed when it was positive and above the null 97.5% quantile. Xenium panels without BMP5 used BMP4 only. Anatomy control: each fibroblast took the nine-class probabilities of the nearest whole-section detector tile; fibroblasts in tiles with *p*_MUS_ *≥* 0.2 or *p*_NORM_ *≥* 0.2 were removed; the test passed if Δ*B >* 0 in 3/3 HD sections and above the null in *≥*2/3, and replicated if positive on Xenium 3/3. CAF zonation compared fibroblast log_2_ fold changes between near and distant stroma with an independent measurement over the gene set *G* with *≥*20 counts in both zones,

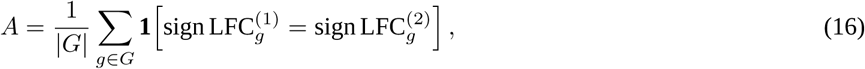

passing at *A ≥* 0.70 and above the block-null 97.5% quantile. Neighbourhood enrichment used 50 within-compartment label permutations.

### State and niche transcriptomes and pathway activity

Fibroblast states (whole section, confident fibroblasts) and interior tumour cells (*d < −*50 µm) in GREM1 versus BMP niches were aggregated to pseudobulk counts per section and group on the spill-deconvolved matrix. With *y_gj_*the voom log-CPM of gene *g* in pseudobulk *j* (with precision weights), *z_j_*the group indicator and section as a block, limma ^29,30^ fitted

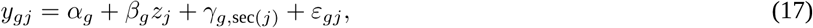

and tested *β_g_*(the log_2_ fold change) with an empirical-Bayes moderated *t*. Defining genes (BMP4, BMP5, GREM1) were excluded. Fibroblast analyses removed genes whose Flex mean in any non-fibroblast lineage was *≥*2-fold the fibroblast mean; tumour analyses kept genes with a Flex tumour mean *≥*3-fold the fibroblast mean. A discovery hit required BH *q <* 0.05, *|* log_2_ FC*| ≥* 1 and the same sign in 3/3 per-section pseudobulk fold changes. Held-out test (Cervilla Xenium 5K, same state and niche definitions, RNA-mask distance): sign agreement (equation 16) over hits with *≥*20 counts in both groups, required to be *≥*0.75 and above the 97.5% quantile of 200 within-400 µm-block permutations of state labels. Oliveira Xenium agreement was reported as a secondary check. Pathway and TF activity were estimated by the univariate linear model of decoupler ^49^: for each source *s* (a pathway or transcription factor) with target weights *w_sg_*, the moderated statistics were regressed on the weights,

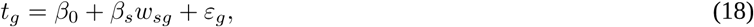

and the activity of *s* is the *t* statistic of *β*^^^*_s_* (positive when the targets of *s* are up in GREM1-type fibroblasts or GREM1-niche tumour cells). Resources were PROGENy ^50^ (top 500 genes), CollecTRI ^51^ and MSigDB Hallmark; activities with BH *q <* 0.05 in discovery were called replicated if they had the same sign on 5K Welch *t* statistics computed the same way. The BMP target set (ID1, ID2, ID3, SMAD6, SMAD7, MSX2, BAMBI) was the pre-stated positive control. The locked panel admitted only replicated genes; it was empty, so its survival test was not run.

### Niche test

Units were confident interior tumour cells (*d < −*50 µm; tumour bins on Cervilla HD) with *≥*1 confident fibroblast within 30 µm. GREM1-niche units had *≥*1 GREM1-type and no BMP-type fibroblast within 30 µm, BMP-niche units the reverse. Readouts were detection of any stem gene (LGR5, ASCL2, AXIN2, RNF43; primary), ID1 (secondary), and ID2, ID3, SMAD6, SMAD7 and MSX2. Detection depends on depth and on position, so niches were compared within strata *k* (25 µm distance bins *×* fibroblast-neighbour count 1, 2, *≥*3) by the Mantel–Haenszel odds ratio

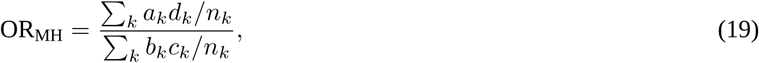

where, in stratum *k*, *a_k_* and *b_k_* are GREM1-niche cells with and without the readout, *c_k_* and *d_k_* the same for BMP-niche cells, and *n_k_*= *a_k_* + *b_k_* + *c_k_*+ *d_k_*. OR_MH_ *<* 1 means the readout is less often detected beside GREM1-type fibroblasts. Depth tertiles were a sensitivity analysis; intervals came from 500 block-bootstrap resamples and the null from 200 block permutations of fibroblast states. The ID1 confirmation rule (odds ratio *<* 1, upper limit *<* 1, below the null 2.5% quantile) was recorded before the held-out run.

### Cell–cell communication

#### Contact test

LIANA consensus pairs ^19^ were kept if directional by Flex ownership (*≥*10 counts per million and *≥*3-fold for the ligand in the sender and every receptor subunit in the receiver), so that RNA spilled between partners could not create either signal. For an ordered sender–receiver type pair and the *n_D_*cell pairs in a distance class *D*, co-expression was measured as observed over expected,

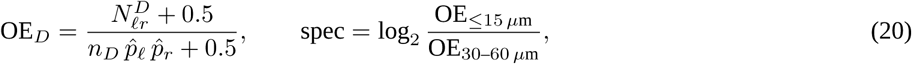

where 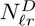 counts pairs with the ligand detected in the sender and the receptor in the receiver and *p̂_ℓ_*, *p̂_r_* are the detection rates in the same pairs. Touching cells (*≤*15 µm) and near but non-touching cells (30–60 µm) share depth and microenvironment, so their ratio isolates what is specific to contact. A pair was contact-specific above the 95th percentile of 500 random directional gene pairs on *≥*2 of 3 slides; power used planted receptor inductions on the real contact graphs. *Zone-resolved LIANA.* LIANA (Python, version 1.10) `rank_aggregate` with the consensus resource on the spill matrix (log1p CP10k), groups at level 1 with fibroblasts split into BMP-type, GREM1-type and other (groups *<*30 cells dropped), run per section in bed (*d < −*50), front (*−*50 to +150) and distant stroma (*>* 500 µm). Front-specific: magnitude rank *≤*0.05 at the front in 3/3 sections and not *≤*0.05 in the bed (or a fourfold better rank). Null: the same run on 10 block-permuted zone maps; the call count must exceed the null 95th percentile, and calls occurring in *≥*10% of null maps were dropped. Ownership: ligand owned by the sender and receptor by the receiver (*≥*2-fold), otherwise flagged. Held-out test on Xenium 5K with RNA-mask zones, against a 10-map null. *Ligand-field target response (LFTR).* For each pre-registered pathway, every receiver cell *i* with *≥*1 sender within 100 µm received a ligand field

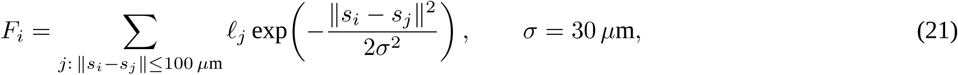

where *ℓ_j_*is the ligand count of sender *j*. *F_i_* summarises how much ligand reaches the receiver from all senders, weighted by distance. High versus low tertiles of *F* were compared within sender-count classes (1, 2–3, 4–7, *≥*8) by Mantel–Haenszel odds ratios (equation 19) of receiver-owned target detection, stratified by sender class, 50 µm distance and receiver-UMI tertile. Nulls: ligand permuted among senders within 400 µm blocks (cell resolution); the field read at receiver positions shifted by 1–2 mm (niche scale; registered after the cell-resolution result); 20 detection-matched random target sets. Coupled: odds ratio *>* 1 in 3/3, beyond both nulls in *≥*2/3, a 15 µm contamination-ring variant agreeing, BH *q <* 0.10. Replicated: the same in at least half of testable validation sections, with *≥*2 sections from *≥*2 levels. A readout-sensitivity test (CCL2 + MMP3; CCL19) was registered before it was run.

### Immune and pathology readouts

With *f*_in_ and *f*_out_ the share of immune cells among confident cells inside the tumour (*d <* 0) and beyond 50 µm outside it (*d >* 50), immune exclusion is

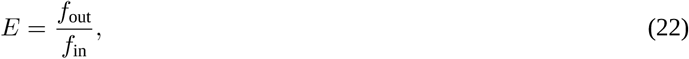

so *E* = 1 means no exclusion and *E* = 5 means immune cells are five times as frequent outside the tumour as inside. Strict NK cells required an NK receptor (KLRF1, NCR1, KLRD1, KLRC1) and a cytotoxic transcript, without CD3, CD8 or myeloid transcripts. Immunoscore-type zones were the core (CT, *d < −*250 µm), the invasive margin (IM, *−*250 *≤ d ≤* 250 µm) and peritumoural stroma (250–1,000 µm). With *ρ_Z_* the density of a population (cells per mm^2^ of occupied tissue on a 25 µm grid) in zone *Z*, the margin-to-core contrast is log_2_(*ρ*_IM_/*ρ*_CT_), with 400 µm block-bootstrap intervals. MMR status was inferred from pseudobulk MLH1, MSH2, MSH6 and PMS2 relative to normal epithelium, with a silencing call at *<* 0.25; arm-level copy number from infercnvpy on 80 µm metacells; iCMS from published markers ^52^. The tumour–stroma ratio inside the tumour envelope is TSR = *A*_STR_/(*A*_STR_ + *A*_TUM_), with *A* the area of detector stroma and tumour classes (stroma-high *>*50% ^28^). Lymphoid aggregates (DBSCAN on B and T cells) were classed as lymphoid aggregate, primary follicle-like or secondary follicle-like by follicular dendritic cell and germinal-centre transcripts. The fetal/revival test compared the front (*−*100 to 0 µm) with the core (*< −*300 µm).

### Meta-analysis

Per-section effects *y_k_*with variances *v_k_*(*k* = 1*, …, K* validation sections) were pooled by DerSimonian–Laird random effects, which allows the true effect to differ between sections:

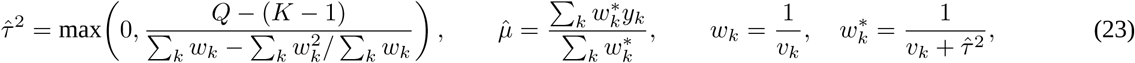

with *Q* = Σ*_k_w_k_*(*y_k_ − y̅_w_*)^2^ and *y̅_w_* the fixed-effect mean. 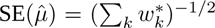; heterogeneity is *I*^2^ = max*{*0, (*Q − K* + 1)/*Q}*, the share of variation due to real between-section differences; the 95% prediction interval 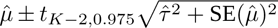 is the range expected for a new section. Results were cross-checked with metafor.

### Bulk scores and survival

A gene-set score is the mean of per-gene z-scores across tumours, 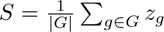. Because a fibroblast-state score also rises with the amount of stroma, every score was residualised on a fibroblast-abundance score *F*_ab_ (COL1A1, COL1A2, DCN, LUM, PDGFRA), *S̃* = *S −* (*α̂* + *β̂F*_ab_), so that it measures the state of fibroblasts rather than their quantity. Survival was modelled by Cox regression,

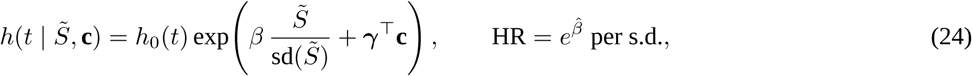

with covariates **c**: stage, age, microsatellite status and site (TCGA), and stage and age (+ MMR in GSE39582) in GEO. Cohort hazard ratios were pooled on the log scale by equation 23. The GEO analysis was pre-registered before the data were opened: primary, the GEO-only pooled hazard ratio of the balance (pass below 1 with the CI excluding 1); secondary, the GREM1-type score above 1. Robustness analyses (leave-one-cohort-out, molecular adjustment, Harrell’s Δ*C*, chemotherapy interaction) and subgroup analyses were registered before running. Coupling was the Spearman correlation *ρ_k_*of the balance with ID1–3, residualised on fibroblast, endothelial, immune and epithelial scores, against 1,000 expression-matched random three-gene sets; cohorts were pooled on Fisher’s *z*, *z̅* = Σ*_k_* (*n_k_ −* 3) atanh *ρ_k_*/Σ*_k_* (*n_k_ −* 3). LFTR bulk coupling residualised ligand and target scores on five composition scores (a logged deviation from the registered adjustment).

### Pre-registration and statistics

Analyses after the initial exploratory phase were written as dated pre-registrations in the analysis record before any run, including hypotheses, readouts, pass rules and validation rules; each was run once, failed tests were not re-tuned, and deviations were recorded (Supplementary Table 2). New methods were first run on planted data. Tests are two-sided unless a direction was pre-registered. Sections are the unit of spatial replication. Analyses were run on a 24-core workstation with 64 GB of memory.

## Data availability

Visium HD, Xenium, post-Xenium H&E and Flex data: GEO GSE280318 and GSE280314^22^. Cervilla CRC Visium HD and Xenium 5K: Zenodo 17999961 and 18000256^26^. Pathologist annotation: Zenodo 11402686. Additional Xenium colon datasets: 10x Genomics (Xenium_V1_Human_Colorectal_Cancer_Addon_FFPE, Xenium_V1_hColon_Cancer_Add_on_FFPE, Xenium_V1_hColon_Non_diseased_Base_FFPE). TCGA expression, clinical data and diagnostic slides: UCSC Xena / GDC; ABSOLUTE purity: GDC PanCancer Atlas. GEO cohorts: GSE39582, GSE17536, GSE17537. Source data are provided with this paper.

## Code availability

OmiCoreTumorDetector is publicly available as open-source software under the Apache-2.0 licence at https://github.com/OmiCore-Japan/omicoretumordetector and can be installed via pip install omicoretumordetector.

All other custom analysis scripts used to generate the results presented in this study, including those for the spill kernel, cell-level inversion, MIW, SCPSD, LFTR and meta-analysis, are available from the corresponding authors upon reasonable request. OmiCoreSeg and OmiCoreCloud are proprietary software developed and owned by OmiCore Inc. Both programs and their underlying technologies remain the intellectual property of OmiCore Inc. and are not publicly available. Access may be considered for non-commercial academic research on request, under terms to be agreed with OmiCore Inc.; any commercial use would require a separate agreement. Requests should be addressed to OmiCore Inc.

The processed outputs of both programs, including nucleus and cell boundaries, bin-to-cell assignments, decontaminated counts, denoised expression profiles, state probabilities, entropy and enthalpy, are available upon reasonable request to facilitate independent verification and reproduction of the downstream analyses. Access to the relevant analysis scripts and processed outputs will be provided to editors and reviewers during peer review.

## Author contributions

S.N. conceived the study, developed the three software tools (OmiCoreSeg, OmiCoreTumorDetector and OmiCoreCloud) and the analytical methods, performed the analyses and wrote the manuscript. J.P.R. contributed to the data analysis and reviewed the manuscript. A.F. performed some of the cell segmentation experiments. G.Y. provided clinical and pathological interpretation, contributed to writing the manuscript and critically reviewed it. All authors approved the final manuscript.

## Supplementary Information

OmiCoreSeg and OmiCoreCloud are proprietary software of OmiCore Inc. and are described here by their inputs, outputs and performance only. OmiCoreTumorDetector is open source (Apache-2.0). Open-source segmenters appear only as benchmark comparators run on unmodified H&E.

**Supplementary Fig. 1:**
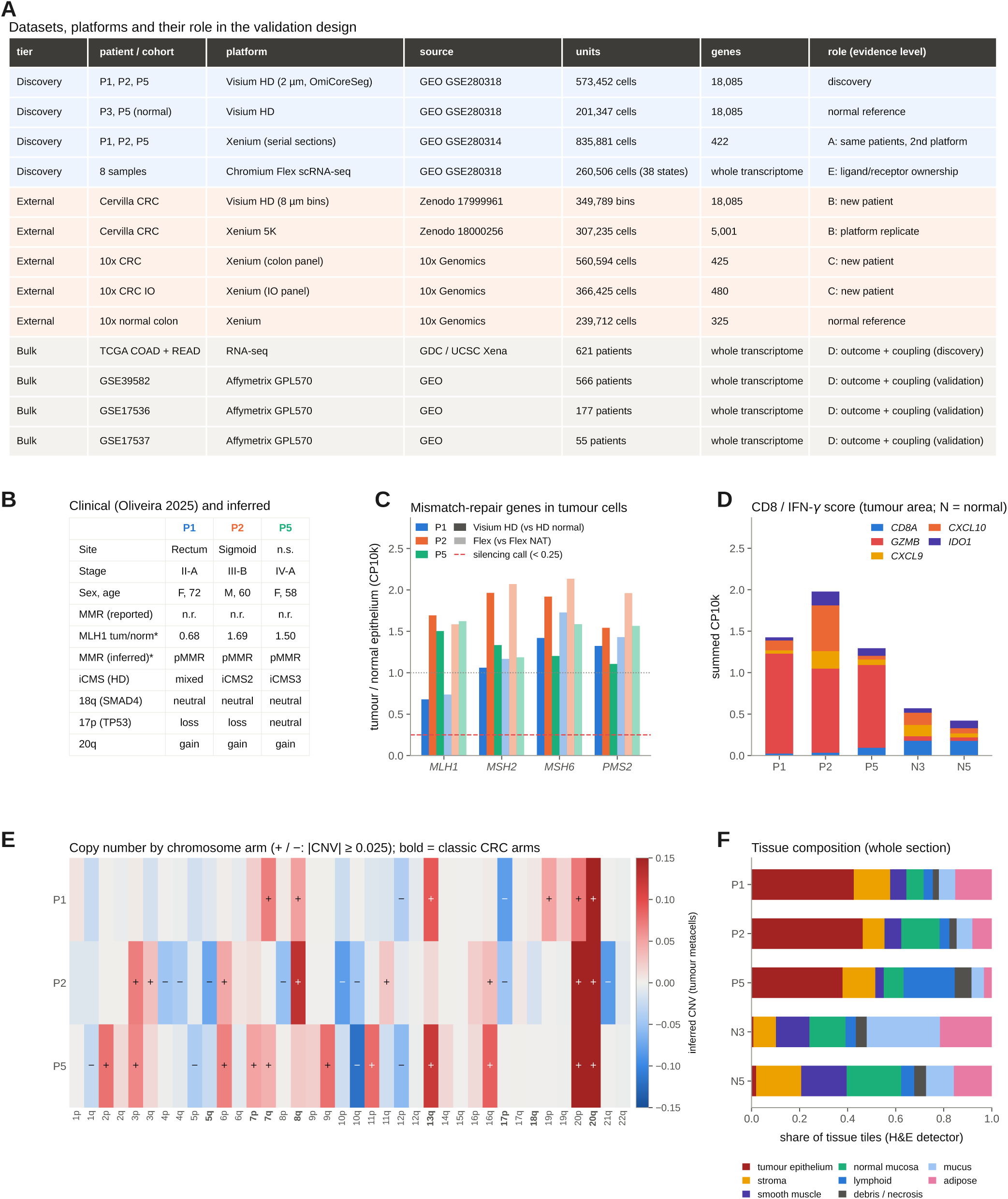
Datasets and the three discovery tumours. **A**, Patients, platforms, units (cells, bins or patients), gene panel size and evidence level of each dataset. The Cervilla Xenium 5K section is the same patient as Cervilla Visium HD (platform replicate, not an additional patient). **B**–**F**, Patient portrait. **B**, Clinical features (Oliveira et al. 2025, Table 1); rows marked * are inferred from expression and are not a diagnosis; n.r., not reported. **C**, MLH1, MSH2, MSH6 and PMS2 in tumour cells relative to normal epithelium (solid, Visium HD; light, Flex); dashed line, pre-registered silencing threshold (0.25). **D**, CD8/IFN-*γ* score (CD8A, GZMB, CXCL9, CXCL10, IDO1; CP10k). **E**, Inferred copy number by chromosome arm in tumour metacells; +/*−*, *|*CNV*| ≥* 0.025; no tumour shows 18q (SMAD4) loss. **F**, Whole-section tissue composition from the detector’s nine classes.

**Supplementary Fig. 2:**
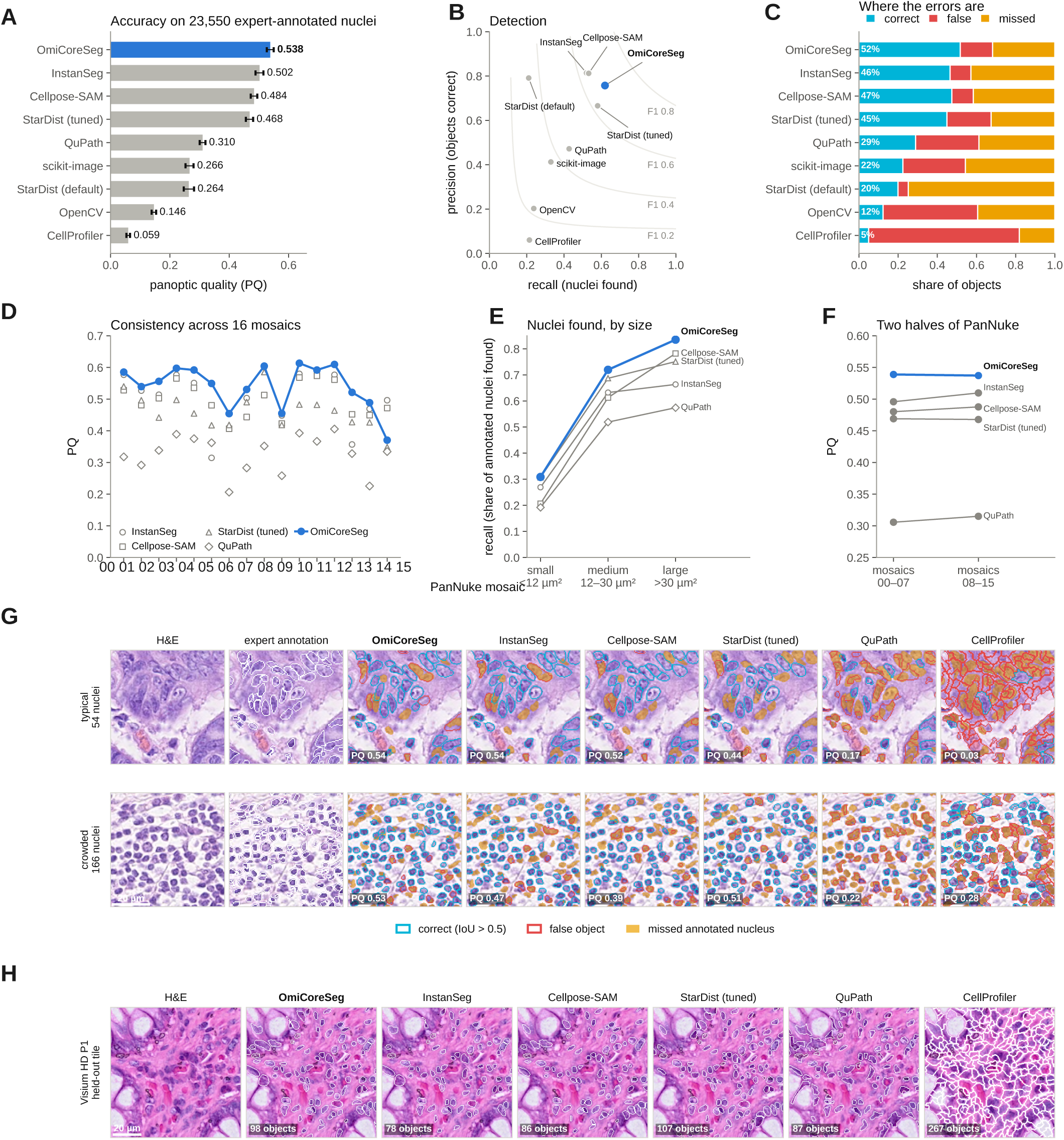
OmiCoreSeg against widely used open-source nucleus segmenters on 23,550 expert-annotated PanNuke colon nuclei (962 patches, 16 mosaics). Every comparator was run on the unmodified H&E with its published model. **A**, Panoptic quality. **B**, Precision versus recall: OmiCoreSeg gains recall (0.619) at lower precision (0.757 versus 0.814 for InstanSeg and 0.812 for Cellpose-SAM). **C**, Outcome of every object. **D**, PQ per mosaic. **E**, Recall by nucleus size. **F**, PQ on the two halves of PanNuke (settings fixed on mosaics 00–07: 0.539 / 0.537). **G**, Example patches chosen by a fixed rule. **H**, Held-out Visium HD tile (qualitative). OmiCoreSeg takes the H&E image as its only input and returns nucleus and cell boundaries. Full metrics in Supplementary Table 1.

**Supplementary Fig. 3:**
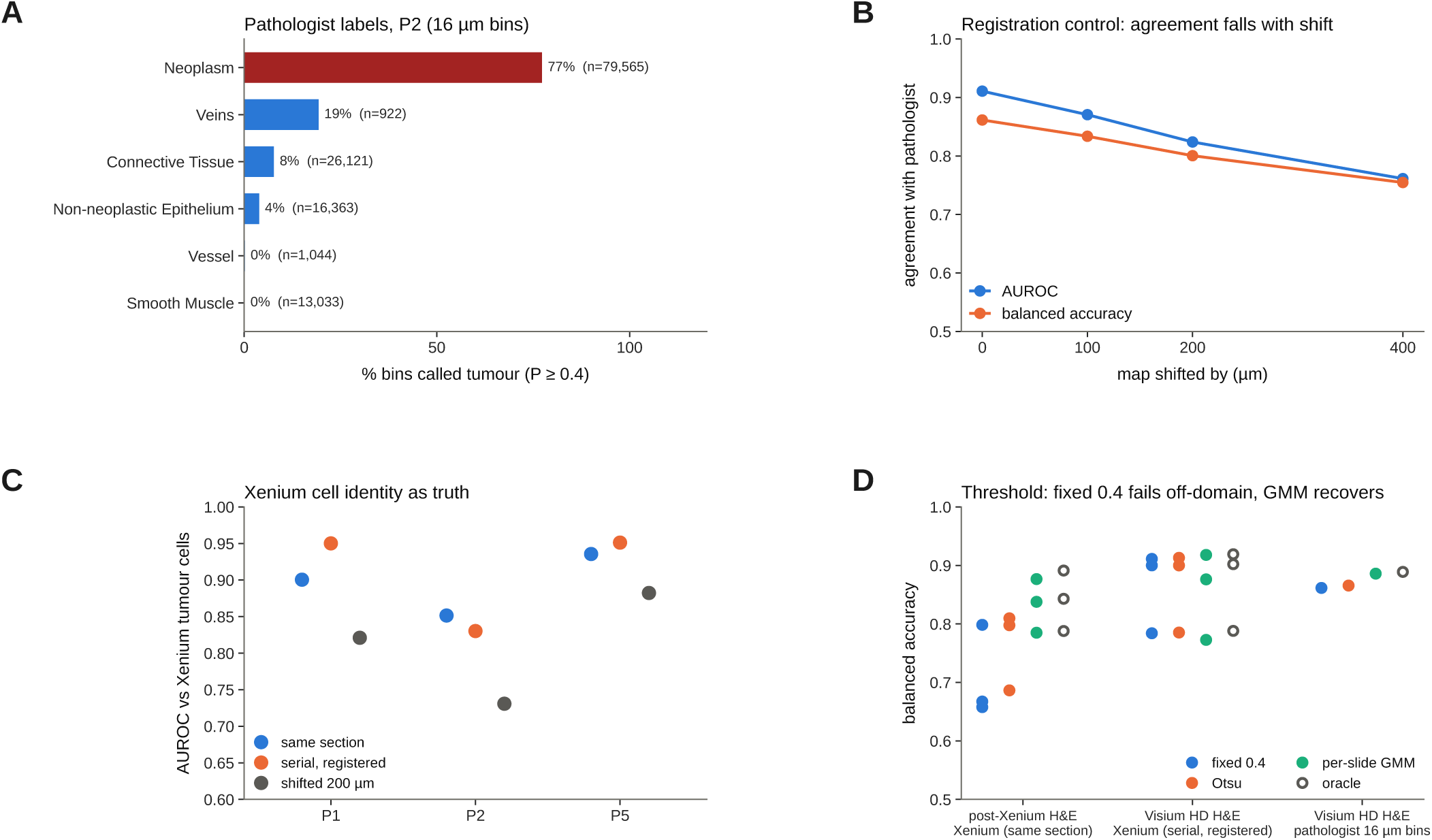
Independent validation of OmiCoreTumorDetector. **A**, Share of bins called tumour (*P ≥* 0.4) by pathologist label on section P2 (10x Genomics annotation, 16 µm bins; AUROC 0.911, specificity 0.950, sensitivity 0.773, *κ* 0.697). **B**, Agreement as the map is shifted by 100–400 µm (0.87, 0.82, 0.76). **C**, AUROC against confident Xenium tumour cells on post-Xenium H&E (0.900/0.851/0.936), on the registered Visium HD serial section (0.950/0.830/0.951) and on a shifted map. **D**, Balanced accuracy of threshold rules, including the label-free per-slide two-component mixture threshold (within 0.01 of the oracle). On faded 10x post-Xenium H&E the detector failed (AUROC 0.599 and 0.705), so external sections used an RNA mask.

**Supplementary Fig. 4:**
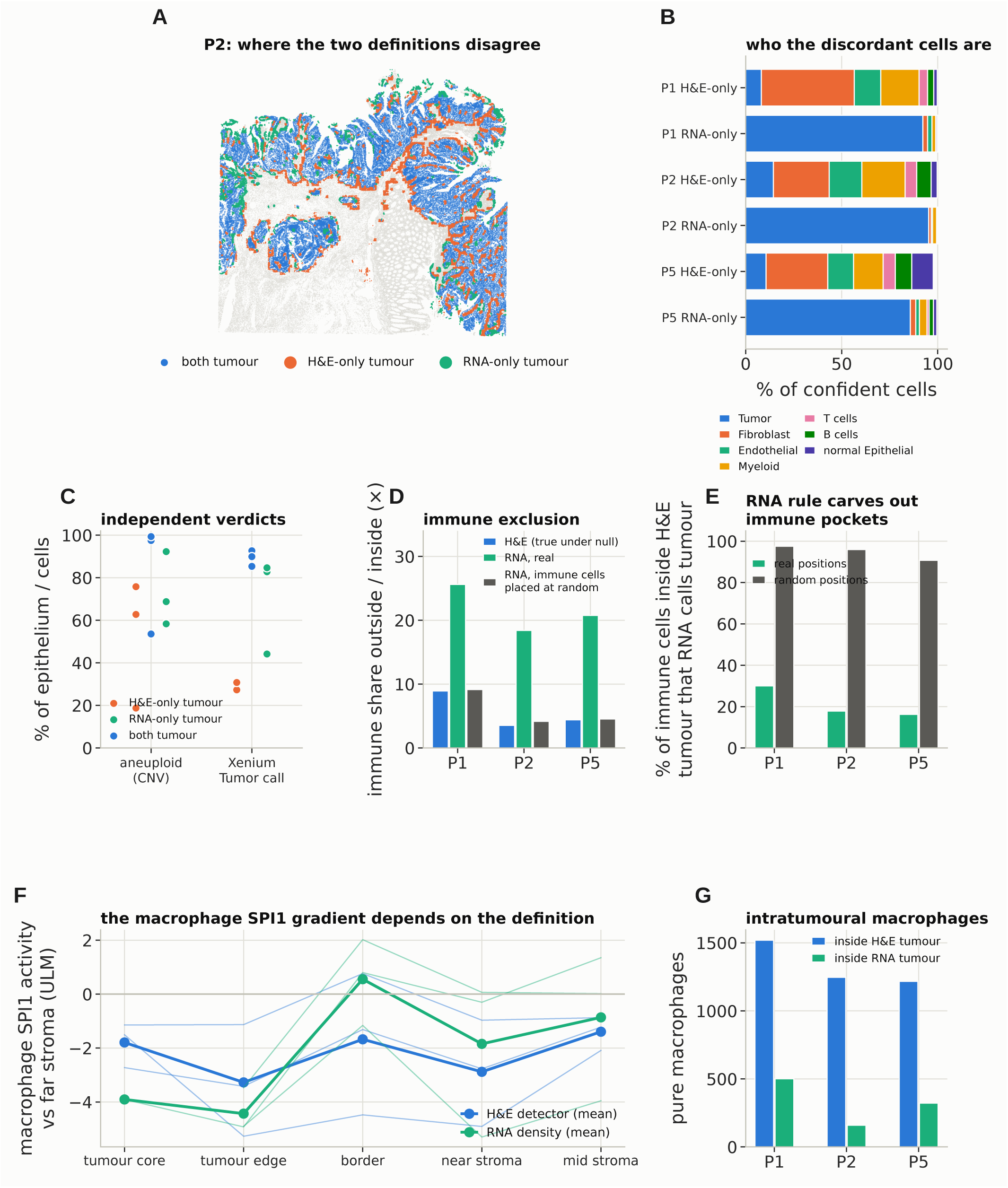
Region-definition bias: H&E detector versus RNA-density rule on the same cells. **A**, **B**, Where and in whom the two definitions disagree (P2 map; composition of discordant cells): H&E-only tumour is intratumoural stroma (fibroblasts, vessels, myeloid cells) that the RNA rule removes; RNA-only tumour is mostly genuine tumour cells beyond the morphological edge. **C**, Aneuploid share of epithelial metacells and RCTD-tumour share of registered Xenium cells per class: RNA wins on tumour-cell identity, the detector on the compartment. **D**, Immune exclusion under both definitions and after placing intratumoural immune cells at random tumour positions (bars as drawn, 8.9/3.5/4.4; with the final detector map reported in the text, 9.0/3.6/5.1; null 0.7–1.7; RNA rule 25.6/18.4/20.8). **E**, Share of H&E-intratumoural immune cells kept inside by the RNA rule: 16–30% at real positions versus 91–98% at random positions. **F**, Macrophage SPI1 activity by zone under both definitions; this gradient later failed the block null and was withdrawn. **G**, Purity-filtered macrophages inside each tumour definition (1,520/1,247/1,217 versus 501/159/322).

**Supplementary Fig. 5:**
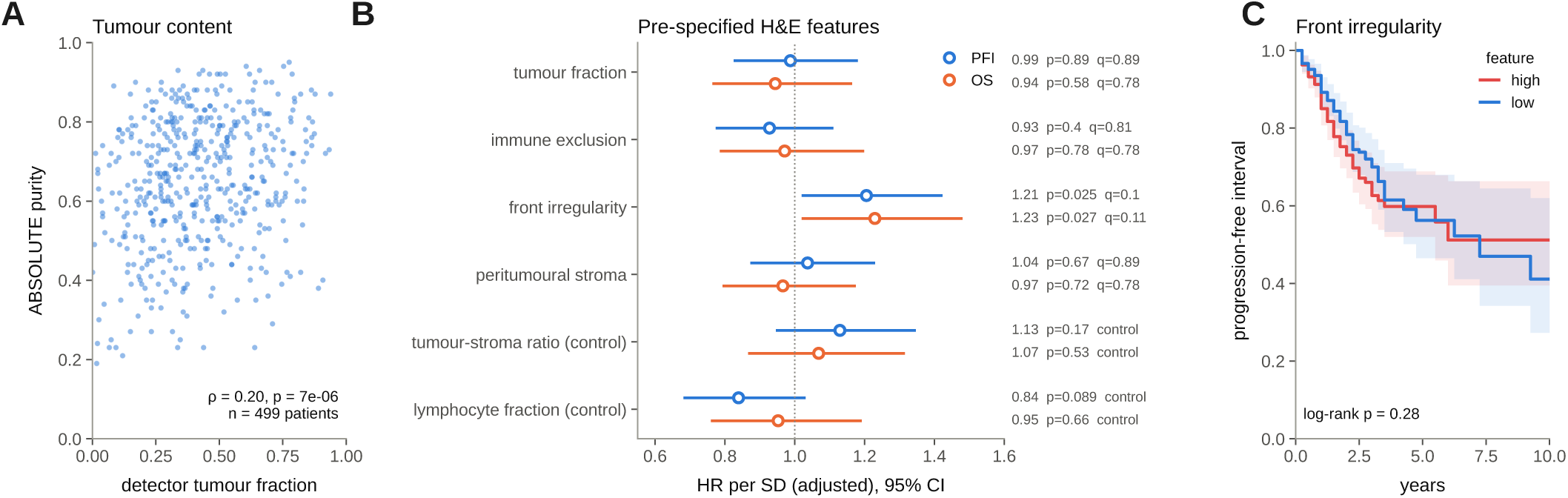
OmiCoreTumorDetector on 599 TCGA diagnostic slides: H&E map features are not prognostic. **A**, Detected tumour fraction versus ABSOLUTE purity (499 patients; Spearman *ρ* 0.20): a compartment map, not a purity estimator. **B**, Adjusted hazard ratios per s.d. for the four pre-specified H&E features (immune-exclusion index PFI 0.93 [0.78–1.11], OS 0.97 [0.79–1.19]; front irregularity PFI 1.21, *q* 0.10) and two positive controls. **C**, Progression-free interval by front irregularity (median split).

**Supplementary Fig. 6:**
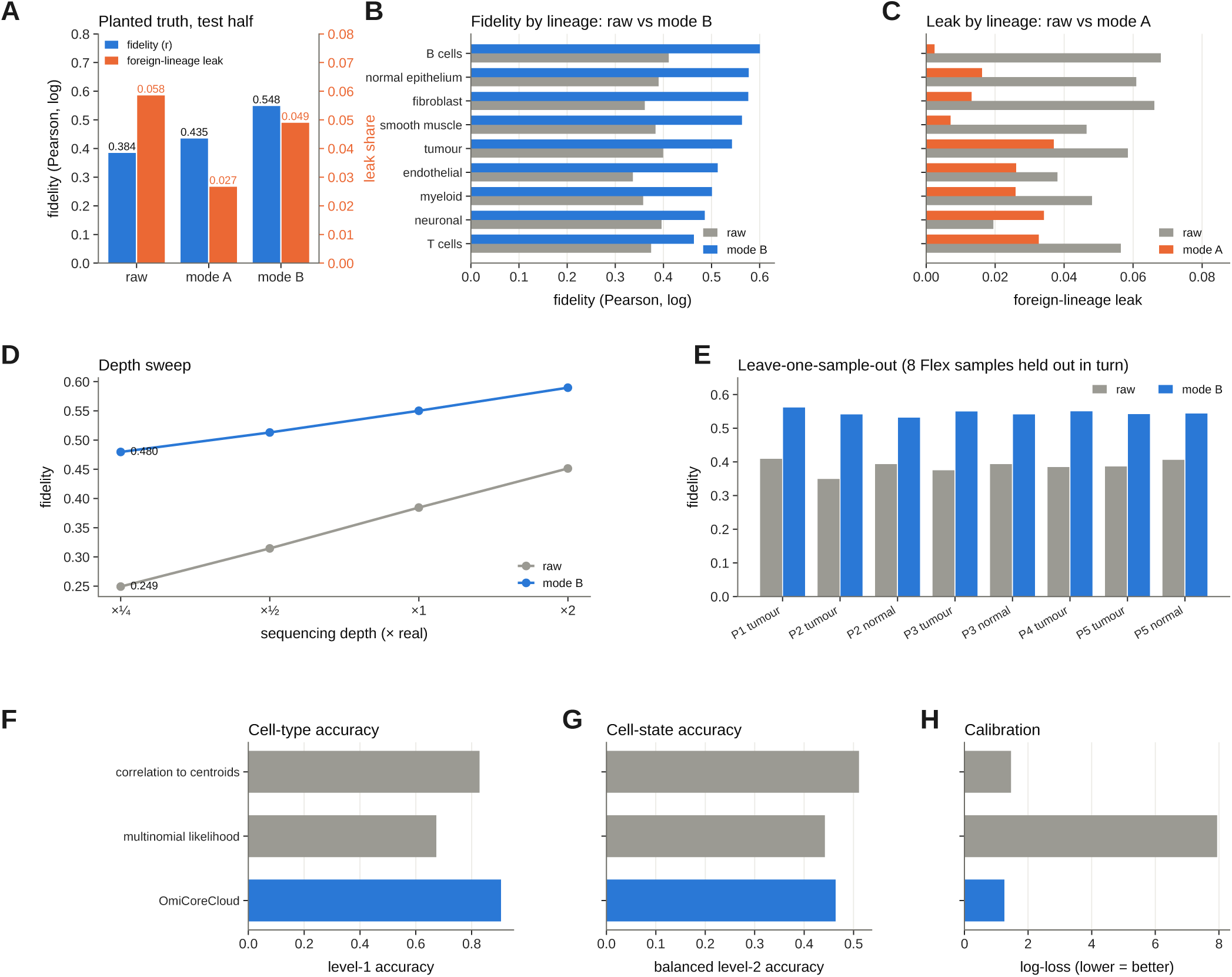
OmiCoreCloud benchmark against planted ground truth. Inputs: cell counts, positions, reference atlas and ambient prior; outputs: decontaminated (mode A) and denoised (mode B) profiles, state probabilities, their entropy and an atlas-misfit score (enthalpy). Planted truth: Flex cells of a held-out patient placed on real P1 geometry at real depth and contamination; test half only. **A**, Fidelity (Pearson, log) and foreign-lineage leak: raw 0.384, mode A 0.435, mode B 0.548; leak 0.058, 0.027 and 0.049. **B**, Fidelity by lineage: mode B above raw in 9/9 lineages. **C**, Foreign-lineage leak by lineage: mode A below raw in 8/9 (exception: neuronal). **D**, Depth sweep: at *×*0.25 depth, mode B 0.480 versus 0.249 raw. **E**, Leave-one-sample-out over the 8 Flex samples: 0.53–0.56 versus 0.35–0.41. **F**–**H**, Limits: cell-type accuracy, cell-state accuracy and calibration against standard callers; OmiCoreCloud is not a better cell-state caller (balanced level-2 accuracy 0.46 versus 0.51 for correlation to centroids). Mode B adds leak where little exists, and the planted-novelty AUROC of enthalpy is about 0.59, so atlas-misfit readouts are supportive only.

**Supplementary Fig. 7:**
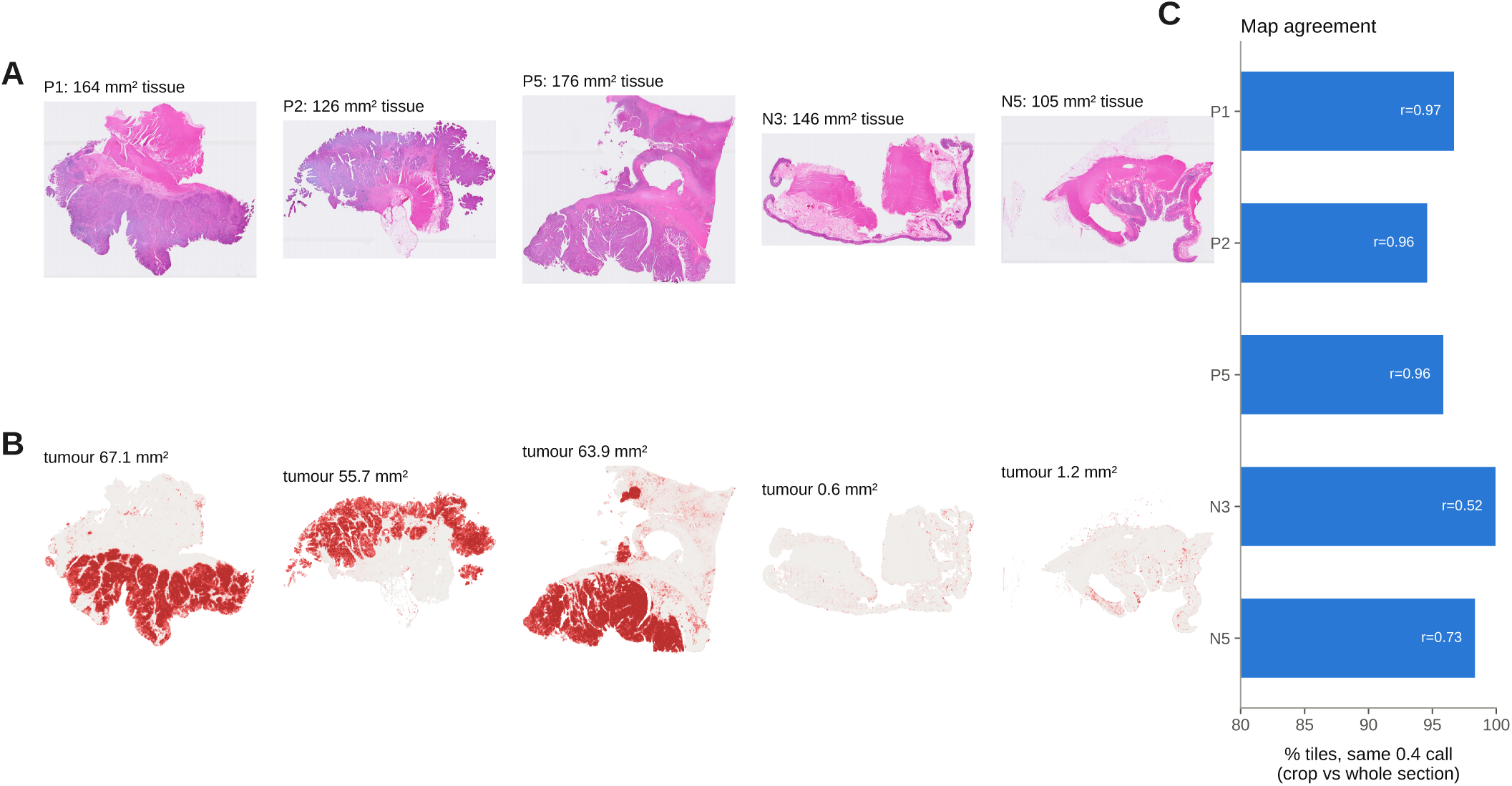
Whole-section OmiCoreTumorDetector maps. **A**, H&E overviews of the five discovery sections. **B**, P(tumour) over each whole section. **C**, Agreement of the capture-area crop map and the whole-section map (overlap *r* 0.957–0.968; call agreement 94.6–96.7%).

**Supplementary Fig. 8:**
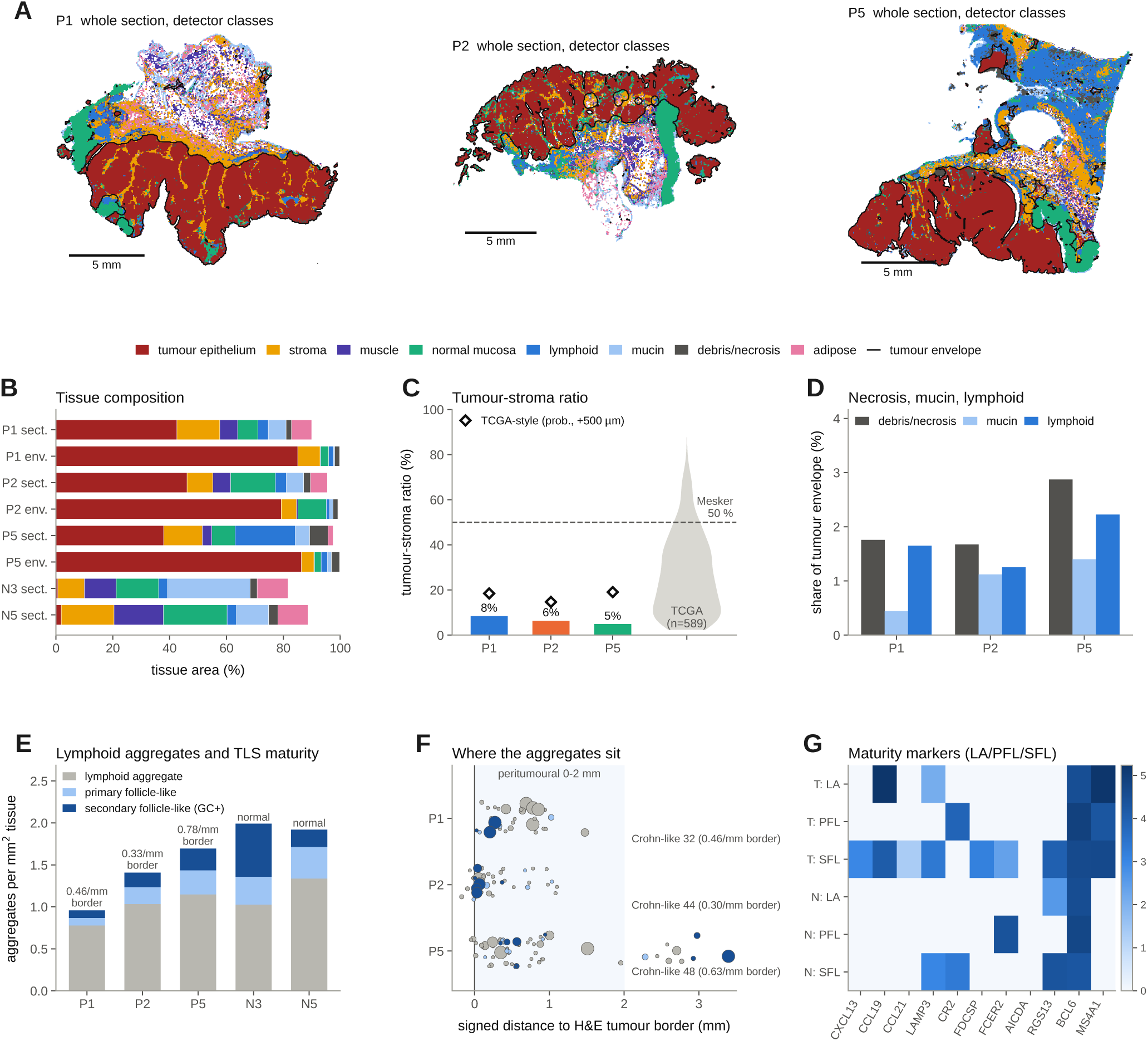
Histopathology and tertiary lymphoid structures (descriptive). **A**, Detector nine-class map of each whole section with the tumour envelope (*P ≥* 0.4 closed with a 250 µm disc). Scale bars, 5 mm. **B**, Tissue composition of each whole section and inside the envelope; normal sections N3 and N5 for reference. **C**, Tumour–stroma ratio inside the envelope (4.9–8.4%) with the Mesker 50% cut-off; diamonds, TCGA-style ratio (14.7–19.0%); violin, 589 TCGA patients. **D**, Debris/necrosis (1.7–2.9%), mucin (0.4–1.4%) and lymphoid area as shares of the envelope. **E**, Lymphoid aggregates per mm^2^ by maturity (lymphoid aggregate; primary follicle-like; secondary follicle-like, 9–15%); numbers, aggregates per mm of border. **F**, Signed distance of each aggregate to the border; shaded, the 0–2 mm Crohn-like zone (32/44/48 aggregates). **G**, TLS and maturity genes by class in tumour (T) and normal (N) sections. No test was performed.

**Supplementary Fig. 9:**
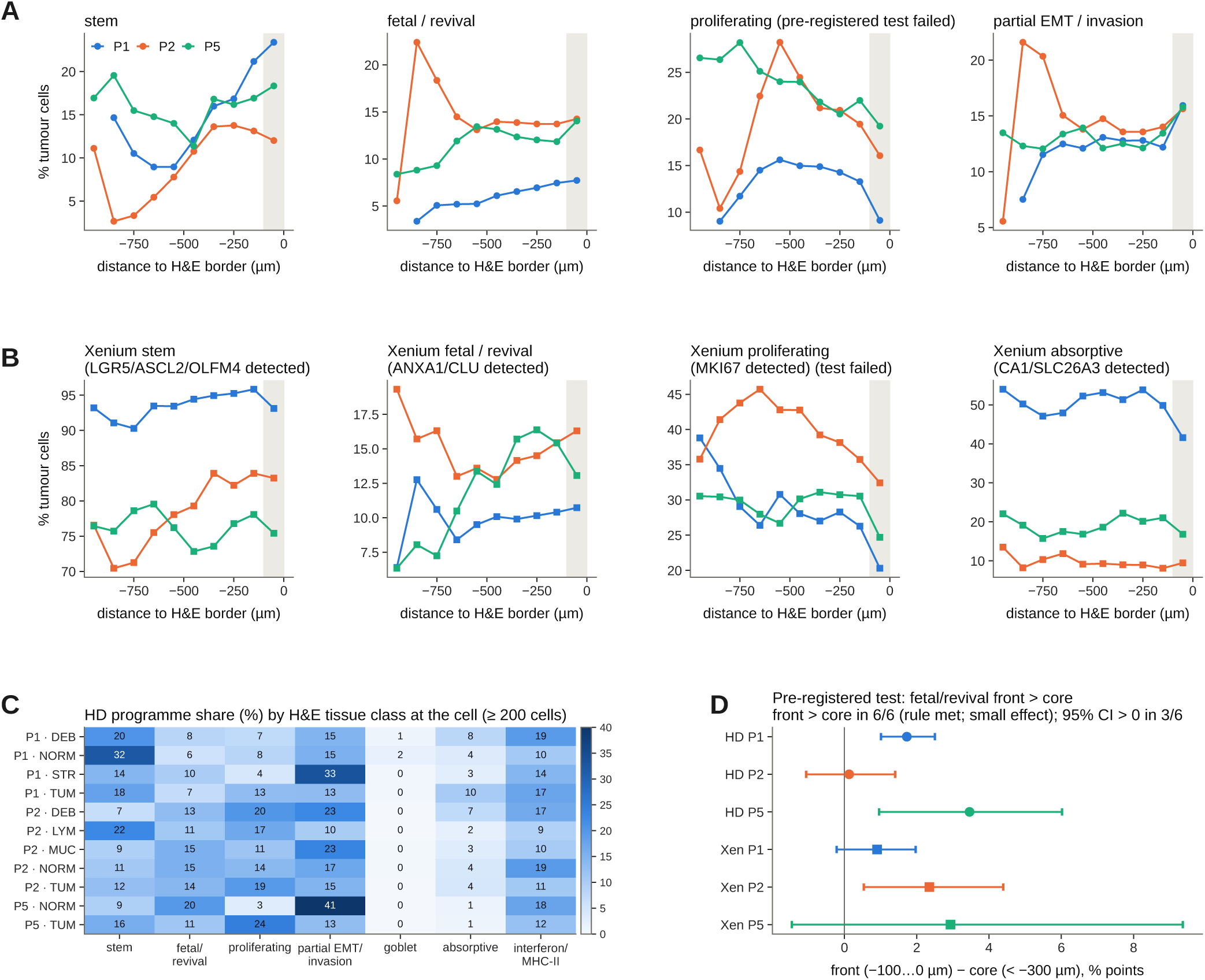
Tumour-cell state zonation across the H&E border. **A**, Share of confident tumour cells in each epithelial programme on Visium HD by 100 µm bin of the signed distance; grey band, front (*−*100 to 0 µm). Proliferation is descriptive only. **B**, The same on Xenium from panel markers (stem LGR5/ASCL2/OLFM4; fetal/revival ANXA1/CLU; proliferating MKI67; absorptive CA1/SLC26A3). **C**, Programme share by detector tissue class. **D**, Pre-registered test: fetal/revival share, front minus core (*< −*300 µm), percentage points, with 400 µm block-bootstrap 95% intervals: HD +1.7/+0.1/+3.5, Xenium +0.9/+2.4/+2.9; rule met 6/6, interval excludes 0 in 3/6. CLU is contamination-prone.

**Supplementary Fig. 10:**
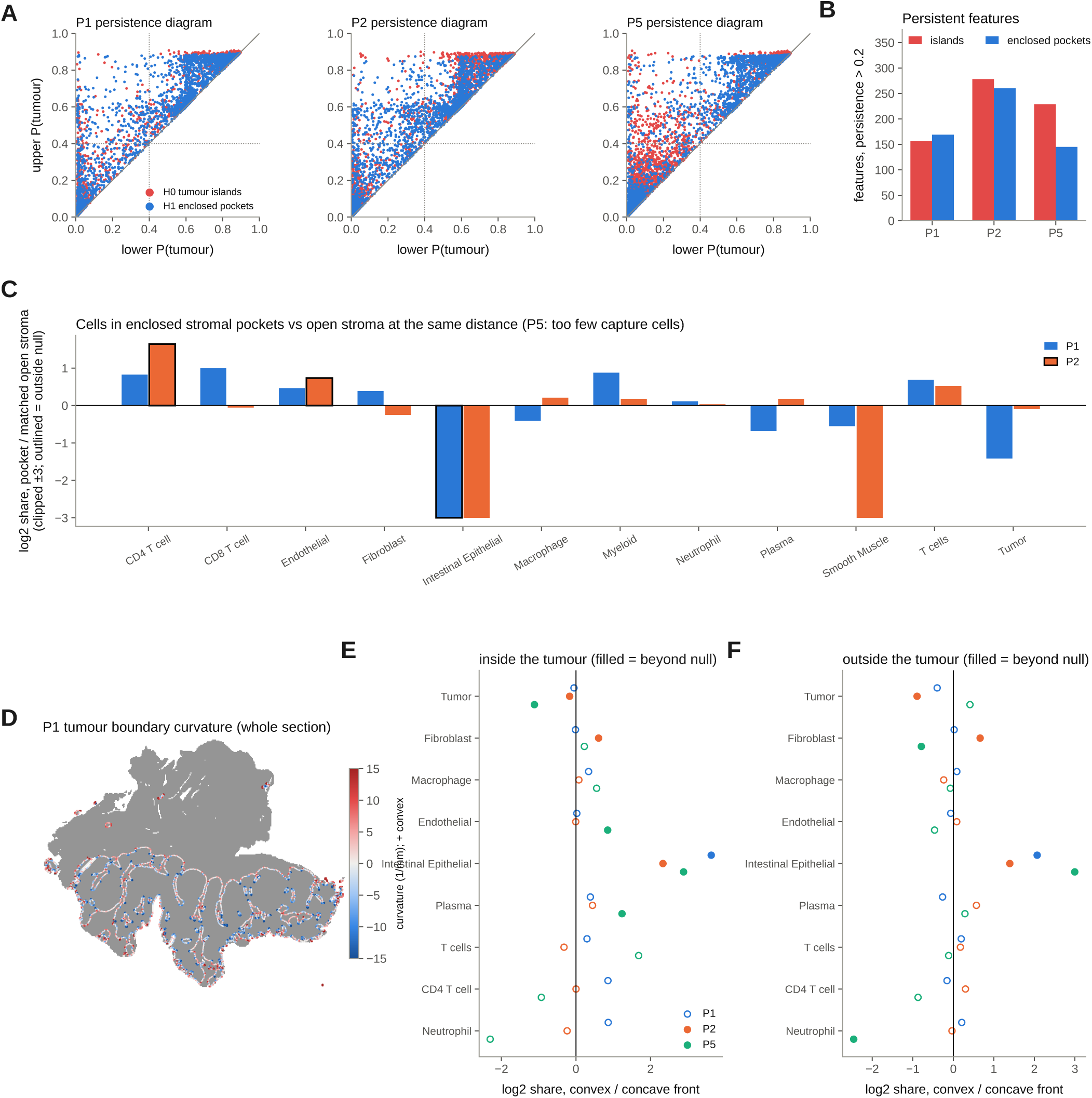
Topology and curvature of the tumour map. **A**–**C**, Exact cubical persistent homology of P(tumour). **A**, Persistence diagrams. **B**, Persistent tumour islands (157/278/229) and enclosed stromal pockets (169/260/145). **C**, CD4 T cells are enriched in pockets (log_2_ +0.82, +1.64; P2 beyond null); normal epithelium is absent from pockets. Descriptive. **D**–**F**, Level-set curvature of the H&E front (tortuosity 4.2/5.8/4.1). **D**, Curvature along the P1 tumour boundary. **E**, **F**, Normal epithelium at convex fronts (log_2_ +2.3 to +3.6 inside the tumour, **E**; +1.4 to +3.0 outside, **F**; 3/3 beyond a 200-map block null); validation SUPPORTED by direction only. Pre-registered front-shape *×* CAF/TAM and GREM1-at-convex tests failed.

**Supplementary Fig. 11:**
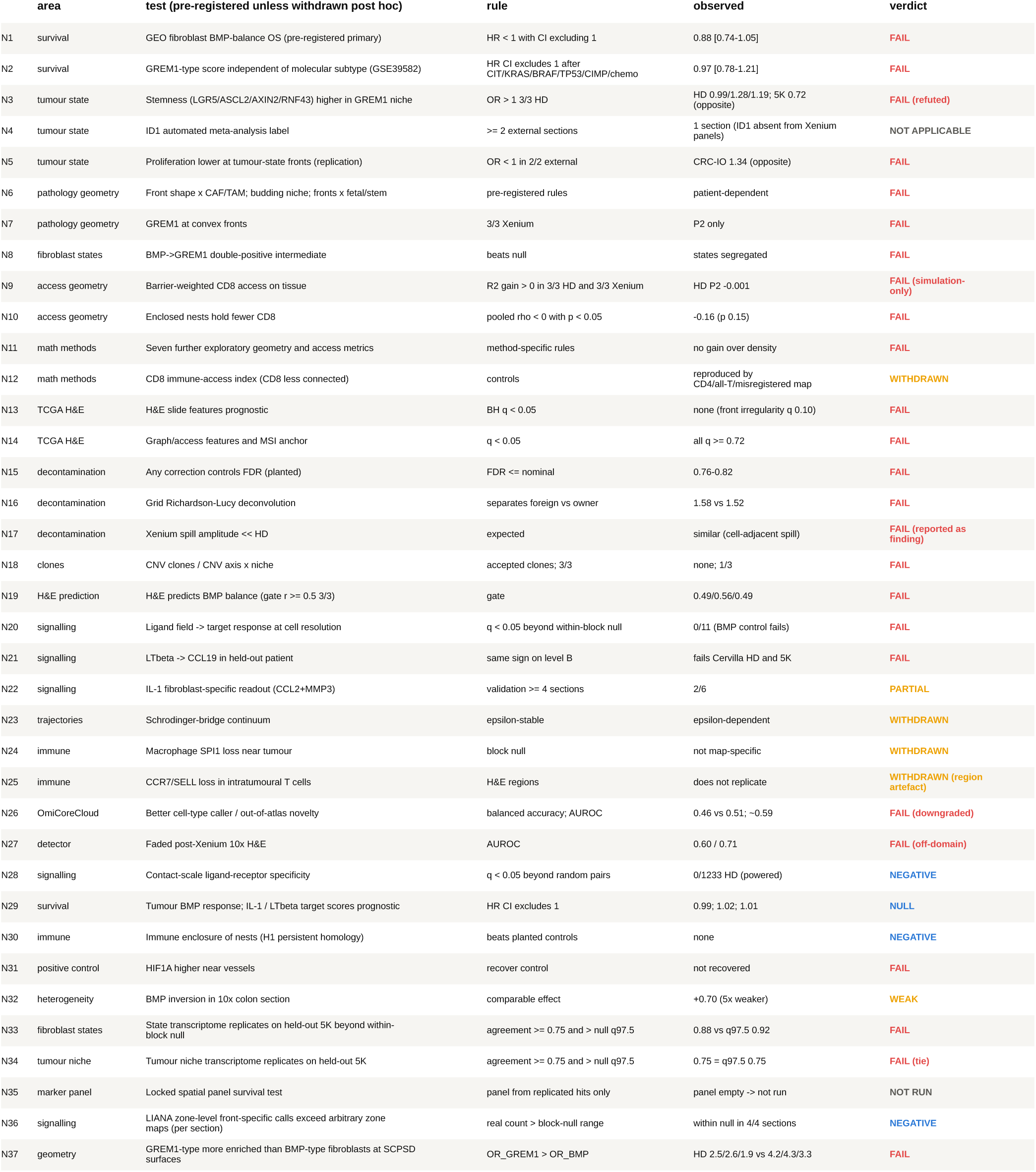
Pre-registered negative and failed results. Every test that failed, was withdrawn, was null or could not be run, with the rule fixed in advance, the observed value and the verdict. N33 and N34: gene-level replication of the fibroblast-state (0.88 versus null 97.5% 0.92) and tumour-niche (0.75, tie with the null 97.5%) transcriptomes on Xenium 5K. N35: the locked marker panel, empty and therefore not tested. N36: zone-level LIANA front-specific calls lie within the block-null range in every section. N37: the pre-stated GREM1-over-BMP enrichment at SCPSD surfaces fails.

**Supplementary Fig. 12:**
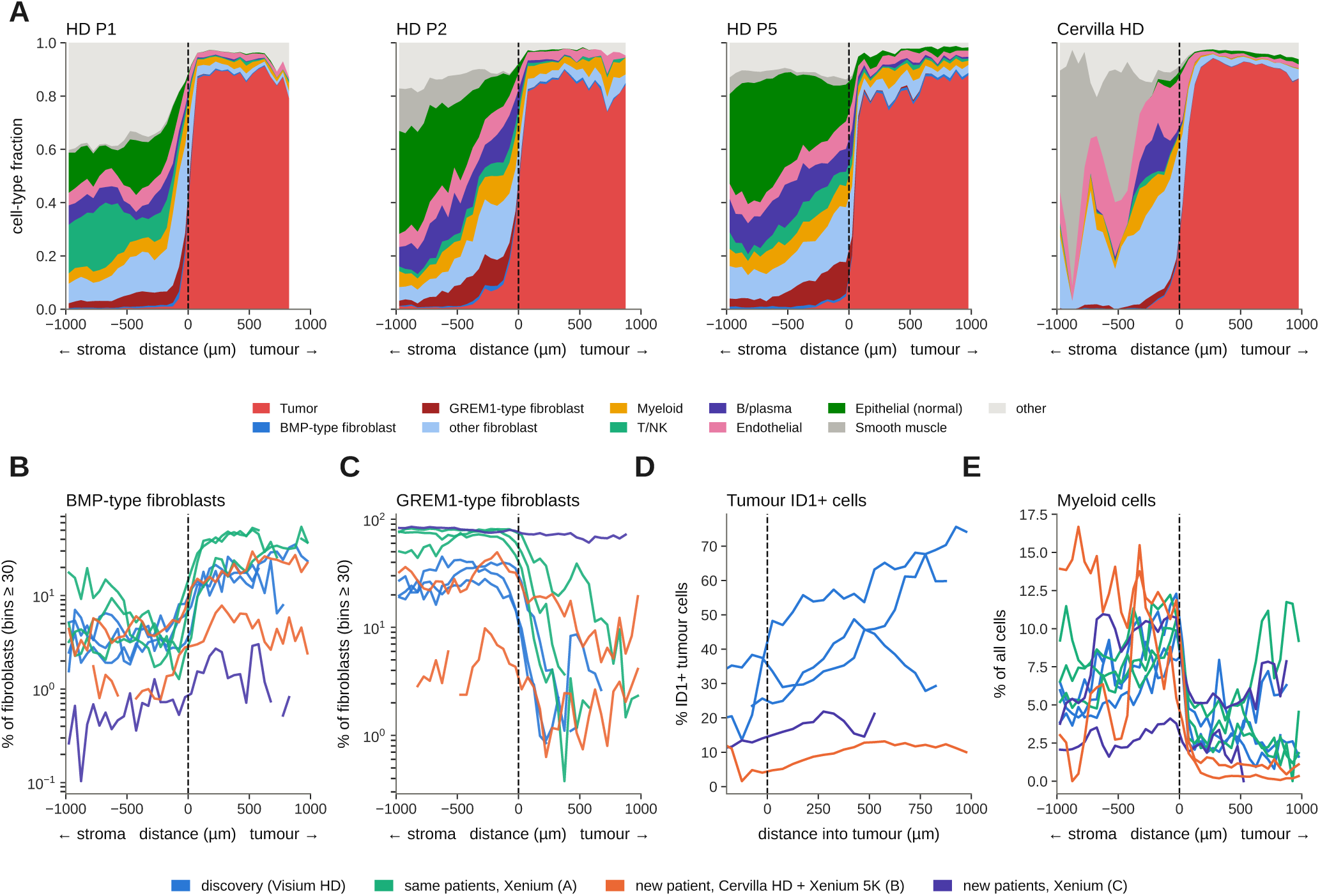
Profiles along the signed distance. (stroma *<* 0 *<* tumour as plotted; 50 µm bins). **A**, Stacked cell-type fractions (discovery and Cervilla). **B**, **C**, BMP-type and GREM1-type share of fibroblasts (bins with *≥*30 fibroblasts); in the 10x colon-cancer section GREM1-type fibroblasts dominate at every depth (79% stroma versus 71% tumour); the 10x CRC IO panel lacks BMP4/5 and GREM1. **D**, % ID1^+^ tumour cells with depth. **E**, The myeloid fraction peaks at the boundary (a fraction, not a subtype; no C1Q^+^ macrophage claim is made). In P2 and P5 the far side includes normal mucosa and smooth muscle; the anatomy-controlled contrast is in main Fig. 5F.

**Supplementary Fig. 13:**
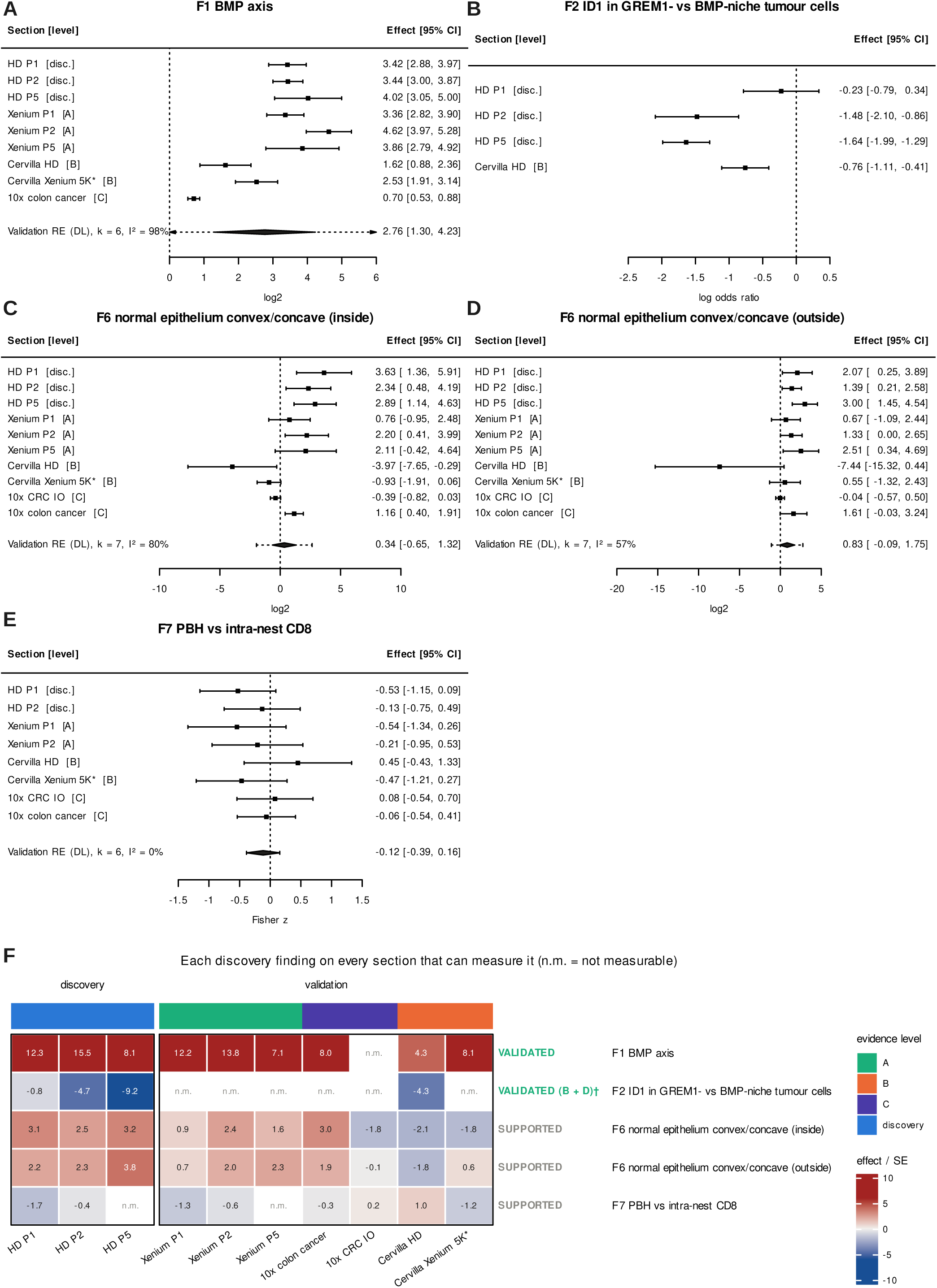
Meta-analysis and validation matrix. **A**–**E**, Random-effects meta-analysis per finding (metafor): each section’s effect (95% CI) by evidence level; diamond, DerSimonian–Laird pool of validation sections only with the 95% prediction interval (dashed). **A**, F1 (BMP axis): 6/6 positive, +2.76 [1.30, 4.23], *I*^2^ 98%. **B**, F2 (ID1): one validation section (Cervilla HD, log OR *−*0.76 [*−*1.11, *−*0.41]), so no pool. **C**, **D**, F6 (normal epithelium at convex fronts, inside and outside the tumour) and **E**, F7 (nest geometry versus intra-nest CD8): both SUPPORTED by direction only. **F**, Effect/s.e. per finding and section, split into discovery and validation columns with evidence level and verdict; n.m., not measurable. *†* F2 is validated by the held-out patient (B) plus population-level coupling in TCGA and GEO (D); the automated spatial-only rule could not be applied because ID1 is absent from the Xenium panels.

**Supplementary Fig. 14:**
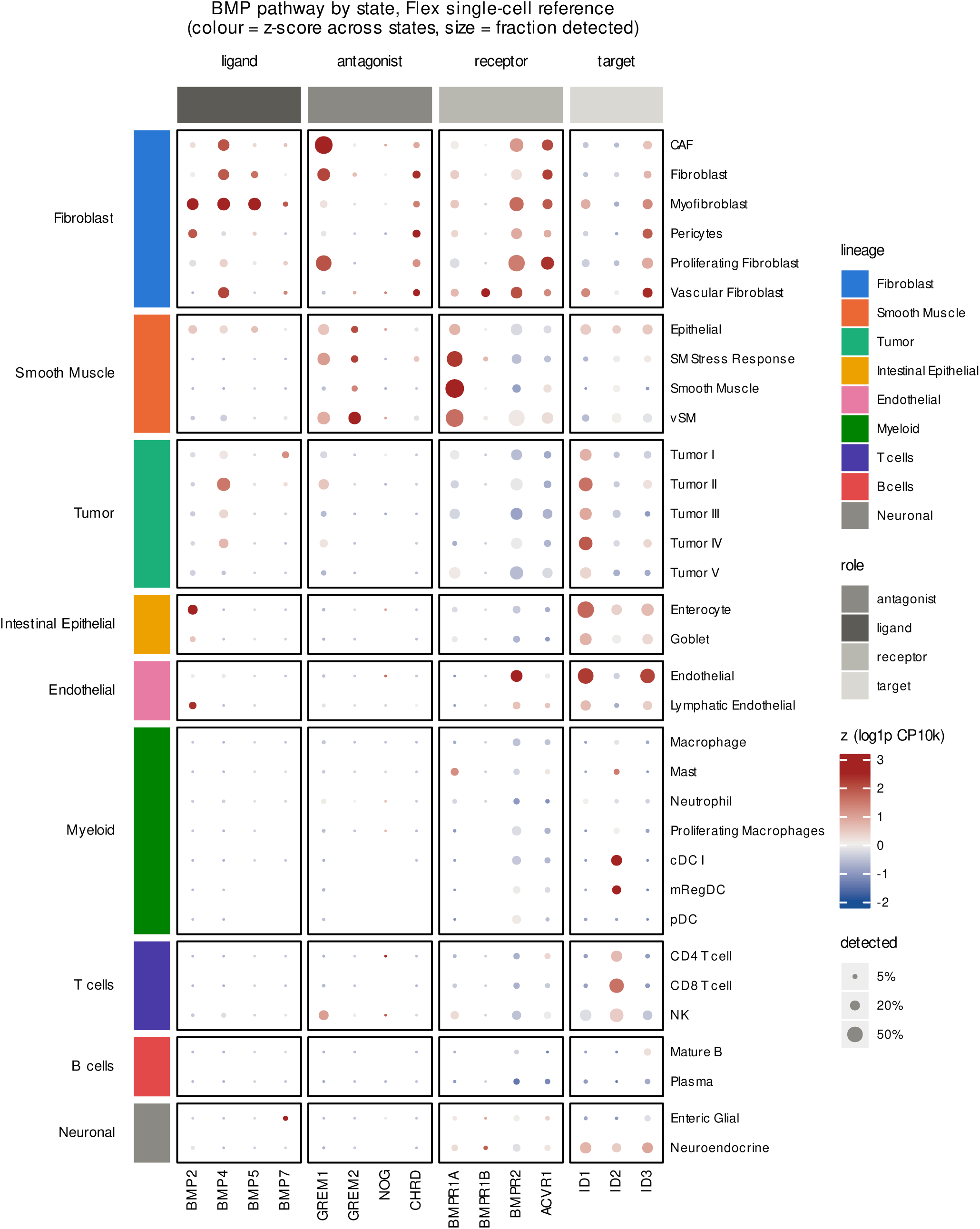
BMP pathway ownership in the Flex single-cell reference. Ligands, antagonists, receptors and targets by cell state; colour, z-score of log1p CP10k across states; dot size, fraction detected. GREM1 is CAF-owned (18.7 CP10k; smooth-muscle-like state 20.8), BMP4/5 fibroblast/myofibroblast-owned, and ID1 high in tumour (Tumor I–IV 2.57–6.39; Tumor V 1.65), epithelial and endothelial states; tumour cells express BMPR2, BMPR1A and ACVR1.

**Supplementary Fig. 15:**
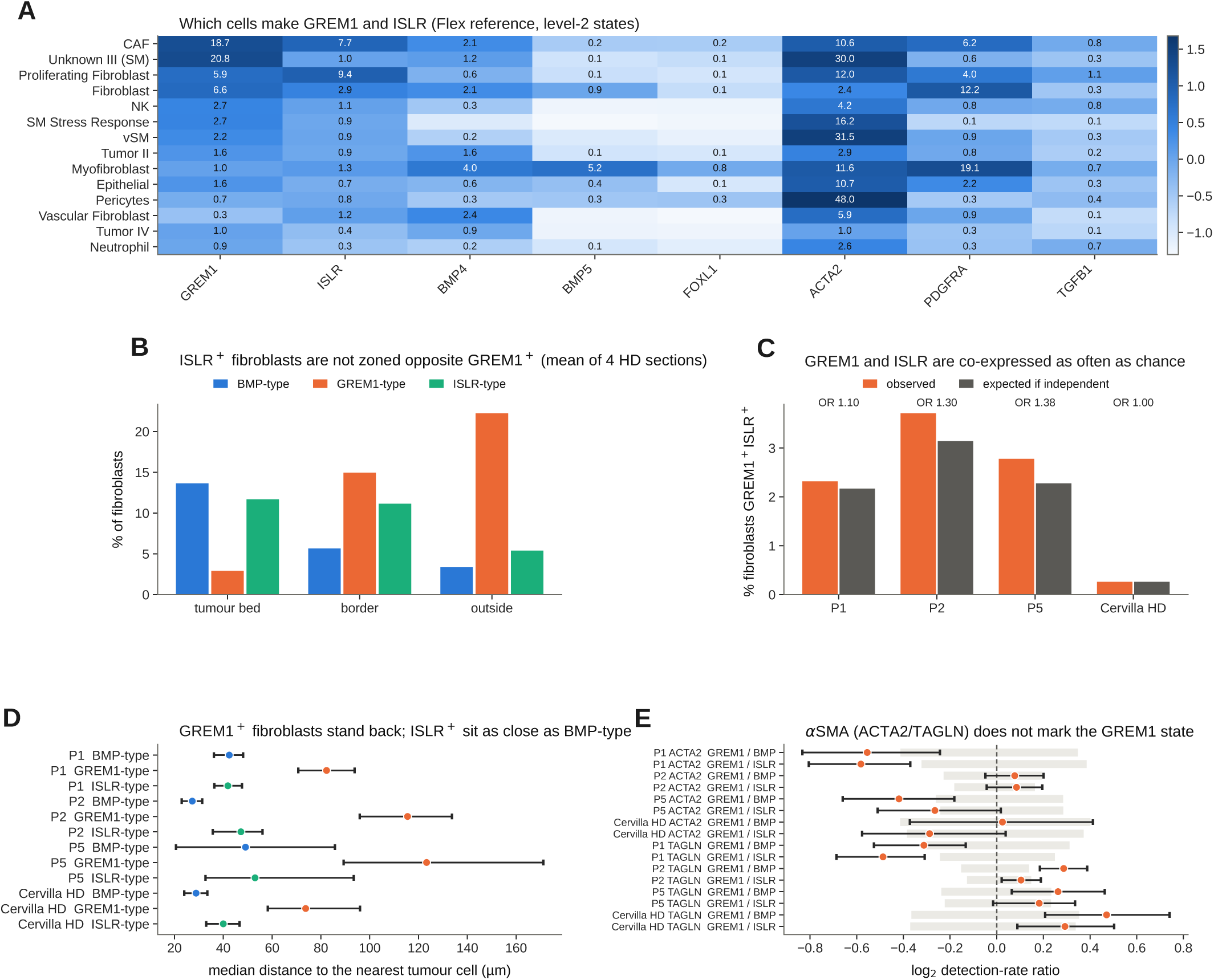
ISLR does not mark a fibroblast state opposite GREM1 in human colorectal cancer. **A**, Mean CP10k of the genes of the prior model across Flex level-2 states (14 states with *≥*100 cells, ranked by GREM1 + ISLR). **B**, Share of fibroblasts in each state by zone (tumour bed *d < −*50 µm; border *|d| ≤* 50 µm; outside), mean of the four Visium HD sections: ISLR-type fibroblasts follow the bed-enriched BMP-type pattern, not the front-enriched GREM1-type one. **C**, GREM1 and ISLR co-detection among fibroblasts, observed against the product of the marginals, with the co-detection odds ratio. **D**, Median distance from each fibroblast state to the nearest confident tumour cell, fibroblasts within 250 µm of the border; bars, 400 µm block-bootstrap 95% CI. **E**, *α*SMA markers by state, log_2_ detection-rate ratio with block-bootstrap CI; grey band, 400 µm block-permutation null. Source data are provided as a Source Data file.

**Supplementary Fig. 16:**
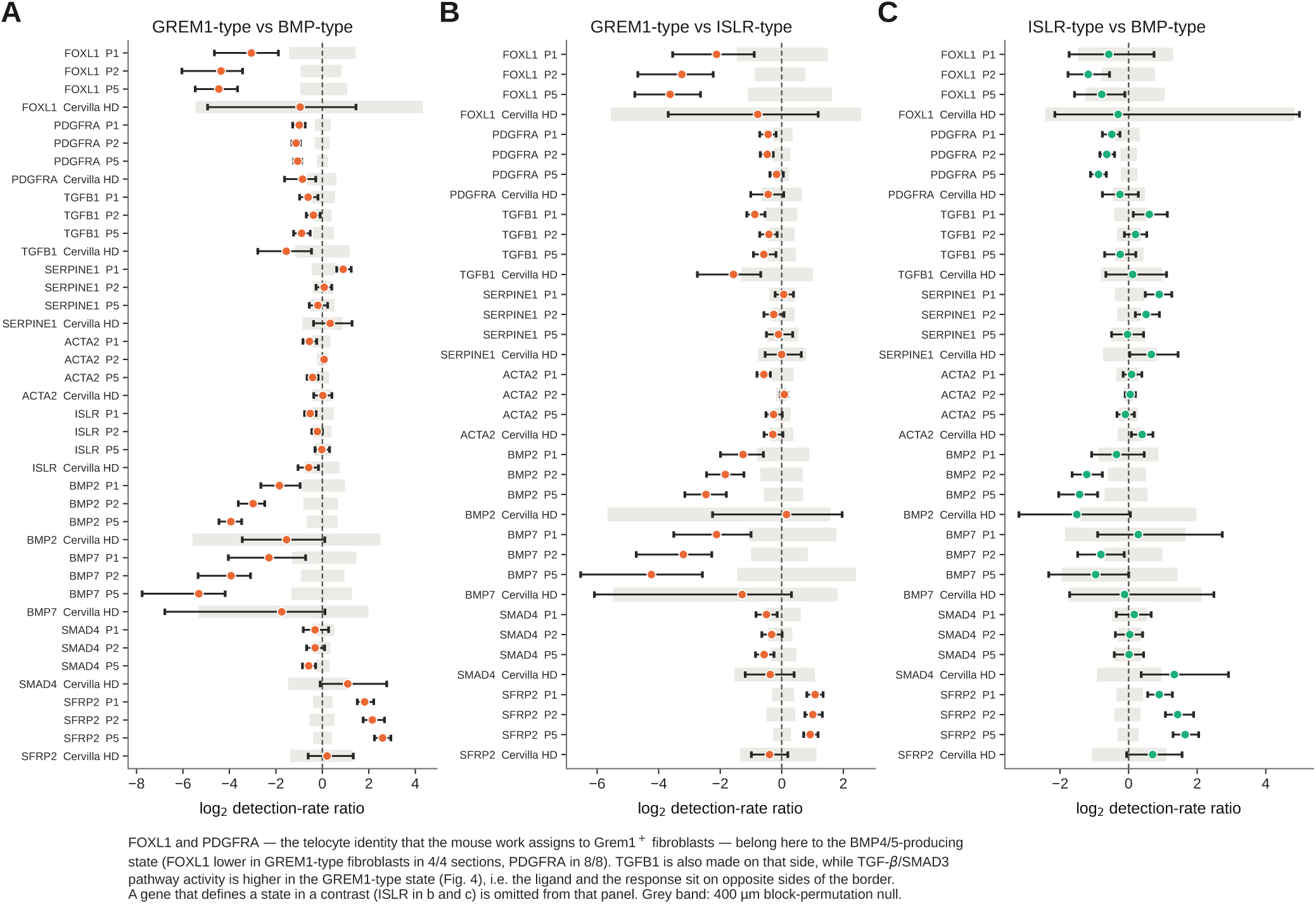
The TGF-*β*–FOXL1 arm of the prior model, measured in human tissue. Log_2_ detection-rate ratios between fibroblast states, per Visium HD section, with 400 µm block-bootstrap 95% CI and the 400 µm block-permutation null (grey band): **A**, GREM1-type versus BMP-type; **B**, GREM1-type versus ISLR-type; **C**, ISLR-type versus BMP-type. A gene that defines one of the two states in a contrast is omitted from that panel. FOXL1 and PDGFRA, the telocyte identity the mouse work assigns to Grem1^+^ fibroblasts, are higher in the BMP4/5-producing state here, as are BMP2, BMP7 and TGFB1, while SFRP2 is higher in the GREM1-type state.

**Supplementary Fig. 17:**
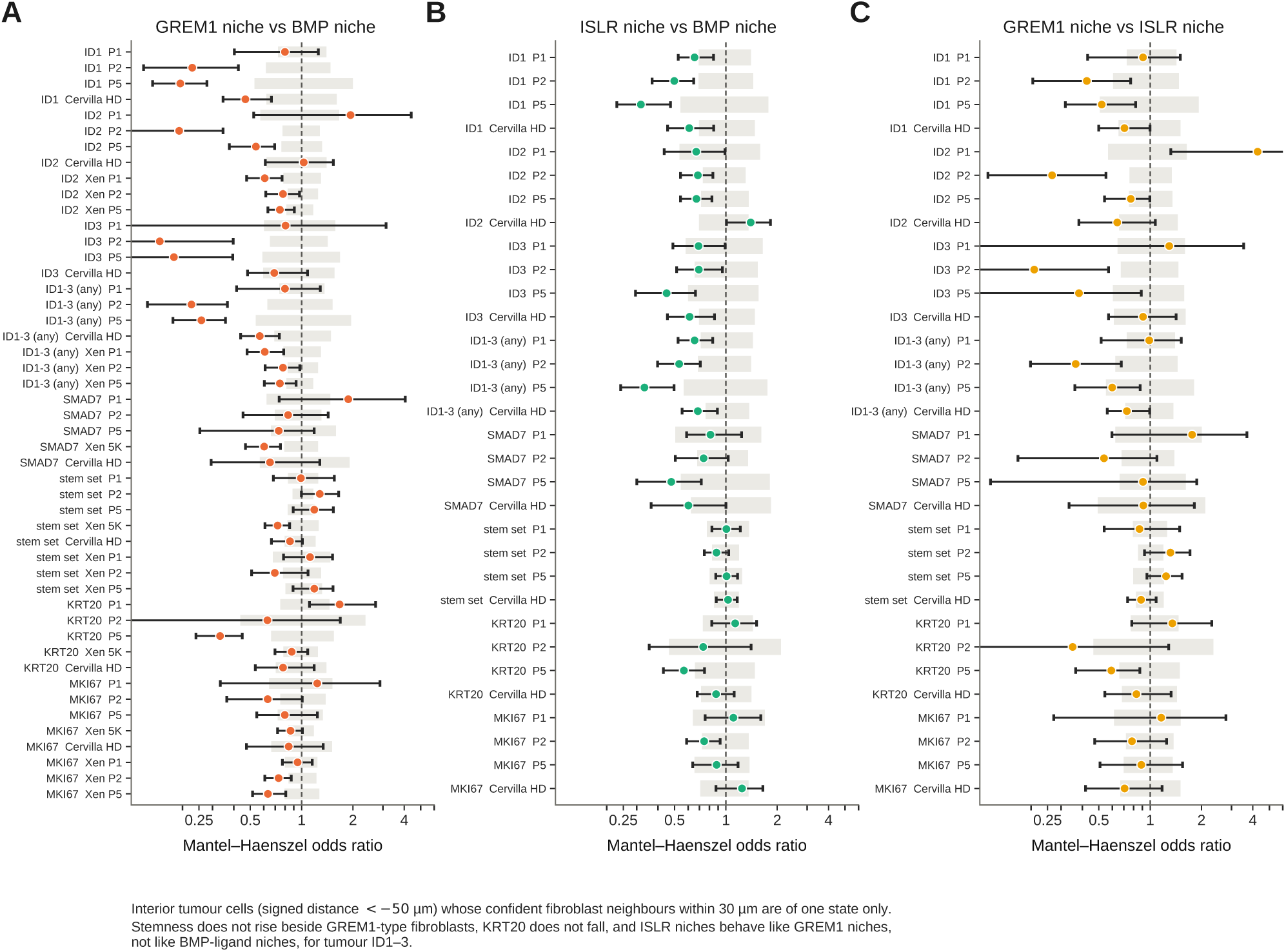
Tumour BMP output, stemness and differentiation by the state of the neighbouring fibroblasts. Mantel–Haenszel odds ratios for interior tumour cells (*d < −*50 µm) whose confident fibroblast neighbours within 30 µm are of one state only, stratified by 25 µm distance bin and neighbour count, with 400 µm block-bootstrap 95% CI and the block-permuted state null (grey band): **A**, GREM1-type versus BMP-type niche; **B**, ISLR-type versus BMP-type; **C**, GREM1-type versus ISLR-type. ID1–3 fall beside both GREM1-type and ISLR-type fibroblasts relative to BMP-type; the stem set does not rise and KRT20 does not consistently fall.

**Supplementary Fig. 18:**
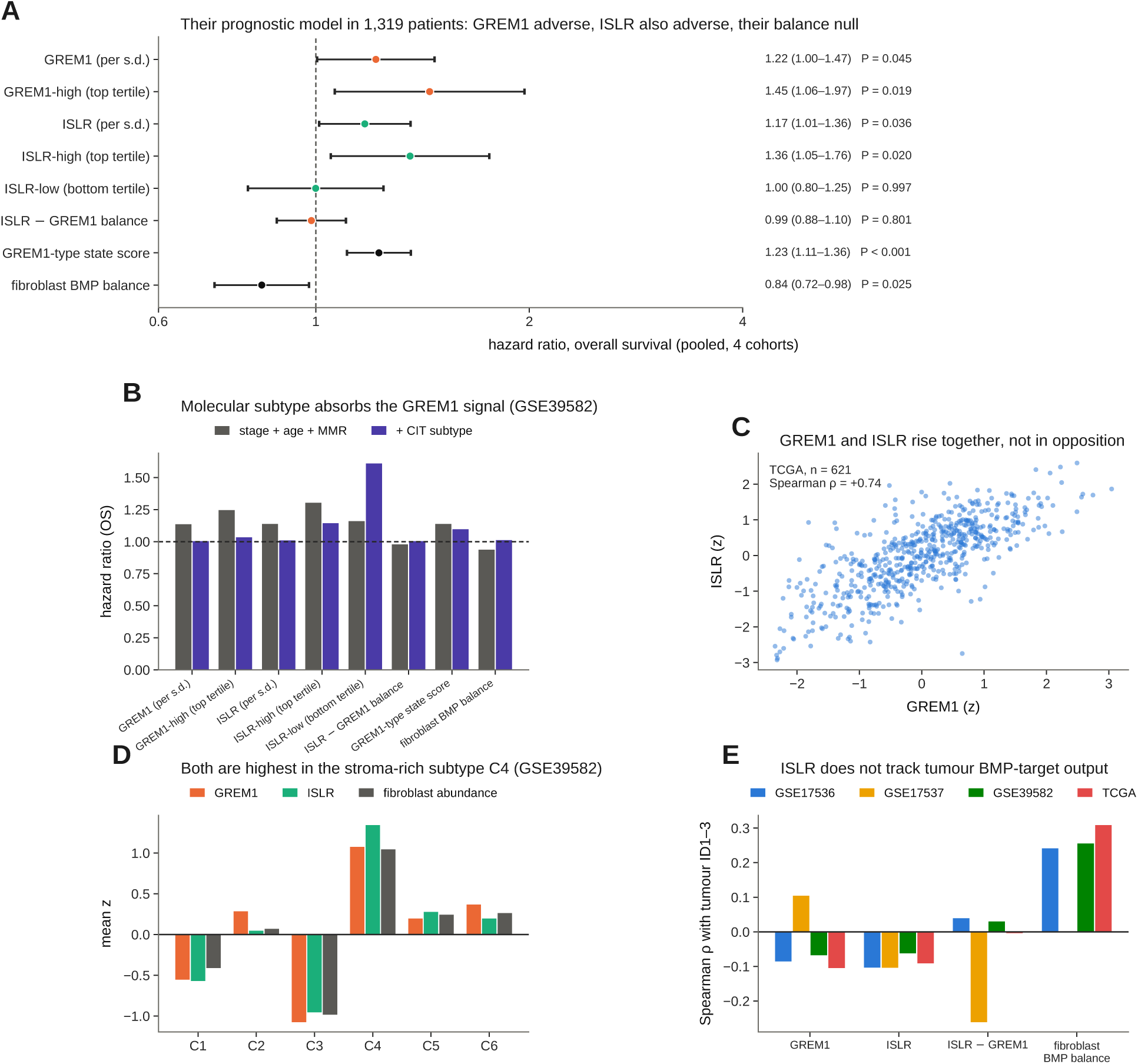
The GREM1/ISLR prognostic model in four bulk cohorts (1,319 patients). **A**, DerSimonian–Laird pooled hazard ratios for overall survival, adjusted for stage and age (+ MMR where available), for the prior model’s exposures and, for reference, the state scores of this study. **B**, GSE39582: the same hazard ratios before and after adding CIT molecular subtype. **C**, GREM1 against ISLR across TCGA tumours. **D**, Mean gene z and fibroblast abundance by CIT subtype. **E**, Spearman correlation of each score with residualised tumour ID1–3 expression, per cohort. Scores are gene z of log expression; tertile groups reproduce the GREM1-high and ISLR-low definitions of the prior work ^13^.

**Supplementary Fig. 19:**
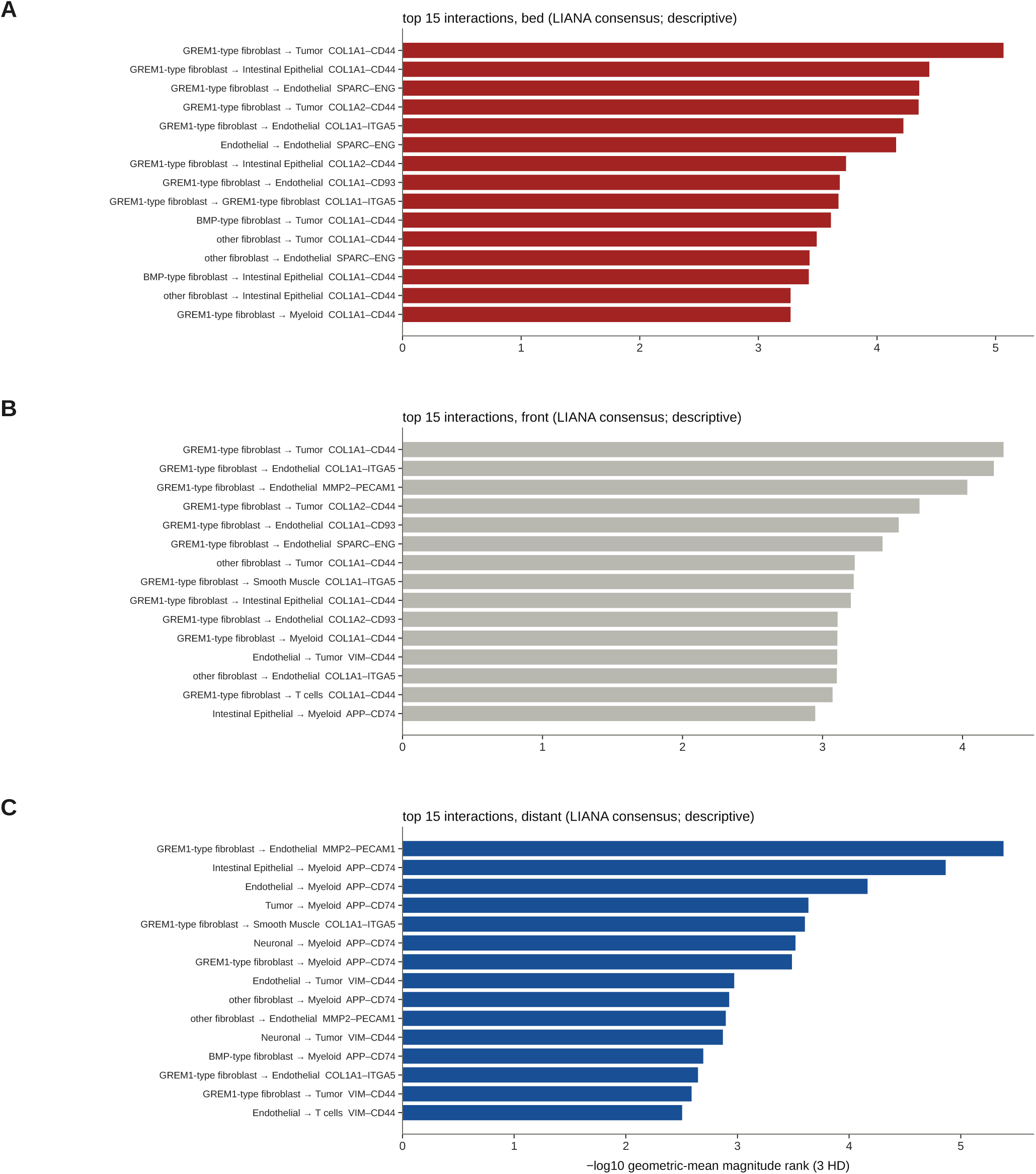
Zone-resolved LIANA results in full. Top 15 interactions in the tumour bed (**A**), at the front (**B**) and in distant stroma (**C**), ranked by the geometric mean over the three discovery sections of the LIANA consensus magnitude rank (fibroblasts split into BMP-type, GREM1-type and other). The same collagen–CD44/integrin pairs lead in every zone. Descriptive; the spatial null and the front-specific rule are in Fig. 7A–D. Full results are provided as Source Data.

**Supplementary Fig. 20:**
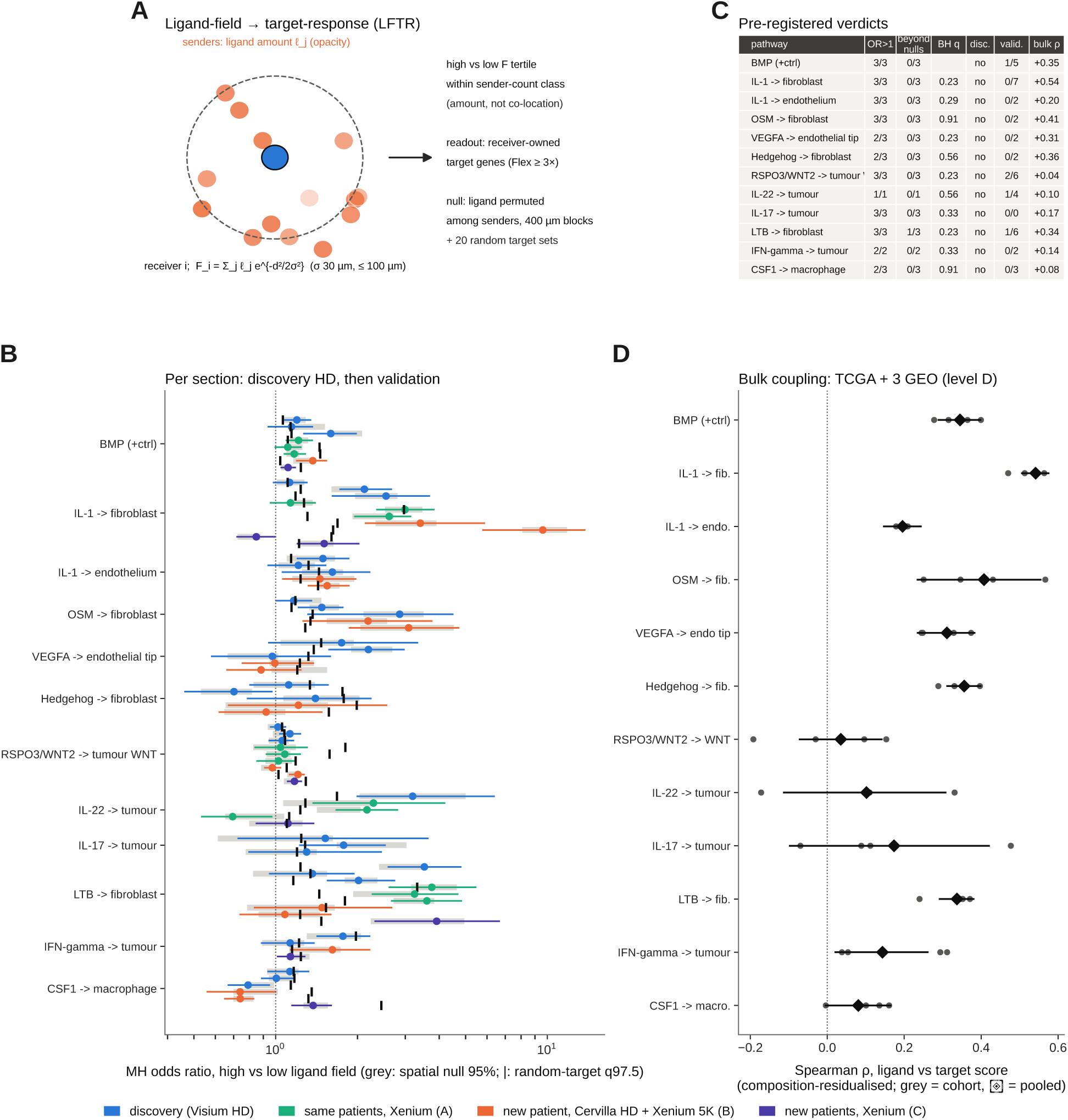
Pre-registered ligand-field*→*target-response (LFTR) screen at cell resolution. **A**, Design: each receiver’s paracrine field 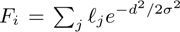 over senders within 100 µm (*σ* = 30 µm); high versus low tertile of *F* compared within sender-count classes; readout, detection of receiver-owned targets. **B**, Mantel–Haenszel odds ratio per pathway and section with block-bootstrap 95% CI; grey, null from ligand permuted among senders within 400 µm blocks; *|*, 97.5% quantile of 20 detection-matched random target sets. **C**, Verdicts: no pathway, including the BMP positive control, passes at cell resolution (calibration 2.0% versus 2.5% expected). **D**, Bulk coupling of composition-residualised ligand and target scores in TCGA and three GEO cohorts (7/11 test pathways coupled). The contact-scale co-expression test (0/1,233 HD pairs; *≤*1/158 Xenium pairs; powered to detect a 9-point juxtacrine rise in 68% of slides) is described in the main text.

**Supplementary Table 1.** OmiCoreSeg and open-source comparators on PanNuke colon nuclei. 962 patches, 23,550 nuclei; IoU *>* 0.5 matching; 95% CI from 2,000 patch bootstraps. Comparators were run on unmodified H&E with their published models (benchmark comparators only).

| Tool | Correct | False | Missed | Precision | Recall | F1 | SQ | PQ | PQ 95 % CI | PQ 00–07 / 08–15 |
|---|---|---|---|---|---|---|---|---|---|---|
| **OmiCoreSeg** | 14,588 | 4,675 | 8,962 | 0.757 | 0.619 | 0.681 | 0.790 | **0.538** | 0.526–0.550 | 0.539 / 0.537 |
| InstanSeg | 12,253 | 2,804 | 11,297 | 0.814 | 0.520 | 0.635 | 0.791 | 0.502 | 0.489–0.515 | 0.496 / 0.510 |
| Cellpose-SAM | 12,531 | 2,896 | 11,019 | 0.812 | 0.532 | 0.643 | 0.752 | 0.484 | 0.473–0.494 | 0.480 / 0.488 |
| StarDist (tuned) | 13,662 | 6,856 | 9,888 | 0.666 | 0.580 | 0.620 | 0.755 | 0.468 | 0.456–0.480 | 0.469 / 0.468 |
| QuPath | 10,059 | 11,256 | 13,491 | 0.472 | 0.427 | 0.448 | 0.691 | 0.310 | 0.300–0.320 | 0.306 / 0.315 |
| scikit-image | 7,755 | 11,042 | 15,795 | 0.413 | 0.329 | 0.366 | 0.726 | 0.266 | 0.253–0.280 | 0.247 / 0.287 |
| StarDist (default) | 4,956 | 1,308 | 18,594 | 0.791 | 0.210 | 0.333 | 0.793 | 0.264 | 0.246–0.281 | 0.253 / 0.276 |
| OpenCV | 5,595 | 22,017 | 17,955 | 0.203 | 0.238 | 0.219 | 0.668 | 0.146 | 0.139–0.153 | 0.140 / 0.153 |
| CellProfiler | 5,046 | 78,539 | 18,504 | 0.060 | 0.214 | 0.094 | 0.629 | 0.059 | 0.053–0.066 | 0.061 / 0.058 |

**Supplementary Table 2.** Pre-registered tests and outcomes. Every test was written into the analysis record before it was run and was run once. PASS/FAIL refers to the pre-set rule.

| Test | Rule | Result | Outcome |
|---|---|---|---|
| Spill kernel | σ<sub>1</sub> within 8–16 µm; 1 mm shift removes gain | σ<sub>1</sub> 8–12 µm all sections; 82–100% removed | PASS |
| Xenium leak ≪ HD | expected | similar (cell-adjacent spill) | FAIL (reported as finding) |
| Grid Richardson–Lucy | separates foreign from owner | 1.58 vs 1.52 | FAIL; replaced by cell inversion |
| Headlines after deconvolution | rules fixed in advance | all survive; IFN/MHC-II stays closed | PASS |
| External BMP axis | bed − front > null 97.5% | Cervilla HD +1.62 (0.43); 10x colon +0.70 (0.16); 5K +2.53 (0.62) | PASS |
| F1 meta-analysis | validation-only CI excludes 0, ≥2 levels | +2.76 [1.30, 4.23], 6/6, *I*<sup>2</sup> 0.98 | VALIDATED |
| SCPSD at intrusions (F4) | OR above both controls; sign test | 7/7 validation (*P* 0.0078); 10/10 | VALIDATED |
| Normal epithelium at convex fronts (F6) | validation pool CI excludes 0 | inside +0.34 [−0.65, 1.32]; outside +0.83 [−0.09, 1.75] | SUPPORTED |
| CAF zonation, Cervilla | ≥70% sign agreement and > null 97.5% | 21/22 (0.955; null 0.947) | PASS |
| CAF zonation, Xenium 5K | same | 11/12 (0.917; null 0.877) | PASS |
| BMP switch without muscle/mucosa tiles | > 0 3/3 HD, > null in ≥2/3; Xenium 3/3 | HD +3.59/+3.48/+4.37; Xenium +3.48/+4.73/+4.26 | REPLICATED |
| State-transition direction (BMP- to GREM1-type intermediate) | directional transitions beyond null | segregated; no intermediate | FAIL |
| Stemness in GREM1 niche (primary) | OR > 1 3/3 HD | 0.99/1.28/1.19; 5K 0.72 [0.60–0.89] | FAIL (refuted) |
| ID1 confirmation, Cervilla HD | OR < 1, upper CI < 1, below null 2.5% | 0.47 [0.33–0.67]; null 0.65 | CONFIRMED |
| F2 automated meta rule | ≥2 external sections | 1 section | not applicable; VALIDATED (B + D) |
| TCGA / GEO coupling | ρ > 0 beyond random sets / pooled CI > 0 | +0.31 (>0.15); GEO +0.22 [0.12–0.32] | PASS |
| CD8 IM > CT | 3/3 HD and 3/3 Xenium | HD 2.27/3.25/1.90; Xenium 2.06/1.40/1.64 | REPLICATED |
| Fetal/revival front > core | 6/6 | +0.1 to +3.5 points; CI > 0 in 3/6 | rule met (small) |
| Proliferation at state fronts | OR < 1 in 2/2 external | CRC IO 1.34 (1.02–1.71) | FAIL |
| Front shape, budding, GREM1 at convex fronts | pre-registered rules | patient-dependent | FAIL |
| Contact-scale LR | > random-pair null on ≥2/3 slides | 0/1,233 HD; ≤1/158 Xenium; powered | NEGATIVE |
| LFTR cell resolution | beyond within-block null | 0/11; BMP control fails | FAIL |
| LFTR niche scale | beyond shift null; replication | IL-1→fibroblast 4/7; LTβ →fibroblast 4/6 | REPLICATED (2 axes) |
| Readout sensitivity | as niche-scale LFTR, specific targets | CCL19 4/6 (fails level B); CCL2+MMP3 2/6 | robust / partial |
| GEO OS, BMP balance (primary) | HR < 1, CI excludes 1 | 0.88 [0.74–1.05] | **FAIL** |
| GEO OS, GREM1-type score (secondary) | HR > 1 | 1.18 [1.05–1.34], *I*<sup>2</sup> 0 | REPLICATES |
| GREM1 score, leave-one-out | every pooled CI excludes 1 | 1.19–1.32 | ROBUST |
| GREM1 score, molecular adjustment | HR > 1 | 0.97 [0.78–1.21] (*n* 280) | DOES NOT HOLD |
| Score × SMAD4 | – | no mutation calls | NOT TESTABLE |
| TCGA H&E features | BH *q* < 0.05 | none (front irregularity *q* 0.10) | FAIL |
| H&E predictor of BMP balance | held-out tile *r* ≥ 0.5 in ≥2/3 | 0.49/0.56/0.49 | FAIL (narrow) |
| Fibroblast-state transcriptome on 5K | agreement ≥0.75 and > within-block null 97.5% | 0.881 (135 genes); null 0.919 | NOT REPLICATED |
| Tumour-niche transcriptome on 5K | same | 0.75 (8 genes); null 0.75 | NOT REPLICATED (tie) |
| Locked marker panel survival | panel from replicated genes only | panel empty | NOT RUN |
| Zone-resolved LIANA | front-specific calls > null 95%; 5K replication | 7 vs null 0.1 (q95 0.55); per section within null; 0 testable on 5K | PASS (count) / NOT TESTABLE (5K) |
| SCPSD content | 3/3 HD, CI excludes 1 in ≥2/3; Xenium 3/3; OR<sub>GREM1</sub> > OR<sub>BMP</sub> | 7/7 neighbour types and fetal/revival, proliferation confirmed; BMP > GREM1 | CONFIRMED / hypothesis FAIL |

**Supplementary Table 3.** CAF zonation genes in the held-out Cervilla patient. Discovery direction (near versus distant stroma) and Cervilla Visium HD fibroblast pseudobulk log_2_FC (RNA mask *σ* 100 µm). The pre-registered test counts only the 22 genes with *≥*20 counts in both zones (21/22 agree); CP has no counts.

| Gene | Disc. | Cervilla | Gene | Disc. | Cervilla | Gene | Disc. | Cervilla |
|---|---|---|---|---|---|---|---|---|
| COL11A1 | up | 2.54 | THBS4 | down | −0.68 | LRRN4CL | down | −1.58 |
| MMP11 | up | 2.87 | SLIT3 | down | −2.56 | MEDAG | down | −0.87 |
| WNT2 | up | 2.04 | LAMA2 | down | −1.08 | CILP | down | −0.05 |
| INHBA | up | 1.32 | TCF21 | down | −0.97 | C16orf89 | down | −4.59 |
| WNT5A | up | 0.51 | ABCB5 | down | −2.03 | MFAP4 | down | −1.21 |
| TMEM158 | up | −0.18 | STEAP2 | down | −1.14 | CFD | down | −3.04 |
| AOX1 | down | −2.68 | TFPI2 | down | −1.02 | C3 | down | −1.19 |
| CCDC80 | down | −1.72 | SCARA5 | down | −1.42 | CPXM1 | down | −0.51 |
| CP | down | 0.00 | FAM110B | down | −1.11 | RIPOR3 | down | −0.29 |
| CHRD | down | −0.81 | OSR2 | down | −0.91 | GPC3 | down | −2.94 |
| ADH1B | down | −8.72 | SVEP1 | down | −0.72 |  |  |  |
| SFRP2 | down | −2.51 | MUSK | down | −0.52 |  |  |  |
| C7 | down | −4.34 |  |  |  |  |  |  |

**Supplementary Table 4.** Discovery hits of the state and niche transcriptomes. limma-voom on per-section pseudobulks (three Visium HD sections; spill-deconvolved counts). Fib., GREM1-type versus BMP-type fibroblasts (the 25 most negative and 25 most positive of 309 hits, ranked by *t*); Tum., GREM1-niche versus BMP-niche interior tumour cells (all 26 hits). 5K log_2_FC is shown for genes testable on the held-out Xenium 5K panel (n.t., not testable). Neither gene-level held-out test passed its pre-registered within-block null.

| Set | Gene | log<sub>2</sub>FC | *t* | *q* | 5K log<sub>2</sub>FC |
|---|---|---|---|---|---|
| Fib. | F3 | -4.53 | -23.0 | 5.1e-09 | n.t. |
| Fib. | TRPA1 | -5.49 | -21.2 | 6.5e-09 | +0.52 |
| Fib. | SPRY4 | -3.42 | -20.6 | 7.5e-09 | n.t. |
| Fib. | PCSK6 | -2.80 | -19.4 | 1.5e-08 | n.t. |
| Fib. | COL4A5 | -4.22 | -17.7 | 4.2e-08 | -3.92 |
| Fib. | WNT4 | -5.02 | -15.6 | 1.7e-07 | -3.88 |
| Fib. | SALL1 | -4.63 | -15.2 | 2.1e-07 | n.t. |
| Fib. | EMID1 | -3.19 | -15.0 | 2.1e-07 | n.t. |
| Fib. | PCDH18 | -1.87 | -15.0 | 2.1e-07 | -2.14 |
| Fib. | LRP4 | -2.74 | -14.7 | 2.3e-07 | n.t. |
| Fib. | NSG1 | -4.98 | -14.7 | 2.3e-07 | n.t. |
| Fib. | WNT5A | -2.14 | -13.8 | 4.1e-07 | -1.48 |
| Fib. | EDNRB | -1.94 | -13.7 | 4.1e-07 | -1.28 |
| Fib. | ADGRL3 | -4.04 | -13.4 | 4.9e-07 | n.t. |
| Fib. | PLAT | -2.01 | -13.4 | 5.0e-07 | -1.06 |
| Fib. | OBSCN | -2.48 | -12.6 | 9.2e-07 | n.t. |
| Fib. | PTCH1 | -1.90 | -12.6 | 9.2e-07 | -0.68 |
| Fib. | NRG1 | -4.73 | -12.6 | 9.2e-07 | n.t. |
| Fib. | ITGA8 | -2.00 | -12.4 | 1.1e-06 | n.t. |
| Fib. | BMP2 | -3.05 | -12.3 | 1.1e-06 | -1.94 |
| Fib. | SCUBE1 | -4.62 | -12.0 | 1.5e-06 | n.t. |
| Fib. | TBX2 | -2.11 | -11.9 | 1.6e-06 | -1.02 |
| Fib. | CXCL14 | -2.61 | -11.8 | 1.7e-06 | n.t. |
| Fib. | FOXF2 | -1.76 | -11.2 | 3.1e-06 | n.t. |
| Fib. | BMP8A | -1.90 | -11.1 | 3.6e-06 | -2.00 |
| Fib. | IGFBP4 | +1.41 | +9.5 | 1.5e-05 | n.t. |
| Fib. | IGF1 | +2.07 | +9.7 | 1.3e-05 | +1.04 |
| Fib. | NT5E | +2.01 | +9.8 | 1.2e-05 | +1.74 |
| Fib. | COL14A1 | +1.55 | +9.9 | 9.7e-06 | n.t. |
| Fib. | FAP | +1.88 | +10.0 | 9.7e-06 | +1.70 |
| Fib. | ADAM12 | +1.76 | +10.0 | 9.3e-06 | +2.39 |
| Fib. | CXCL12 | +1.57 | +10.1 | 8.5e-06 | +1.72 |
| Fib. | C7 | +1.82 | +10.3 | 7.4e-06 | n.t. |
| Fib. | C3 | +1.70 | +10.4 | 6.6e-06 | n.t. |
| Fib. | RSPO3 | +2.89 | +10.4 | 6.6e-06 | +2.85 |
| Fib. | ANGPTL2 | +1.43 | +10.5 | 6.0e-06 | +1.28 |
| Fib. | STEAP1 | +2.26 | +10.6 | 5.3e-06 | +2.39 |
| Fib. | PTGDS | +1.61 | +10.9 | 4.0e-06 | n.t. |
| Fib. | STEAP2 | +2.12 | +11.0 | 3.9e-06 | n.t. |
| Fib. | PTGIS | +1.90 | +11.0 | 3.8e-06 | +1.34 |
| Fib. | FGF7 | +1.88 | +11.1 | 3.7e-06 | +2.66 |
| Fib. | SRPX | +2.29 | +11.6 | 2.2e-06 | +1.37 |
| Fib. | CCDC80 | +2.16 | +12.2 | 1.3e-06 | +1.35 |
| Fib. | SFRP1 | +2.73 | +12.6 | 9.2e-07 | +1.62 |
| Fib. | PRELP | +2.14 | +13.2 | 5.6e-07 | n.t. |
| Fib. | MEDAG | +2.72 | +13.9 | 4.0e-07 | n.t. |
| Fib. | MGP | +2.55 | +14.2 | 3.2e-07 | n.t. |
| Fib. | OGN | +2.71 | +15.0 | 2.1e-07 | n.t. |
| Fib. | TNXB | +2.23 | +15.6 | 1.7e-07 | +1.13 |
| Fib. | SFRP2 | +3.02 | +17.0 | 6.2e-08 | n.t. |
| Tum. | ID1 | -2.27 | -12.3 | 2.2e-07 | n.t. |
| Tum. | SELENBP1 | -1.55 | -8.0 | 7.7e-05 | -0.99 |
| Tum. | LRATD1 | -1.15 | -8.0 | 7.7e-05 | n.t. |
| Tum. | HSD11B2 | -1.14 | -7.7 | 1.1e-04 | -0.54 |
| Tum. | MYBL2 | -1.17 | -7.3 | 2.0e-04 | +0.06 |
| Tum. | FABP1 | -1.41 | -7.0 | 3.2e-04 | n.t. |
| Tum. | HMGCS2 | -1.29 | -6.8 | 4.0e-04 | n.t. |
| Tum. | SLC26A3 | -1.43 | -5.3 | 5.9e-03 | -0.35 |
| Tum. | C10orf99 | -1.23 | -5.3 | 5.9e-03 | n.t. |
| Tum. | AGR3 | -1.01 | -5.1 | 8.5e-03 | -0.56 |
| Tum. | TM4SF20 | -1.63 | -4.8 | 1.4e-02 | n.t. |
| Tum. | KIFC1 | -1.23 | -4.7 | 1.5e-02 | n.t. |
| Tum. | SORD | -1.25 | -4.4 | 3.0e-02 | +0.09 |
| Tum. | CCNE2 | -1.34 | -4.3 | 3.2e-02 | n.t. |
| Tum. | SPTBN5 | +1.79 | +4.2 | 3.5e-02 | n.t. |
| Tum. | ANKRD18A | +1.34 | +4.3 | 3.2e-02 | n.t. |
| Tum. | GDPD5 | +1.28 | +4.5 | 2.3e-02 | n.t. |
| Tum. | PTPRN2 | +1.03 | +5.4 | 5.3e-03 | n.t. |
| Tum. | SLC2A12 | +1.22 | +5.4 | 4.6e-03 | n.t. |
| Tum. | PROM1 | +1.10 | +5.6 | 3.6e-03 | +1.46 |
| Tum. | LEFTY1 | +1.03 | +5.9 | 2.0e-03 | n.t. |
| Tum. | CEL | +1.43 | +6.0 | 1.7e-03 | n.t. |
| Tum. | MMP7 | +2.33 | +6.2 | 1.3e-03 | n.t. |
| Tum. | PMFBP1 | +1.34 | +6.3 | 1.2e-03 | +0.42 |
| Tum. | LAMC2 | +1.22 | +6.3 | 1.2e-03 | +0.17 |
| Tum. | PLEKHB1 | +1.67 | +7.1 | 2.8e-04 | n.t. |

**Supplementary Table 5.**
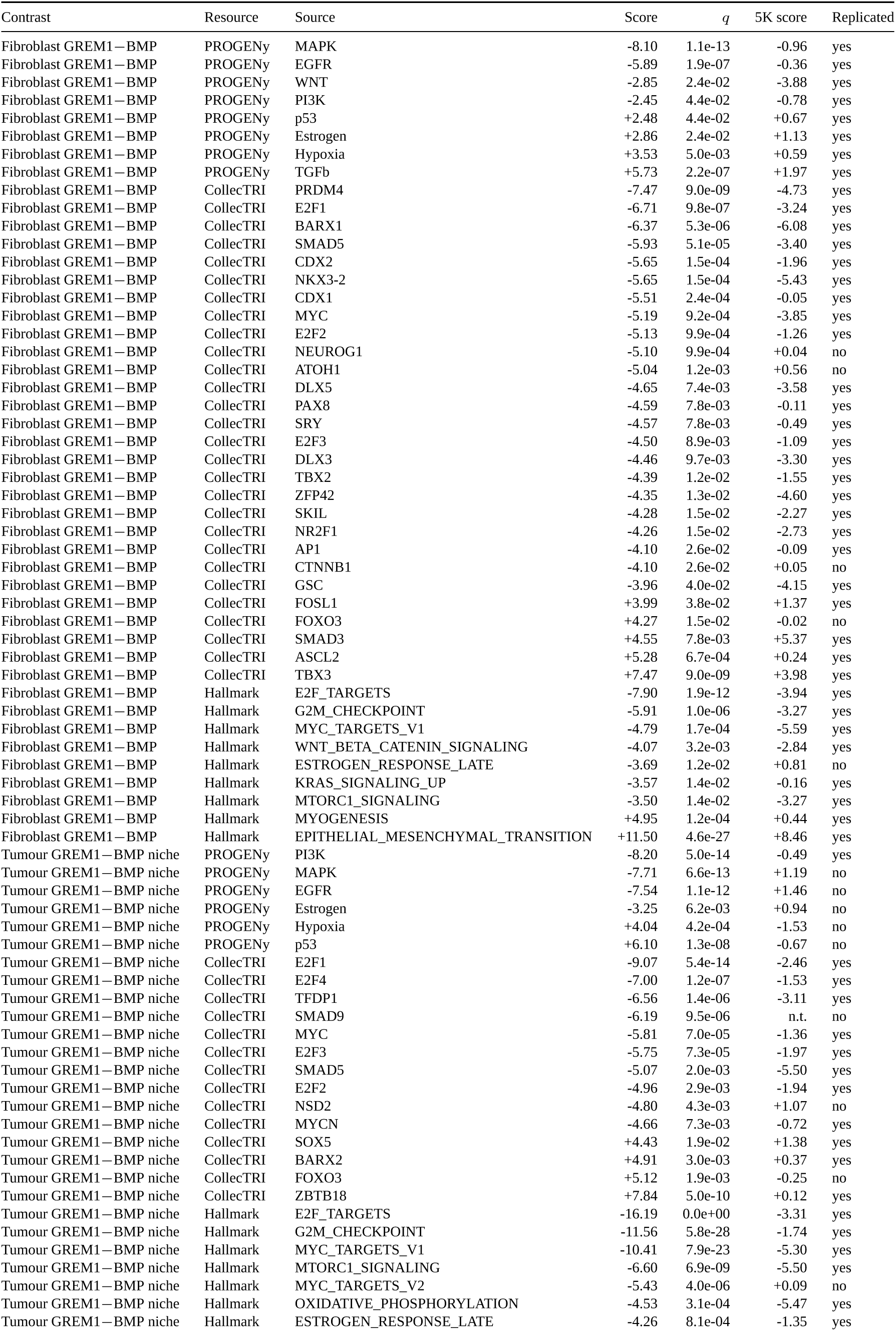
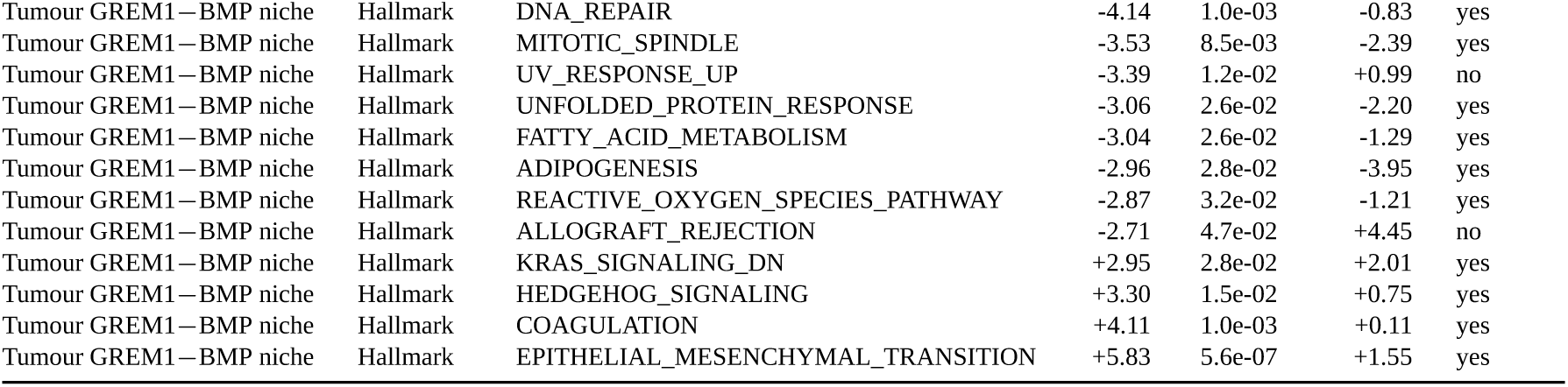
Pathway and transcription-factor activity of the fibroblast states and tumour niches. decoupler univariate linear model on the limma moderated *t* (discovery, three Visium HD sections) and on Welch *t* (held-out Xenium 5K, panel genes). Activities with discovery BH *q <* 0.05 are listed; “replicated” is the pre-stated same-sign rule (no spatial null at pathway level); n.t., regulator or set not testable on the 5K panel. Positive scores are higher in GREM1-type fibroblasts or in GREM1-niche tumour cells.

**Supplementary Table 6.**
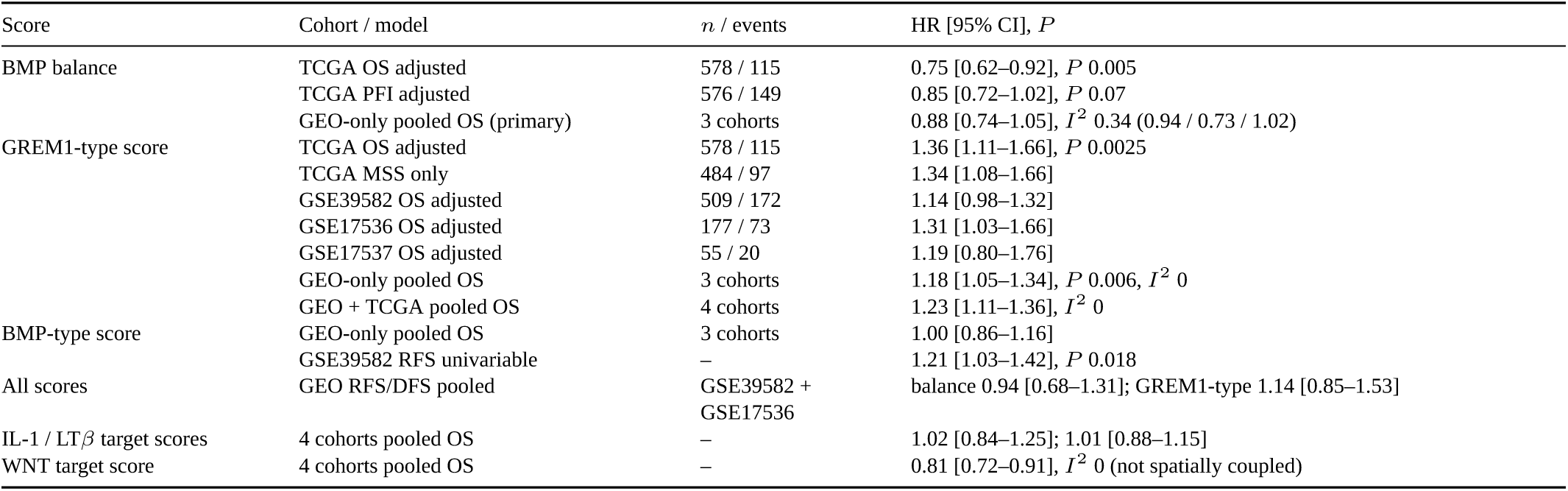
Bulk survival of the fibroblast BMP-axis scores. HR per s.d.; scores residualised on fibroblast abundance. TCGA adjusted for stage, age, MSI and site; GEO for stage and age (+ MMR in GSE39582). Pooling: DerSimonian–Laird on log HR, complete cases.

| Score | Cohort / model | *n* / events | HR [95% CI], *P* |
|---|---|---|---|
| BMP balance | TCGA OS adjusted | 578 / 115 | 0.75 [0.62–0.92], *P* 0.005 |
|  | TCGA PFI adjusted | 576 / 149 | 0.85 [0.72–1.02], *P* 0.07 |
|  | GEO-only pooled OS (primary) | 3 cohorts | 0.88 [0.74–1.05], *I*<sup>2</sup> 0.34 (0.94 / 0.73 / 1.02) |
| GREM1-type score | TCGA OS adjusted | 578 / 115 | 1.36 [1.11–1.66], *P* 0.0025 |
|  | TCGA MSS only | 484 / 97 | 1.34 [1.08–1.66] |
|  | GSE39582 OS adjusted | 509 / 172 | 1.14 [0.98–1.32] |
|  | GSE17536 OS adjusted | 177 / 73 | 1.31 [1.03–1.66] |
|  | GSE17537 OS adjusted | 55 / 20 | 1.19 [0.80–1.76] |
|  | GEO-only pooled OS | 3 cohorts | 1.18 [1.05–1.34], *P* 0.006, *I*<sup>2</sup> 0 |
|  | GEO + TCGA pooled OS | 4 cohorts | 1.23 [1.11–1.36], *I*<sup>2</sup> 0 |
| BMP-type score | GEO-only pooled OS | 3 cohorts | 1.00 [0.86–1.16] |
|  | GSE39582 RFS univariable | – | 1.21 [1.03–1.42], *P* 0.018 |
| All scores | GEO RFS/DFS pooled | GSE39582 + GSE17536 | balance 0.94 [0.68–1.31]; GREM1-type 1.14 [0.85–1.53] |
| IL-1 / LTβ target scores | 4 cohorts pooled OS | – | 1.02 [0.84–1.25]; 1.01 [0.88–1.15] |
| WNT target score | 4 cohorts pooled OS | – | 0.81 [0.72–0.91], *I*<sup>2</sup> 0 (not spatially coupled) |

**Supplementary Table 7.** Robustness of the GREM1-state score (pre-registered) OS; HR per s.d.; adjusted for stage and age (+ MSI and site in TCGA, + MMR in GSE39582) unless stated.

| Analysis | Cohort | *n* / events | HR [95% CI] or Δ*C* |
|---|---|---|---|
| All four cohorts, DL pooled | TCGA + 3 GEO | 1,319 / 380 | 1.23 [1.11–1.36], *I*<sup>2</sup> 0 |
| Leave-one-out: without TCGA | 3 GEO | 741 / 265 | 1.19 [1.05–1.34] |
| Without GSE39582 | 3 cohorts | 810 / 208 | 1.32 [1.14–1.52] |
| Without GSE17536 | 3 cohorts | 1,142 / 307 | 1.21 [1.08–1.36] |
| Without GSE17537 | 3 cohorts | 1,264 / 360 | 1.24 [1.11–1.38] |
| + CIT subtype, KRAS, BRAF, TP53, CIMP, adjuvant chemo | GSE39582 | 280 / 98 | 0.97 [0.78–1.21] |
| Same, without TP53 | GSE39582 | 420 / 135 | 1.04 [0.87–1.26] |
| + CIT subtype only | GSE39582 | 509 / 172 | 1.10 [0.93–1.30] |
| Harrell’s Δ*C* | TCGA; GSE39582;<br>GSE17536; GSE17537 | – | +0.022 [0.002, 0.052]; +0.005; +0.015; +0.009 |
| Pooled Δ*C* (DL) | 4 cohorts | 1,319 | +0.009 [−0.001, 0.019] |
| Score × adjuvant chemotherapy | GSE39582 stage II–III | 416 / 128 | interaction 1.79 [1.20–2.67] |
| Treated / untreated | GSE39582 stage II–III | 190 / 226 | 1.56 [1.11–2.18] / 0.84 [0.67–1.05] |

